# A consensus single-cell transcriptomic atlas of dermal fibroblast heterogeneity

**DOI:** 10.1101/2024.09.05.611379

**Authors:** Alex M. Ascensión, Ander Izeta

## Abstract

Single-cell RNA sequencing (scRNAseq) studies have unveiled large transcriptomic heterogeneity within both human and mouse dermal fibroblasts, but a consensus atlas that spans both species is lacking. Here, by studying 25 human and 9 mouse datasets through a semi-supervised procedure, we categorize 15 distinct human fibroblast populations across 5 main axes. Analysis of human fibroblast markers characteristic of each population suggested diverse functions, such as position-dependent ECM synthesis, association with immune responses or structural roles in skin appendages. Similarly, mouse fibroblasts were categorized into 17 populations across 5 axes. Comparison of mouse and human fibroblast populations highlighted similarities suggesting a degree of functional overlap, though nuanced differences were also noted: transcriptomically, human axes seem to segregate by function, while mouse axes seem to prioritize positional information over function. Importantly, addition of newer datasets did not significantly change the defined population structure. This study enhances our understanding of dermal fibroblast diversity, shedding light on species-specific distinctions as well as shared functionalities.

## INTRODUCTION

Dermal fibroblasts not only produce most of the extracellular matrix (ECM) in the skin, they are also a dynamic, heterogeneous and highly plastic cell type (Ahuja et al., 2022; Capolupo et al., 2022; Lendahl et al., 2022), which is key in specifying natural variations among the diverse skin areas and skin appendages (Myung et al., 2022; Jacob et al., 2023; Wiedemann et al., 2023; Lee et al., 2024). In fact, the true extent of fibroblast heterogeneity is most possibly beginning to unravel (Parker et al., 2023). In homeostasis, fibroblast subsets are also involved in relevant cutaneous functions such as immune regulation, cell motility and progenitor cell replenishment (Plikus et al., 2021). Of note, both positional information (Chitturi and Leask, 2024) and mechanical cues (Tan et al., 2022; Younesi et al., 2024) seem to be highly relevant in determining fibroblast population diversity. Furthermore, the key roles of dermal fibroblast subsets in establishing and maintaining pathological states such as chronic wounds, skin scarring/fibrosis and cutaneous autoimmune diseases are now well-established (Ascensión et al., 2022a; Ganier et al., 2022; Gur et al., 2022; Bensa et al., 2023; Knoedler et al., 2023; Shi et al., 2024; Zhu et al., 2024). Similarly, distinct fibroblast states seem to have a central role in orchestrating the dynamic phases of cutaneous wound repair (Correa-Gallegos et al., 2023; Almet et al., 2024).

Single-cell methods are very useful to unravel cell and tissue heterogeneity or to find details on how a physiological process develops (Yuan et al., 2024). In the last decade, single-cell RNA sequencing (scRNAseq) has been widely applied to decipher skin cell heterogeneity (Liu and Plikus, 2023), in some cases with large cell numbers sequenced and coupled with spatial transcriptomics or even parallel multiomics information (Reynolds et al., 2021; Thompson et al., 2022; Thrane et al., 2023; Ganier et al., 2024). However, there are issues arising from the analysis of some of the large datasets that need be taken into account (Ascensión et al., 2022b). Additionally, a lack of consensus across the definitions of fibroblast subsets in the different publications is still a major drawback in the field, because it largely hinders cross-laboratory replication and validation of original studies (Phan et al., 2021). A first attempt to unify the heterogeneity found in the literature classified human dermal fibroblasts into 10 populations distributed in 3 major axes (Ascensión et al., 2021). However, the consensus analysis was based in a reduced number of datasets and fibroblast populations were manually assigned. Furthermore, as of today there is no way to correlate human cell population data with those of the laboratory mouse, where most functional studies are performed.

For said reasons, we aimed to expand our initial study to increase the number of datasets analyzed and include mouse cells. Here, we explored the dermal fibroblast heterogeneity in a total of 25 human datasets using a semi-supervised population assignment algorithm. Additionally, we extended the analysis to mouse dermal fibroblasts by examining 9 published datasets. In total, >150,000 fibroblasts (108,440 human cells and 46,549 mouse cells) were included in our combined analysis. We analyzed the similarities between mouse and human populations, and inferred putative functions of each human fibroblast population by performing an extensive literature review of the most relevant markers. Addition of newer datasets did not increase the number or main characteristics of the newly defined fibroblast populations. Thus, our results provide a definitive consensus cell atlas of dermal fibroblast cell populations that will be instrumental in exploring the involvement of specific fibroblast subsets in cutaneous pathology, as well as injury repair.

## RESULTS

### Human dermal fibroblasts are divided into 5 main axes and 15 populations

Previous efforts to build a consensus human dermal fibroblast cell atlas were based on a limited number of datasets. To update our previous analysis, twenty-five publicly available human dermal fibroblast scRNAseq datasets encompassing diverse skin areas (Table S1) were downloaded and processed by *Uniform Manifold Approximation and Projection (UMAP)* and *Leiden* packages, as detailed in Materials & Methods. Within each dataset, fibroblasts were defined as cell populations expressing *COL1A1*, *DCN*, *LUM*, and *PDGFRA*. To minimize human selection bias, two algorithms (a marker-to-population algorithm and a population-to-marker algorithm, further described as Supplementary Materials & Methods) were developed and combined to determine consensus fibroblast populations. Axes (major fibroblast subtypes) and populations were named by expanding the notation that we used in a previous article (Ascensión et al., 2021). Combined analysis of the 25 human datasets (108,440 sequenced fibroblasts, isolated from 168 adult donors of both sexes and diverse ethnicity; age range 18-82 years) yielded a total of 15 populations divided in 5 axes that were present in at least 2/3 of the datasets, and which were named A to E. UMAP plots of human fibroblast populations across datasets (Figure S1) revealed a fully consistent (25 of 25; 100%) pattern of division in two main axes: axis A and axis B; with an additional axis C present in most datasets (20 of 25; 80%). Two additional axes, D and E, were less obvious, having consistently fewer cells: D was detected in 18/25 (72%) and E in 17/25 (68%) datasets, respectively. Lastly, two other potential axes were considered unreliable because they were detected in 13/25 (52%) datasets (axis T, for *transitional*) and in 7/25 (28%) datasets (axis U, for *unknown*). A selection of four marker genes whose overexpression defines each fibroblast population is shown in Figure 1a and Table 1.

**Figure 1.**
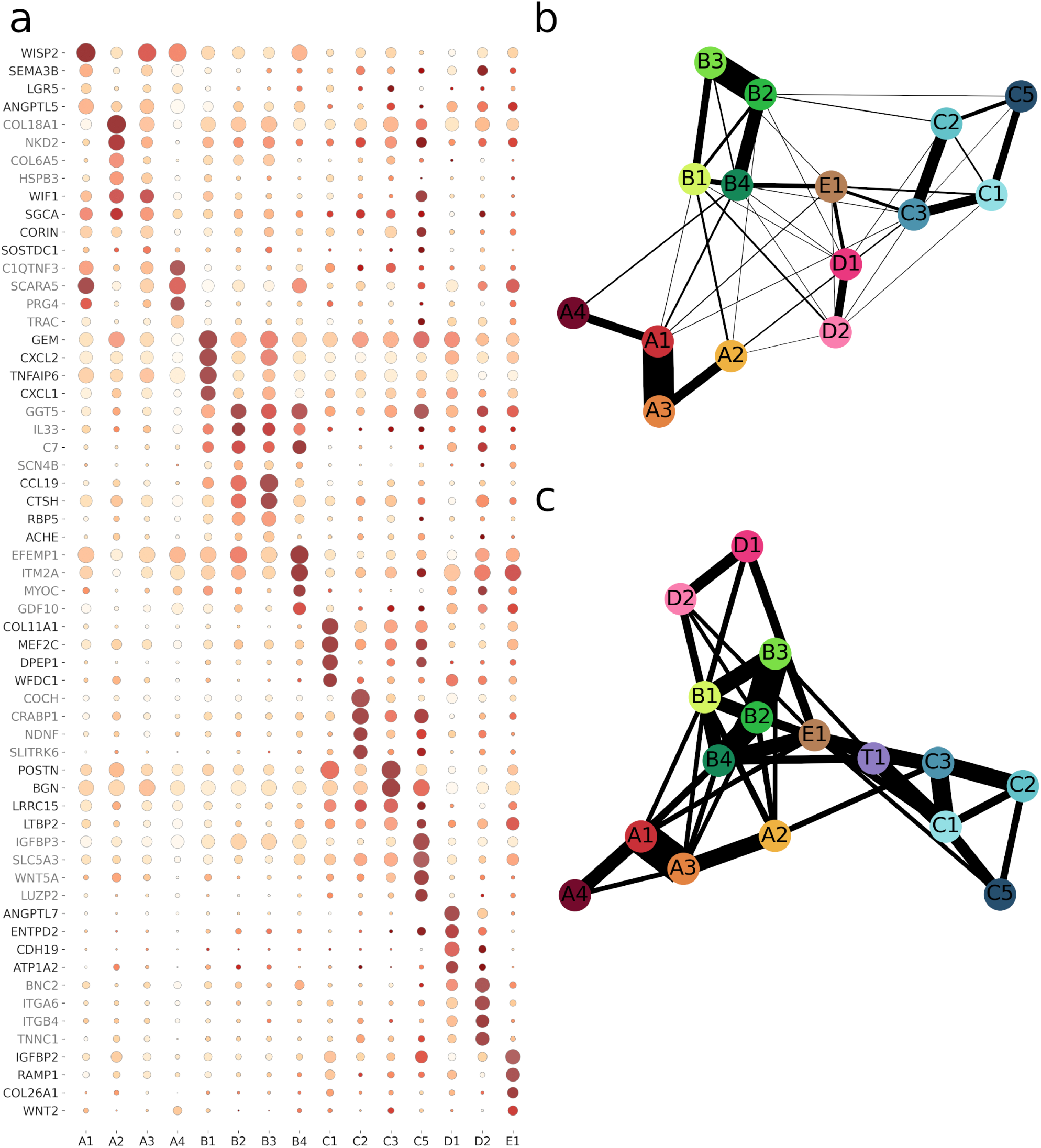
Summary of human dermal fibroblast populations. (a) Dot-plot with human dermal fibroblast markers. 4 markers are chosen for each population. The size of the circle represents the proportion of cells in that population expressing the marker–the larger the circle the larger the proportion–; and the colour represents the mean expression of the gene in that population–the redder/browner the colour the higher the expression–. (b) and (c) PAGA graphs showing transcriptomic relationships between populations. Graph (B) is constructed from merging all PAGA trees (Figure S2), and graph (c) is constructed from merging all PAGA graphs (Figure S3).

**Table 1.**
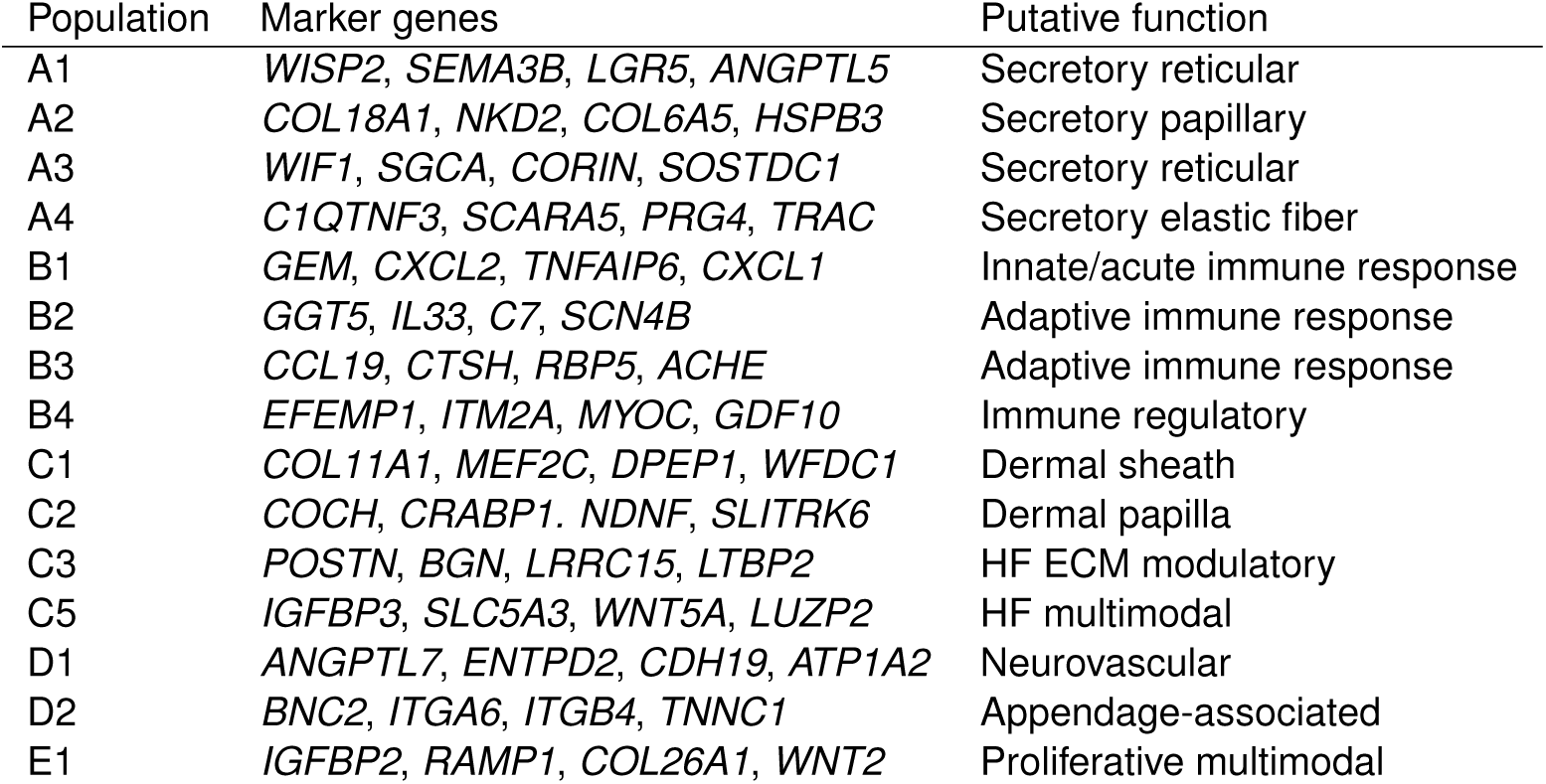
Summary table of human dermal populations. Each population is associated with its most representative four markers. Putative functions are assigned based on the bibliography analysis described in the Discussion section and in Supplementary Discussion.

| Population | Marker genes | Putative function |
| --- | --- | --- |
| A1 | *WISP2, SEMA3B, LGR5, ANGPTL5* | Secretory reticular |
| A2 | *COL18A1, NKD2, COL6A5, HSPB3* | Secretory papillary |
| A3 | *WIF1, SGCA, CORIN, SOSTDC1* | Secretory reticular |
| A4 | *C1QTNF3, SCARA5, PRG4, TRAC* | Secretory elastic fiber |
| B1 | *GEM, CXCL2, TNFAIP6, CXCL1* | Innate/acute immune response |
| B2 | *GGT5, IL33, C7, SCN4B* | Adaptive immune response |
| B3 | *CCL19, CTSH, RBP5, ACHE* | Adaptive immune response |
| B4 | *EFEMP1, ITM2A, MYOC, GDF10* | Immune regulatory |
| C1 | *COL11A1, MEF2C, DPEP1, WFDC1* | Dermal sheath |
| C2 | *COCH, CRABP1, NDNF, SLITRK6* | Dermal papilla |
| C3 | *POSTN, BGN, LRRC15, LTBP2* | HF ECM modulatory |
| C5 | *IGFBP3, SLC5A3, WNT5A, LUZP2* | HF multimodal |
| D1 | *ANGPTL7, ENTPD2, CDH19, ATP1A2* | Neurovascular |
| D2 | *BNC2, ITGA6, ITGB4, TNNC1* | Appendage-associated |
| E1 | *IGFBP2, RAMP1, COL26A1, WNT2* | Proliferative multimodal |

Axis A comprises fibroblasts seemingly involved in extracellular matrix (ECM) secretion, and is divided into four main populations: A1 to A4 (whenever possible, originally assigned population names were kept). Population A1 is defined by the expression of *WISP2*, *SEMA3B*, *LGR5* and *ANGPTL5*; A2 by *COL18A1*, *NKD2*, *COL6A5* and *HSPB3*; A3 by *WIF1*, *SGCA*, *CORIN* and *SOSTDC1*; and A4 by *C1QNTF3*, *SCARA5*, *PRG4*, and *TRAC* (Figure 1a). Potential function(s) of each of the populations are discussed below and in Supplementary text. Based on UMAP populations, all four A axis populations seemed to form a continuum. This was confirmed by individual *Partition-based graph abstraction (PAGA)* graphs showing transcriptomic relationships between populations within each dataset. *PAGA trees* and full *PAGA graphs* of dermal fibroblast populations are shown in Figure S2 and Figure S3, respectively. A consensus combination of all trees and graphs (Figures 1b, and 1c, respectively) confirmed a generalized close relationship among A populations. A1 and A2 are originally the most prominent and archetypal populations, whereas A3 is a bridge population between A1 and A2. Lastly, A4 is a population close to A1–most A4 and A1 markers are cross-expressed–, but its own marker expression–e.g. *PRG4* or *TRAC*–separates it from A1. Based on UMAP plots and PAGA graphs, A4 might be a *differentiated* state, or a subtype, of A1.

Axis B comprises fibroblasts involved in cutaneous immune responses and is divided into 4 populations: B1 to B4. Population B1 is defined by the expression of *GEM*, *CXCL2*, *TNFAIP6* and *CXCL1*; B2 by *GGT5*, *IL33*, *C7* and *SCN4B*; B3 by *CCL19*, *CTSH*, *RBP5* and *ACHE* ; and B4 by *EFEMP1*, *ITM2A*, *MYOC*, and *GDF10*. Similar to axis A, populations B1 and B2 are “archetypal”, and B3 appears to be a bridge population, which expresses several “canonical” B2 markers, such as *CCL19* or *CD74*; as well as B1 markers such as *CXCL2* and *CXCL3*. Relationships between B populations in the PAGA graphs are not as clear as with axis A: B1 seems to be partially independent from the rest of populations, although connections between B1, B2 and B4 are observable.

Axis C encompasses more specialized fibroblast subsets and is divided into 4 populations: C1, C2, C3, and C5. Of note, we skipped the C4 denomination to avoid confusion with the C4 population from Ascensión et al. (2021), which is now reclassified as the novel axis D (see below). C1 is defined by the expression of *COL11A1*, *MEF2C*, *DPEP1*, and *WFDC1*; C2 by *COCH*, *CRABP1*, *NDNF*, and *SLITRK6*; C3 by *POSTN*, *BGN*, *LRRC15*, and *LTBP2*; and C5 by *IGFBP3*, *SLC5A3*, *WNT5A* and *LUZP2*. C1 and C2 have their own transcriptomic profile, but C3 does not–with the slight exception of *POSTN*, which is more expressed in C3 than in C1 or C2–, and is instead defined as a “less distinctive” population due to its reduced number of markers, most of which are partially shared with C1 or C2. Population C5, when present, appears to branch from C1, C2 or C3 without a clear pattern.

Populations D1 and D2 of axis D are derived from the original C4 population from Ascensión et al. (2021). D1 population is defined by the expression of *ANGPTL7*, *ENTPD2*, *CDH19* and *ATP1A2*; and D2 population is defined by the expression of *BNC2*, *ITGA6*, *ITGB4* and *TNNC1*. Although these two populations are different, most characteristic population markers are co-expressed by both D1 and D2. Looking at the PAGA graphs, the joint graphs show that D1 and D2 populations are quite independent of the rest, although they may have a certain degree of connection with B axis populations (Figure 1b).

Lastly, E1 is a new population not assigned to any of the previous axes, and shows a specific transcriptomic profile, defined by the expression of *IGFBP2*, *RAMP1*, *COL26A1* and *WNT2*. Looking at the UMAPs, it seems likely that this population acts as a bridge or a transitional fibroblast status between axes B, C and D.

Addition of the few last datasets did not significantly change the axis/population structure. Based on this analysis, and due to the large number of datasets from human dermal fibroblasts included, as well as the heterogeneity of sampling sites, it seems unlikely that new fibroblast populations will emerge in case new datasets are integrated, and thus virtually all human dermal fibroblast heterogeneity is captured in this work.

### Murine dermal fibroblasts are divided into 5 main axes and 17 populations

Human and mouse skin present numerous structural and functional differences (Lynch and Watt, 2018). Notwithstanding, a significant proportion of functional dermatological research is performed in genetically modified mouse models and thus correlative data of mouse and human cell populations are of great interest to understand their potential role(s) in human pathology. To initiate efforts to provide a consensus mouse dermal cell atlas, nine dermal fibroblast scRNAseq datasets of C57BL/6J and BALB/c mouse strains (Table S2) were downloaded and processed as detailed in the previous section. To avoid confusion with the human axes and populations, and before potential similarities among species could be inferred, the mouse fibroblast axes were named in small case and using the last letters of the alphabet. Combined analysis of the 9 mouse datasets (46,549 sequenced fibroblasts, isolated from 25 mice of both sexes; age range 3 weeks to 18 months) yielded 17 fibroblast populations in 5 major axes (v to z) and 2 additional axes that were considered *bridge* or *transitional* between two of them (w/x and x/y). A UMAP representation of all datasets is available in Figure S4.

Axis x populations presented two constitutive populations (x1 and x2) and two transitional populations (w/x and x/y), defined as follows (Figure 2a and Table 2): population x1 is defined by the expression of *Fgfr4*, *Gpha2*, *Cib3* and *Serpina3m*; population x2 is defined by the expression of *Igfbp2*, *Stc1*, *Sema3a* and *Enho*; population w/x is defined by the expression of *Coch*, *Emid1*, *Kera* and *Ntn5*; and population x/y is defined by the expression of *Crp*, *Akr1cl*, *Lgr5* and *Mup20*. Individual PAGA trees (Figure S5), and PAGA graphs of the mouse populations in each dataset (Figure S6) were combined to present a consensus picture (Figure 2b-c). From the PAGA graphs, x2 seems to be a terminal state of x1; that is, x1 and x2 are consistently related, and x2 is in terminal points of UMAPs and individual tree graphs. Although x2 is related to other populations, x1 shows stronger relationships, especially with x/y and w/x. Axis y presented five distinct populations, defined as follows: population y1 is defined by the expression of *Postn*, *Fabp4*, *Cd36* and *Pparg*; population y2 is defined by the expression of *Hmcn2*, *Col6a6*, *Fbln7* and *Bmp5*; population y3 is defined by the expression of *Ccn5*, *Ccn2*, *Ecrg4* and *Fgf9*; population y4 is defined by the expression of *C2*, *C4b*, *Chrdl1* and *Gdf10*; population y5 is defined by the expression of *Vwa1*, *Vit*, *P2ry14* and *Kcnk2*. The net of interactions within and with other axes is complex, almost behaving like a clique (Figure 2b-c). Based on the joint PAGA graphs, individual PAGA graphs, and UMAP plots, y2 seems to be a nexus of interactions between y3, y4, and, to a lesser extent, y1; and y1 also interacts with y4. y5 is closer in interaction to y4. Therefore, axis y is highly connected, probably due to the large number of populations within it.

**Figure 2.**
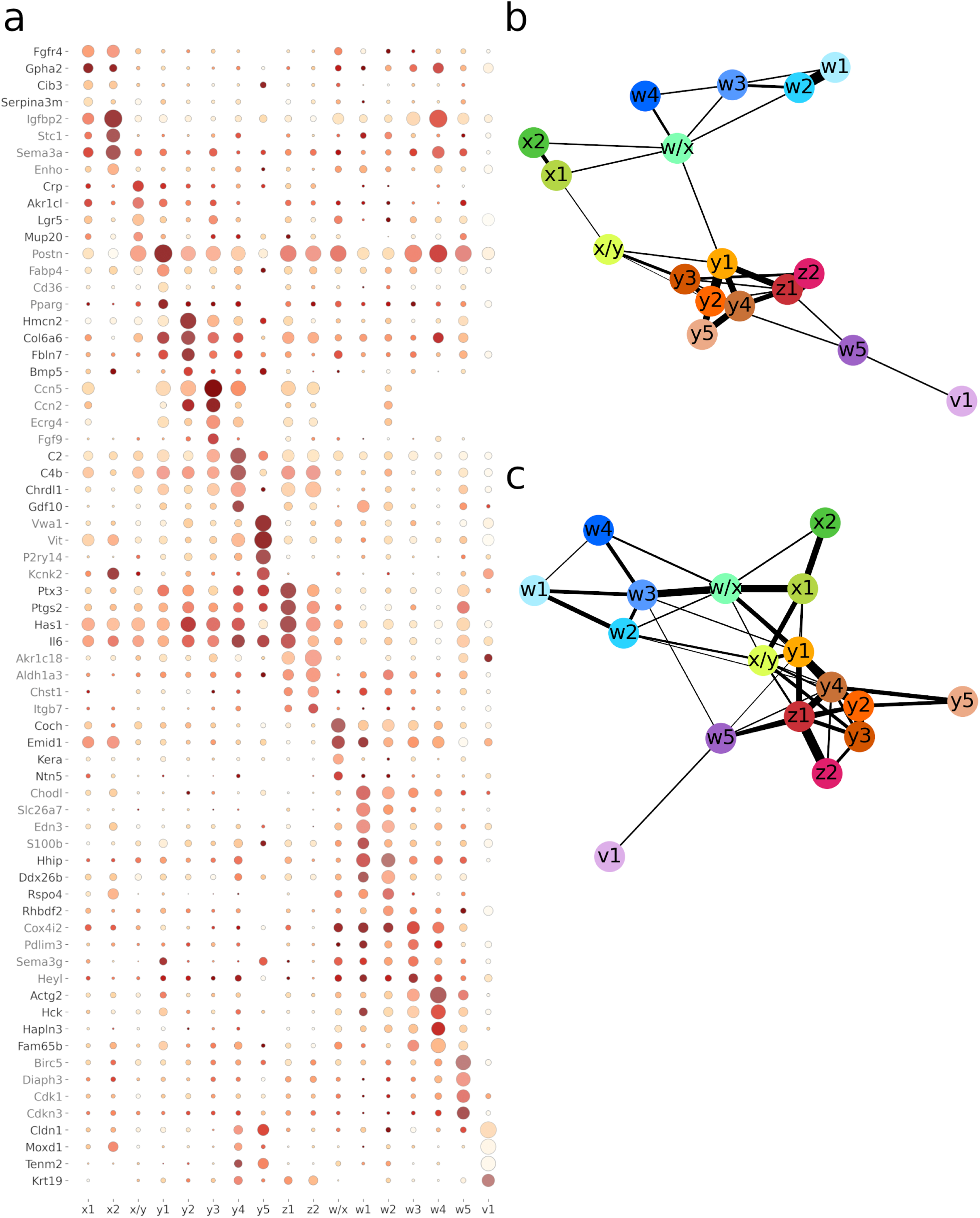
Summary of mouse dermal fibroblast populations. (a) Dot-plot with mouse dermal fibroblast markers. 4 markers are chosen for each population. The size of the circle represents the proportion of cells in that population expressing the marker–the larger the circle the larger the proportion–; and the colour represents the mean expression of the gene in that population–the redder/browner the colour the higher the expression–. (b) and (c) PAGA graphs showing transcriptomic relationships between populations. Graph (B) is constructed from merging all PAGA trees (Figure S5), and graph (c) is constructed from merging all PAGA graphs (Figure S6).

**Table 2.**
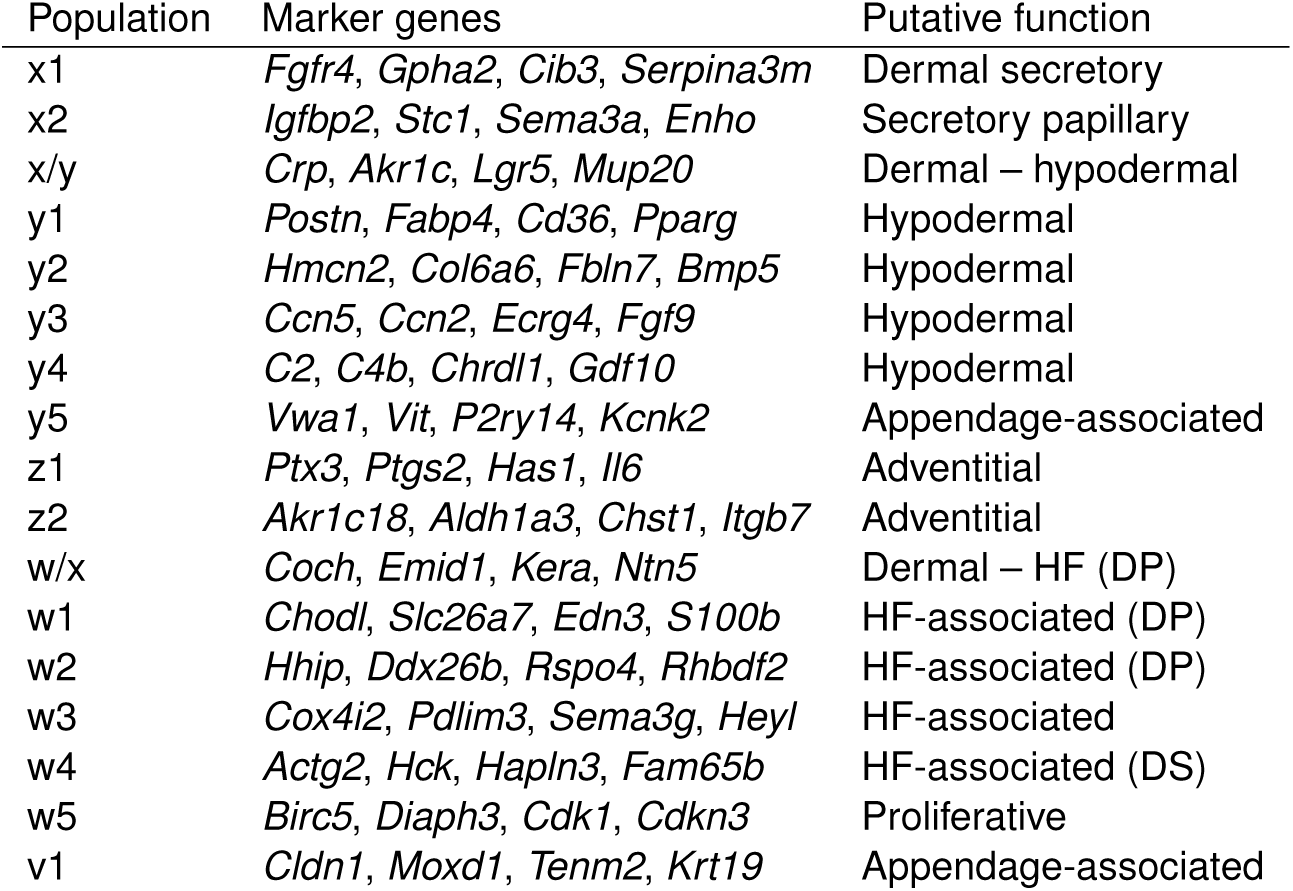
Summary table of mouse dermal populations. Each population is associated with its most representative four markers. Putative functions are assigned mainly based marker similarity with human populations and based on the locations described by Joost et al. (2020).

Axis z presented two independent populations: population z1 is defined by the expression of *Ptx3*, *Ptgs2*, *Has1* and *Il16*; and population z2 is defined by the expression of *Akr1c18*, *Aldh1a3*, *Chst1* and *Itgb7*. There is some variation between datasets regarding the topology and relationships between axis z populations. In the UMAPs of Abbasi et al. (2020); Shook et al. (2020); Vorstandlechner et al. (2020) datasets, we see that these two clusters are independent, whereas in the rest of datasets–Boothby et al. (2021); Buechler et al. (2021); Haensel et al. (2020); Joost et al. (2020)–axis z is integrated next to axis y and tends to be close to populations y2 and y3. The joint tree PAGA graph in Figure 2b–and individual PAGA graphs from some datasets show that z1 and z2 share a high resemblance and that axis z links to axis y mainly via the interaction between z1 and y1 or y4. Based on the unsupervised axis assignation, z1 and z2 are mainly assigned to the z axis, although the z1 population is sometimes assigned to the y axis, maybe due to the z1-y1 interaction described before. This discrepancy between z being a separate entity in certain datasets and being integrated within y axis in others is intriguing. One hypothesis is that the discrepancy presents an underlying biological basis (see below).

Axis w presented five fibroblast populations, defined as follows: population w1 is defined by the expression of *Chodl*, *Slc26a7*, *Edn3* and *S100b*; population w2 is defined by the expression of *Hhip*, *Ddx26b*, *Rspo4* and *Rhbdf2*; population w3 is defined by the expression of *Cox4i2*, *Pdlim3*, *Sema3g* and *Heyl*; population w4 is defined by the expression of *Actg2*, *Hck*, *Hapln3* and *Fam65b*; and population w5 is defined by the expression of *Birc5*, *Diaph3*, *Cdk1* and *Cdkn3*. Regarding the relationships between populations, it is clear that w1 and w2 populations share similar transcriptomic profiles, based on either the UMAPs or PAGA graphs. Additionally, w3 and w4 are also connected, although w3 seems connected to w1/w2 and w/x, as observed in the condensed graph from Figure 2b. Finally, the w5 population shows an inconsistent pattern related to axes z, w and v in the PAGA graphs and UMAP plots.

Lastly, population v1 is defined by the expression of *Cldn1*, *Moxd1*, *Tenm2* and *Krt19*. It seems to interact with w3, w5 and y5; however, these interactions are not fully consistent due to v1 population’s presence in only two datasets. In fact, in the combined PAGA graphs v1 interacts only with w5.

Based on the mouse dermal fibroblasts characterization published by Joost et al. (2020), our preliminary analyses led us to propose that axis x matches the expression profile of Joost’s FIB1 and FIB2 fibroblasts–*Col1a1*, *Sparc*, *Dcn*–, axis y matches the expression profile of FIB3–*Cxcl12*, *Gpx3*–, axis x/y is a mixture of markers from FIB1/2 and FIB3, and axis z matches the expression of FIB4–*Mfap5*, *Plac8* (Figure S7). Populations v and w are not directly associated with this layering, although axis w maps to DS and DP cells. Lastly, population y5 is possibly associated with FIB2 (further expanded in the Discussion section). Considering that the z axis may represent fibroblasts surrounding the *panniculus carnosus* and that the y axis may represent hypodermal fibroblasts, the close relationship between these two axes may indicate that they come in close contact and may interact with each other in mouse skin.

### Semi-supervised classification algorithm iterations demonstrate fibroblast population robustness

We next addressed the question of how robustly the assigned populations were maintained with algorithm iterations. To test the robustness of the marker-to-population algorithm, we applied a methodology similar to the classical *Jackknife* resampling method (Quenouille, 1956). For a fixed number of iterations–30 in this analysis–, and each dataset, we sampled 99% of the cells and re-ran the marker-to-population algorithm. The sampling was stratified across populations so that all populations were consistently represented.

Of note, a wide range of variation across datasets was detected, indicating that the robustness of the clustering is dataset-dependent (Figure 3a). For instance, data from Hughes et al. (2020), Reynolds et al. (2021), and Vorstandlechner et al. (2020) datasets presented lower mean robustness scores, with high interquartile ranges (statistical dispersion). However, since the sampling percentage is high, we should expect that the population assignment is robust and that the robustness score values of each fibroblast population are high. Figure 3b shows the distribution of the mean robustness score of each population across datasets. In human datasets, we observe a variation in scores between 0.6 and 0.9, with a mean value of about 0.75, which is acceptable considering the diversity of datasets and how their scores may be affected, as seen in Figure 3a. However, some populations, such as B3, B4 or D1 fibroblasts, showed lower robustness values. These observations were confirmed in violin plots of the distribution of scores per population and dataset (Figure S8). For instance, robustness scores of population B3 were high in some datasets–Liu et al. (2021); Rindler et al. (2021); Solé-Boldo et al. (2020); Theocharidis et al. (2022)–and extremely low in other datasets–Ahlers et al. (2022); Burja et al. (2022); Reynolds et al. (2021); Tabib et al. (2021). This comes in agreement with the previously described separation between B3, B2 and B1 populations. Interestingly, datasets that correctly discriminate B populations show high robustness scores, whereas, in the datasets with very low scores, the location of B3 in the UMAP is far more diffuse, or it represents a very small cluster (Figure S1), showcasing the *bridge* nature of this cluster.

**Figure 3.**
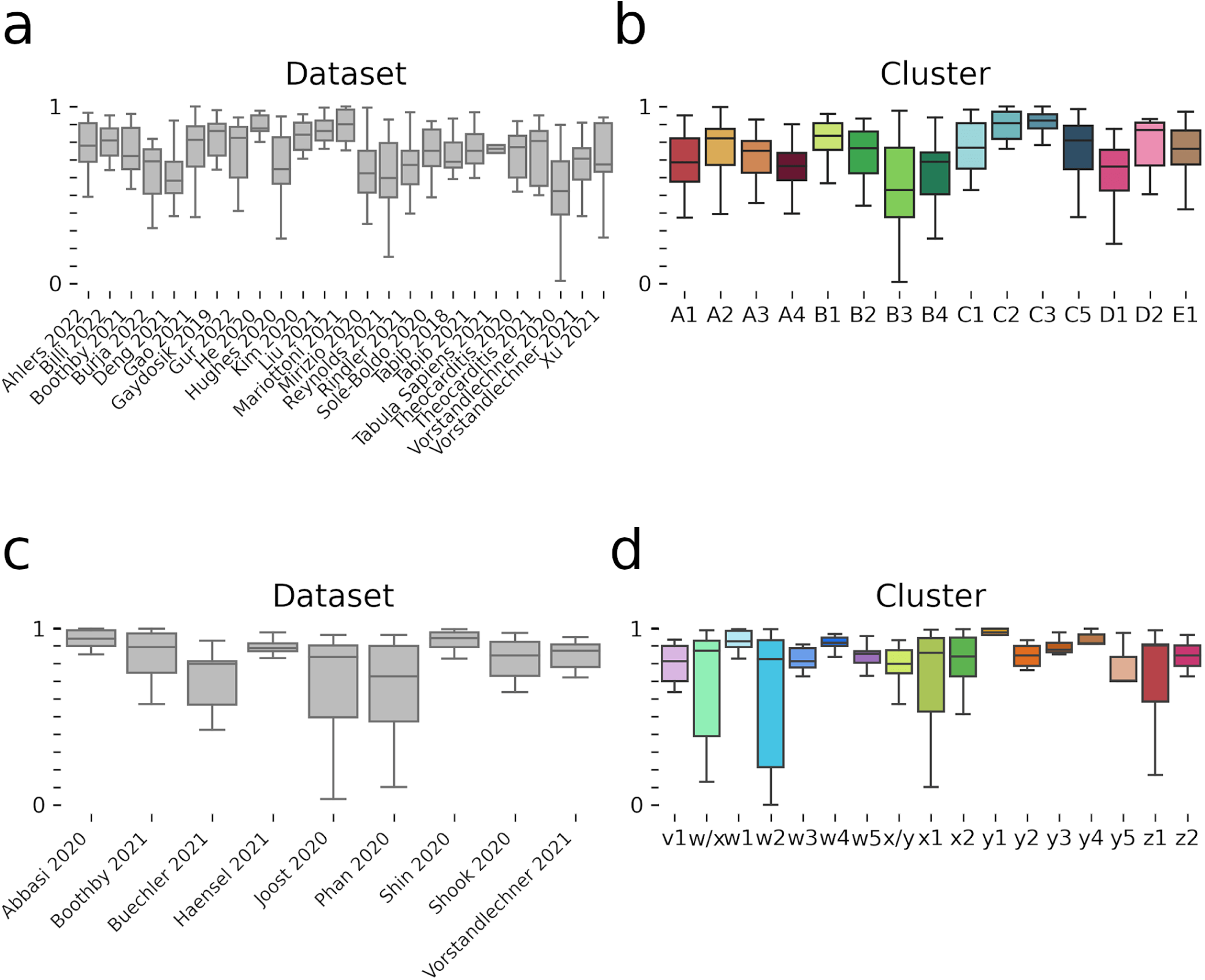
Boxplots of general robustness score. Boxplots in (a) and (c) represent the mean robustness score of each dataset, across populations in human (a) and mouse (c); whereas (b) and (d) represent the mean robustness score of each population, across datasets in human (b) and mouse (d). Each box represents the quartiles of the distribution, the centre bar represents the median, and whiskers extend to Q1 - 1.5 * IQR and Q3 + 1.5 * IQR.

In the case of mouse fibroblast populations, we observed generally higher robustness score values, reaching values >0.85 (Figure 3c). Despite that, there were some exceptions of populations with lower score values or wider dispersions: w/x, w2, x1 and z1. The distribution of scores per population and dataset (Figure S9) confirmed that the general robustness scores for mouse fibroblast populations were high, with some exceptions. For instance, w/x score was low in Joost et al. (2020) and variable in Shook et al. (2020); and z1 score was variable in Phan et al. (2020). In fact, most of this variability comes from the Joost et al. (2020) and Phan et al. (2020) datasets, as we observe in Figure 3c. This variability may indicate higher transcriptomic noisiness, possibly related to sample processing in those scRNAseq experiments (Ascensión et al., 2022b).

In conclusion, a certain degree of inter-dataset variability seems to explain the differences in robustness scores across datasets and fibroblast populations. Robustness scores were overall high enough to justify the assignment of fibroblasts to populations.

#### Non-robust fibroblast populations fuse onto similar populations

Classification algorithms assign the markers of a given cell to the most similar fibroblast population. However, if two populations are highly similar, it is likely that the algorithm fails to assign these populations separately, and this would justify a lack of robustness of the achieved clustering solution. To analyze the fate of non-robust populations in a given dataset, we calculated the adjacency matrix of the resampling results per dataset, as shown in Figures S10 (for human) and S11 (for mouse). The matrix indicates the proportion of assignments for each cell type (row) to the rest of the populations (columns). The median value across datasets was calculated to produce the combined adjacency matrix in Figure S12.

In human datasets, there was a generally high proportion of cases where the assigned cluster was the same as the original one, with some examples such as B1, C2, C5, D1 or E5, with values higher than 0.92. On the other hand, there were other populations, such as A1, A4 or B3, which had lower values–0.73, 0.69 and 0.59, respectively–. When looking at the assigned clusters, we observed that for A1, the most assigned cluster was A3–21% of the times–, for A4 it was A1–21% of the times– and for B3 they were B2 and B1–25% and 16% of the times, respectively–. This pattern is kept for individual datasets (Figure S10). For instance, for datasets which had lower proportions of A1 population assigned to A1, the second most assigned population was A3 (Dataset: proportion originally assigned/proportion of secondary assignation)–Ahlers et al. (2022): 0.49/0.20, Gur et al. (2022): 0.58/0.30, Reynolds et al. (2021): 0.47/0.47, Rindler et al. (2021): 0.56/0.38, Tabib et al. (2018): 0.60/0.32, Tabib et al. (2021): 0.38/0.56, Vorstandlechner et al. (2020): 0.58/0.35, Vorstandlechner et al. (2021): 0.59/0.35–. In some datasets, the second most assigned population was not A3–Kim et al. (2020): 0.45/B4 (0.25), Mirizio 2020: 0.50/A4 (0.21)–. This pattern was repeated for A4 and B3 as well.

For the rest of the populations, we observed similar assignment patterns. For instance, B4 population (0.75) was assigned similarly to B1 and B2 (0.12 and 0.09); D2 (0.89) was assigned to D1 (0.03) but also to B4 (0.06)–as observed in Figure S12–.

Regarding mouse populations, the assignment rate to the original population was generally higher (>0.85) as compared to human fibroblasts, but the assignment to other populations followed similar patterns. For instance, w/x (0.87) was primarily assigned to x1 (0.06); w2 (0.88) was assigned secondarily to w1 (0.12); w4 (0.94) was assigned to w3 (0.06); y3 (0.94) was assigned to y2 (0.04); y2 (0.86) was mainly assigned to y3 (0.09); and z1 and z2 (0.89, 0.92) were assigned to z2 and z1, respectively (0.06, 0.08).

Therefore, the population assignment algorithm showed good robustness levels in human datasets and very good robustness levels in mouse datasets. Moreover, the cases of lower population robustness were attributed to dataset-specific reasons, and therefore with the future addition of new high-quality datasets, population robustness should also increase.

### Comparison between mouse and human populations

As mentioned, the analysis of the transcriptomic similarity among mouse and human fibroblast populations would help clarify functionally relevant similarities and differences in the skin structure and function between the two organisms. To perform that comparison, we used two methods. First, we merged the samples from the two organisms in those datasets that contained information from both human and mouse cells, i.e. Boothby et al. (2021) and Vorstandlechner et al. (2021). Second, we established an overlap of populations based on the transcriptomic profiles of all datasets, assuming that populations with similar functions should have a higher degree of overlap in relevant markers than populations with dissimilar functions.

#### Traditional batch effect correction methods fail to integrate murine and human datasets on a large subset of genes

When merging the populations from Boothby et al. (2021) and Vorstandlechner et al. (2021), we applied an ortholog mapping scheme between mouse and human genes, and a common processing scheme for each merged dataset. Then, we applied a batch-effect correction by setting the organism as a batch to merge the datasets from both organisms. The results of the batch-effect correction method are shown in Figures S13A and S14A for Boothby et al. (2021) and Vorstandlechner et al. (2021) datasets, respectively. The UMAP plots show that the correction method failed to integrate the organism variable, since human and mouse cells were separated from each other.

Apparently, some underlying transcriptomic variability had to be causing this lack of integration between human and mouse populations. In fact, we observed a clear transcriptomic bias on the list of DEGs between human and mouse cells in both datasets (Figures S13B and S14B). Interestingly, these biases were partly dataset-specific: the biggest changes were due to differential expression of *RPL* and *RPS*, *B2M*, and *MALAT1* genes, but specific DEGs were different across organisms and datasets. Additionally, in the human dataset of Boothby et al. (2021), we detected the overexpression of genes associated with a stress response such as *SOD2* or *IER3*, and we also observed an increase of some fibroblast-associated markers such as *Col3a1*, *Col1a2* or *Bgn* in the mouse dataset, which have been previously reported as a possible false-positive signal due to contamination of samples (Vorstandlechner et al. (2021); Rojahn et al. (2020)), that is, markers that are overrepresented in the samples due to the presence of ambient RNA derived from damaged cells during library preparation that can then contaminate other cell types (Caglayan et al., 2022).

In an attempt to correct unwanted sources of variation from *scanpy*, we applied a linear regression method (sc.pp.regress_out). Although the cells from human and mouse organisms were nearer to each other in the UMAP plot, they did not have enough overlap to do the analysis (Figures S13C and S14C). Therefore, we abandoned this approach because applying further processing steps to merge the samples may introduce artifacts into the dataset due to overcorrection (Tran et al., 2020), and because it implicitly assumes identical cell type composition across batches (Haghverdi et al., 2018; Andrews, 2020).

#### Human-mouse marker-based comparison

As a second strategy to compare the human and mouse fibroblast populations, we extracted the most significant 250 markers for each organism and fibroblast population, and computed the Jaccard index for all pairings (Figure 4). Relevant overlapping values ranged between 0.08 to 0.14, similar to those obtained in human-human and mouse-mouse population comparisons (see Supplementary Information). Tables with the top 250 markers for each organism are available in the Supplementary Information (*robust_markers_human.xlsx* and *robust_markers_mouse.xlsx*).

**Figure 4.**
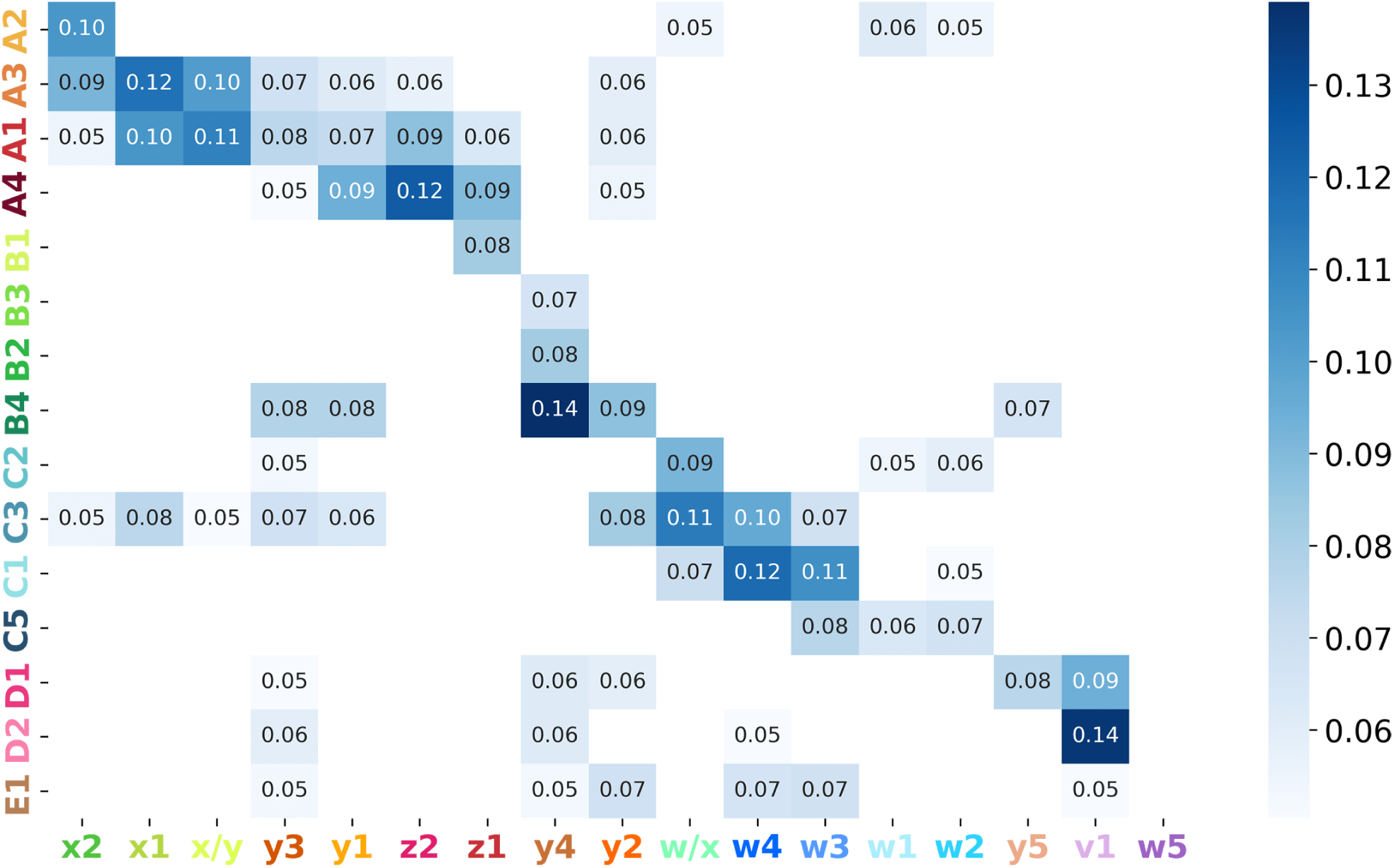
Heatmap of human and mouse marker overlap across populations. For each pair of populations, the number in the cell represents the proportion of overlapping markers (N=150).

Human axis A presented an overlap of its component populations with axes x, y and z in the mouse, indicating that the characteristics and/or functionality of axis A might be more distributed in mice. For instance, the A1 population shared similarities with x1, x/y, y3 and z2, most prominently with x1, which shared 6 genes (*CGREF1*, *CPZ*, *CYBRD1*, *CYP4B1*, *MMP27*, *SEMA3B*) and with z2, which shared 12 genes (*ACKR3*, *ADGRD1*, *CD55*, *DPP4*, *ISLR*, *LIMS2*, *MTCL1*, *NPR1*, *PI16*, *PRKG2*, *SEMA3E*, *TUBB4A*). The A2 population shared 8 relevant markers with the x2 population (*APELA*, *COL13A1*, *COL23A1*, *DAAM2*, *F5*, *NKD2*, *PTK7*, *RSPO3*) and a Jaccard index of 0.10. A3 presented shared similarities with both A1 and A2 with respect best pairings with mouse populations. Lastly, A4 shared more similarities with z1 and z2, reinforcing previously observed similarity of A4 to A1. The highest A4 similarity was with z2, with whom it shared 16 markers (*ADGRD1*, *AIF1L*, *CD248*, *CD55*, *DPP4*, *EMILIN2*, *ISLR*, *LIMS2*, *NPR1*, *PAMR1*, *PTGIS*, *RAB32*, *SEMA3C*, *SEMA3E*, *SFRP4*, *WNT2*).

Axis B is related to several populations in mouse, mainly from axis y; although for some populations, like B4, there was interaction with several axes. Population B1 interacted mainly with mouse population z1, with 4 relevant markers (*CXCL2*, *FOSL1*, *GCH1*, *IL6*). B2 and B3 populations showed certain similarities with y4, sharing 7 relevant markers (*APOE*, *C3*, *IL33*, *MGP*, *NFIB*, *SLCO2B1*, *TNFSF13B*). In the case of the B4 population, despite sharing similarities based on the Jaccard indexes with y1, y2, y3 and y4 populations, none of the overlaps was based on a relevant number of common genes (range 2-3). Therefore, it is likely that B4 fibroblast population is not present in the mouse skin.

Axis C in human overlapped with axis w in mouse, which was expected because both are composed of hair follicle-related cells. C2 population is similar to w1/w2 and w/x. In fact, C2 shared 12 markers with w/x (*CHST15*, *COCH*, *CYP1B1*, *DKK2*, *EMID1*, *FIBIN*, *FMOD*, *MAFB*, *MKX*, *NRP2*, *PTGFR*, *TNMD*). Similarly, C1 population shares the most markers with w4, with a total of 18 markers (*ACAN*, *ACTA2*, *ADAMTS18*, *ADAMTS9*, *BCL11B*, *CALD1*, *CCND1*, *COL11A1*, *COL12A1*, *COL8A2*, *CPXM2*, *EDNRA*, *EDNRB*, *EGFL6*, *MEF2C*, *RAMP1*, *TAGLN*, *TENM3*). An exception to the comparison is the w5 population, with no comparable human population, but with a marked transcriptome profile associated with the cell cycle, based on the expression of several markers, including *Mki67*, *Racgap1*, *Cdca8*, *Kif20a*, *Cenpa*.

Axis D presented considerable overlap with y4, y5 and v1 populations. The best fits were D1 with y5 and D2 with v1. y5 shared 9 relevant markers with D1 (*ABCA8*, *APOD*, *COL8A1*, *FOXS1*, *NR2F2*, *P2RY14*, *SOX9*, *TGFBI*, *VIT*), whereas only 2 with D2. Regarding v1, D1 shared 7 relevant markers, whereas whereas D2 shared 19 (*AQP3*, *BNC2*, *CAV1*, *CAV2*, *CLDN1*, *DOCK9*, *EBF2*, *EFNB1*, *GAB1*, *GPC1*, *ITGA6*, *ITGB4*, *KLF5*, *KRT19*, *MTSS1*, *NDRG2*, *SBSPON*, *SLC2A1*, *TENM2*). Therefore, D2 is likely to be associated with v1. With respect to population y4, it presented relatively little overlap with D1 or D2, and (as mentioned before) its overlap with B4 was not considered meaningful.

In summary, we conclude that human and mouse populations share a certain degree of overlap. In some cases, similar populations may be assigned in one-to-one or one-to-two combinations–e.g. A2-x2, C1-w, A4-z1/z2. In other instances, a set of human and mouse populations are comparable as a set and not individually–e.g. A3 and A1 with x1 and x/y, or D1 and D2 with y5 and v1. Finally, in some cases the most likely situation is a lack of comparable inter-species populations, such as E1 in human or y2 and y3 in the mouse.

## DISCUSSION

In the last decade, the advent of novel technologies at the single-cell level resolution has revolutionized the molecular characterization of cellular states and our understanding of their roles in skin function (Ganier, 2024), cutaneous disease progression (Kumaran et al., 2024) and malignancy (Sinha et al., 2024). However, significant work is needed to harmonize findings in-between independent research groups and different technologies and datasets (Phan et al., 2021), so that a consensus roadmap may be used by researchers in the field. In this work, by analyzing a large number of preexisting scRNAseq datasets, we classify dermal fibroblasts into 15 human and 17 mouse populations, each of them defined with a set of robust markers (Tables 1 and 2). Furthermore, we compare human and mouse populations to start to glimpse potential shared and dissimilar functions. While no functional characterization of the resulting clusters has been pursued, an extensive bibliographic research based on marker genes for each population is available on Supplementary Information. Based on our integrative research, dermal fibroblasts seem to show a wider array of functions than predicted.

What are the potential functions of human fibroblasts? A axis populations are major ECM producers. They all express a large set of ECM components and regulator genes. Interestingly, ECM and regulator gene expression seems to be discriminated by their spatial location. For instance, A2 population, which could be defined as “secretory papillary fibroblasts”, secrete collagens, proteoglycans and other components bound to the dermo-epidermal junction (DEJ) and other structures surrounded by Basal Membranes–e.g., *COL4A2/4* (Barbieri et al., 2014), *COL6A1/2/3* (Sabatelli et al., 2011), *COL13A1* and *COL18A1* (Peltonen et al., 1999; Bonnet et al., 2017; Pehrsson et al., 2019), *COMP* (Farina et al., 2006), *POSTN* (Yamaguchi, 2014)–. In contrast, A1/A3/A4 fibroblasts, which are potential “secretory reticular fibroblasts”, secrete entirely different ECM components associated with providing structure and strength to the dermis–i.e. *COL1A2* and *COL3A1* (Henriksen and Karsdal, 2019; Nielsen et al., 2019), *COL12A1*, *PODN*, *DCN* (Winnemöller et al., 1991)–. With regard to the regulation of ECM production, A2 differs from A1/A3/A4 too. Three main families of ECM-regulation components are necessary for proper ECM regulation: (1) pro-collagen cutting enzymes, (2) collagen and fiber maturation enzymes and (3) ECM-degradation enzymes. Both families of fibroblasts express all three families, but their components differ. *PCOLCE* and *PCOLCE2* pro-collagenases are expressed by A1/A3/A4 reticular fibroblasts (Takahara et al., 1994; Steiglitz et al., 2002), in contraposition to *APCDD1* by A2 papillary fibroblasts (Baicu et al., 2012); *LOX*, *LOXL1* and *P4HA2* fiber maturation enzymes are expressed by reticular fibroblasts, whereas *LOXL2* is expressed by papillary fibroblasts (Liu et al., 2004); and, in order to degrade the ECM, reticular fibroblasts express *MMP2*, whereas papillary fibroblasts express *MMP11*. In accordance with the aforementioned definitions, motility and stiffness sensing are also compartmentalized. A1/A3/A4 express *TNXB*, which restricts cell motility, and A2 expresses *TNC*, which favors fibroblast motility (Lethias et al., 2006). Additionally, A4 also expresses elastic fiber-associated genes such as *ELN*, *FBN1* and *EMILIN2* (Lee et al., 2004; Doliana et al., 2001; Schiavinato et al., 2016); and thus could be defined as a “secretory elastic fiber fibroblast” population. A1 and A4 populations express *PIEZO2*, *DBN1* and *TPPP3*, the first one is a mechanosensory channel in Merkel cells (Wu et al., 2017), and the last two of them are bound to F-actin and found in the terminal protrusions (Butkevich et al., 2015; Vincze et al., 2006); A1/A3 and A2 populations express *SGCA/G* sarcoglycans (Noguchi et al., 1995); and A2 exclusively expresses *ANTXR1*, which binds papillary collagen VI (Nanda et al., 2004).

B axis populations are involved in dermal immune regulation (Ascensión et al., 2021). The B1 population might be an “innate/acute immune response fibroblast”, based on the expression of (i) immediate early genes, which activate the expression of necessary target genes (Bahrami and Drabløs, 2016); (ii) acute phase cytokines and chemokines, such as IL6, most of which are necessary to attract innate cells such as neutrophils and monocytes, but also other APCs, and NK and T cells (Liu et al., 2017; Paquet and Pierard, 1996; West, 2019); and (iii) ECM degradation proteins, mainly matrix metalloproteinases and members of the ADAM family (Cabral-Pacheco et al., 2020; Kelwick et al., 2015). B2 and B3 populations also seem to participate in primary responses but are active in secondary and adaptive responses as well, and could be termed as “adaptive immune response fibroblasts”. This occurs either by (i) the expression of specialized immune cell chemo-attractants, such as *CCL19*, *CSF1*, *IL34*, *CXCL12* (Robbiani et al., 2000; Lin et al., 2008; Nakamichi et al., 2013); (ii) adhesion molecules–e.g. *ICAM1*, *ICAM2* and *VCAM1*–, which are responsible for the adhesion and migration of immune cells (Halai et al., 2013; Mantovani and Dejana, 1998), or (iii) cytokines or other ligands that induce cell maturation such as HLAs, *TNFRSF13* or *CD40* (Rickert et al., 2011; Smith, 2005). Lastly, B4 may play a triple role. First, it may serve an anti-fibrotic and pro-regenerative function based on the expression of markers such as *ADA*, *SLPI* or *IGF1*, which have been shown to suppress TGF-β or modulate *ACTA* expression (Ashcroft et al., 2000; Li et al., 2022; Fernández et al., 2013). Second, it shows an anti-inflammatory potential by expressing markers such as *ITM2A* or *PPARG* (Tai et al., 2014; Zhou et al., 2019; Zhang et al., 2021; Jiang et al., 1998; Szatmari et al., 2004). Third, it expresses markers like *GPX3*, involved in the reduction of ROS (Brigelius-Flohé, 2006), MGST1, involved in glutathione metabolism (Pompella et al., 2003), or *MYOC* and *ITM2A*, shown to act as chaperones against ER stress or other scenarios (Anderssohn et al., 2011; Hedlund et al., 2009). The B4 population may also be related to hair follicles (HF), probably acting as immune regulators. It expresses several markers such as *EFEMP1*, *WNT11*, *HSPG2*, *IGF1*, *FGF10* or *MGST1*, which show expression in different areas of the HF (Takahashi et al., 2020; Lim and Nusse, 2012; Panchaprateep and Asawanonda, 2014; Zhang et al., 2018; Tsutsui et al., 2021; Kobayashi et al., 2019). Additionally, some markers like *IGF1* or *FGF7* and *FGF10* have been associated with HF cycle regulation (Panchaprateep and Asawanonda, 2014; Greco et al., 2009; Zhang et al., 2018).

C axis archetypal populations C1 and C2 belong to the HF. Specifically, C1 seem to be dermal sheath fibroblasts expressing *ACTA2* (Mistriotis and Andreadis, 2013); and C2 dermal papilla fibroblasts expressing *CRABP1* and *COCH* (Collins and Watt, 2008; Ahlers et al., 2022). Both populations actively secrete and modify ECM components, most of which are linkers of collagens, SLRPs and other components. In fact, the ECM-related genes expressed by these populations, such as *ACAN*, *F13A1*, *COL11A1*, or *COL24A1*, are different from the ECM found in other dermal layers (Roughley and Mort, 2014; Luo et al., 2019; Nielsen and Karsdal, 2019). The C3 and C5 populations are also associated with HF. C3 is involved not only in the production of specialized ECM, but also in its modulation and degradation, with markers like *TNMD* or *KLK4* (Obiezu et al., 2006; Lin et al., 2017). C3 fibroblasts could be thus termed “HF ECM regulatory fibroblasts”. C5 fibroblasts show Wnt and non-canonical TGF-β signaling, vascular and neural modulation and putative immune functions.

D1 and D2, which are relatively small populations, are conserved in both species and seem to participate in neural- and vascular-related processes, immune signaling, TGF-β and Wnt signaling (further described in the Supplementary Material). D1 has an increased expression in neural and vascular-related genes; whereas D2 has a more apparent expression of genes related to HF. Additionally, each population has certain specific gene families that are more relevant to each of the populations, like collagens and FMO genes for D1 population, and lipid metabolism, caveolae-related genes, or integrin α4β6 complex-associated *ITGA4* and *ITGA6* genes for D2. These populations seem to be relevant for skin appendage homeostasis, probably by interaction with vasculature, nerve endings–e.g. the lanceolate complex, other sensory terminals–, and the Arrector Pili Muscle (Martino et al., 2020). Based on marker expression, we define D1 as “neurovascular fibroblasts” and D2 more generally as “appendage-associated fibroblasts”. The E1 population had a reduced number of relevant markers, which impeded its classification.

In conclusion, dermal fibroblasts are extremely heterogeneous and dynamic cell populations. They should be regarded as a family of cells with diverse roles in maintaining the structure and function of the dermis and the cells and extracellular matrix that lay within it. The consensus single-cell transcriptomic atlas of dermal fibroblasts may be consulted in a user-friendly CellxGene viewer interface, available at https://cellxgene.cziscience. com/collections/3c4f0970-7614-43de-beb7-6128b3cb74ed for any researcher interested in contrasting their data with the populations here defined. This work lays the foundation for the isolation of distinct populations and the design of functional studies that proof or disproof each of the provided hypotheses, which importantly can now be pursued in mouse and human in a comparative manner.

## MATERIALS AND METHODS

### Dataset downloading and fastq processing

Individual dataset downloading is explained in detail in the Supplementary Information. In general, datasets were downloaded in fastq format (if available) and processed using loompy fromfq pipeline with GRCh38/GRCm38 genomes as a default. If fastq not available, raw count matrices were used instead. Due to the large volume of samples, a unified metadata scheme was applied to all the datasets, which is described in the Supplementary Information.

All datasets were processed using the following scheme:

1. Plotting QC metrics and thresholding: sc.pp.calculate_metrics was used to calculate metrics, pct_counts_mt VS total_counts and n_genes_by_counts VS total_counts scatter plots were used to select the threshold of number of genes and percentage of mitochondrial counts. Generally, if the distribution was unimodal, up and down tails were removed, and if the distribution was multimodal, the top part was selected. If there was more than one sample in the dataset, these stats were plotted per sample and, depending on the differences between samples, either a common set of cutoffs was set, or different cutoffs were set per sample.
2. Basic transformations: the dataset was normalized, log1p-transformed, and genes with no counts were removed.
3. PCA, harmony integration, and kNN calculation: sc.pp.pca was used to calculate PCA with 30 variables. If the dataset contained different samples, harmony was used with sce.pp.harmony_integrate function. To calcu<u>late the</u> kNN graph, sc.pp.neighbors function was used with cosine metric and the number of neighbors as 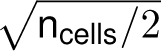.
4. Major cell type assignment (human): UMAP–sc.tl.umap– and *leiden*–sc.tl.leiden– were used to see all major cell types. *Leiden* was run with a high resolution (3 to 10, depending on the dataset). The marker-to-population algorithm (defined afterwards) was used to set different major cell types in an unsupervised manner. The markers used for each cell type are described in the Supplementary Information. Regarding the characterization of fibroblasts, *LUM*, *PDGFRA*, *COL1A1*, and *DCN* genes were used to find this population across different single-cell studies.
5. Fibroblast processing: to do the subprocessing, genes with no expression on fibroblasts were filtered, and PCA and kNN were run as before. *triku* (Ascensión et al., 2022c) was used for feature selection, with default settings. The output from this method is a boolean array indicating if each gene is selected or not as highly variable. With this information, PCA, harmony and kNN calculation were rerun to produce the definitive set of neighbours for the following steps.
6. Analysing fibroblast populations: to obtain the different populations, UMAP and *leiden* were run again, with min_dist and resolution values adjusted for each dataset. The goal was to obtain a high number of clusters to run the marker-to-population algorithm on them. Once populations were obtained, if there was no need to reanalyze populations or investigate new arising populations, the processing was concluded.
7. Additional step - detecting new populations: sometimes the marker-to-population algorithm assigned the same label to more than two clusters, or left some cluster unassigned. In these cases, it is possible that a new cluster is being discovered. To check that possibility, DEGs of that population were calculated, and the marker-to-population algorithm was used with a set of markers. If the algorithm assigned the cluster correctly, it was a putative population. To assert that this putative new population was verifiable, it had to be assigned in other datasets as well.
8. Additional step - removing unwanted populations: In some cases, when assigning fibroblast populations, unwanted populations appeared–e.g. stressed cells, non-fibroblast cells, etc.–. In these cases, these cells were removed from the AnnData, and the whole processing of fibroblast cells was redone.

### Pipeline to obtain a robust set of markers and populations

The pipeline for robust markers and populations consists of a combination of the marker-to-population and population-to-marker algorithms (described in the Supplementary Information) that tries to minimize the bias of supervised selection.

The pipeline consists of three steps:

1. Given a set of handpicked populations and genes, use the marker-to-population algorithm to assign clusters from each dataset their most probable population.

- If one or more unassigned populations arise, gene markers are plotted to see if the cluster belongs *de visu* to one of the populations. If so, the threshold parameter has to be tuned.
- If no markers are expressed in unassigned populations, DEGs from that population that are representative markers can be chosen and, if other clusters in other datasets follow the same pattern, the new population and its markers can be added to the dictionary.
2. From the set of populations, applying the population-to-marker algorithm yielded a “robust” set of markers for each of the populations across datasets. This set was less biased towards a specific dataset than the first set of markers used for the first marker-to-population algorithm. Nonetheless, a change of the initial conditions of the pipeline will considerably affect the set of markers obtained after this algorithm.
3. Finally, with the “robust” set of markers the marker-to-population algorithm was rerun to produce a “robust” set of populations.

### Comparison of human and mouse fibroblast populations

To perform mouse-human comparisons two approaches were used. First of all, and to map genes between the two organisms, a human-mouse dictionary was created using a pan-organism homology table downloaded from the Mouse Genome Database (MGD), Mouse Genome Informatics, The Jackson Laboratory, Bar Harbor, Maine, available from this link (https://tinyurl.com/fc6hrv8m).

#### Integration of human and mouse fibroblasts from the same datasets

Mouse and human populations from Boothby et al. (2021) and Vorstandlechner et al. (2021) were integrated using *harmony* batch effect correction method with default settings. Additionally, *bbknn* was also used but produced similar results. The integration was evaluated visually.

#### Comparison of human and mouse markers

Mouse and human populations were compared by measuring the overlap of their markers. This method was also applied, as a control measurement, within human and mouse populations as well.

To perform the comparisons we selected the optimal number of markers to compare using the procedure described in the Supplementary Information. Then, the matrix of Jaccard indexes between each pair of populations was computed. From that matrix, the most relevant population pairs were selected manually, and for each pair, relevant markers were selected.

A marker was chosen to be *relevant* if it was exclusive of that population. This step was necessary to remove markers that, despite being differentially expressed in a given population, were not exclusively expressed.

### Robustness of semi-supervised algorithm

To check the robustness of the marker-to-population algorithm we applied a methodology similar to the classical jackknife resampling method (Quenouille, 1956). For 30 iterations, and for each dataset, we sampled a number of the cells without repetition and rerun the marker-to-population algorithm. The sampling was stratified across populations so that all populations were constantly represented. Sampling percentages were set to 50%, 70%, 80%, 95% and 99%, and the last one was used for the Results section.

After 30 iterations, a table per dataset with as many rows as the number of cells of the dataset, and 30 columns was obtained. For each cell, the population to which that cell was assigned in each iteration was stored. In some iterations, and for some cells, a NaN value was stored because the marker-to-population algorithm was not run in that sample of cells.

For each cell, the quotient between the number of iterations where the assigned cluster was the same as the original, and the number of assignments performed was calculated and defined as *robustness score*. For instance, for one cell whose original population was A2, and was assigned 20 times to A2, 7 times to A1, and it was not sampled in 3 iterations, its robustness score was calculated as 20/27=0.74.

Additionally, assignment to other populations different to the original cluster was also computed. To do that, an adjacency matrix was created where the rows represent the list of original populations, and the columns represent the list of populations assigned for that dataset. For each row-column combination, the quotient between (1) the number of instances of assignment to the column population when the original labelling was the row population and (2) the number of assignments of cells labelled with the row population was calculated.

## DATA AVAILABILITY STATEMENT

Jupyter notebooks used to analyse the fibroblast populations are stored, in raw .ipynb and .html form, at the following Zenodo repository (doi: 10.5281/zenodo.7492966), and at the alexmascension/fibroblast-population-detection Github repository.

The source code of the marker-to-population algorithm is available at the alexmascension/cell_asign GitHub repository.

Raw and processed .h5ad AnnData files, as well as the adata file for the combined datasets, can be found at the Zenodo repository for human datasets (doi: 10.5281/zenodo.7785089) and the Zenodo repository for mouse datasets (doi: 10.5281/zenodo.7785085).

The *CellxGene* viewer with the processed human and mouse datasets is available through the CZIscience CellxGene platform. Preprocessing for this section is indicated in the Supplementary Information.

The metadata file with the information of all human and mouse datasets is available at the following GitHub repository, and at the Zenodo repository (doi: 10.5281/zenodo.7492966) together with the HTML and .ipybn files.

Lastly, information pieces from the secondary analysis are available in a Google Docs spreadsheet–updatable version–, and at the Zenodo repository (doi: 10.5281/zenodo.7492966)–fixed version, last updated as of 2023/03/30–.

## Supporting information

robust_markers_mouse

robust_markers_human

## ACKNOWLEDGEMENTS

We would like to thank Jennifer Yu-Sheng Chien for curating and preparing the datasets for the upload to the CellxGene platform.

The research was financially supported by grants “PI22/01247”, “CERT22/0003” and “PT23/00142” from Instituto de Salud Carlos III (ISCIII, Spain) and co-funded by the European Union. AMA was funded by a fellowship provided by the Department of Education, Universities and Research of the Basque Government (*Programa Predoctoral de Formación de Personal Investigador No Doctor* ; PRE_2018_1_0008, PRE_2019_2_0233, PRE_2020_2_0081, PRE_2021_2_0111) and by a Grant from the Educational Department of the Basque Government (IKUR-Nanoneuro).

## AUTHOR CONTRIBUTIONS

Conceptualization: AI, Methodology: AMA, Software: AMA, Formal Analysis: AMA, Investigation: AMA, Resources: AI, Data Curation: AMA, Writing – Original Draft Preparation: AI and AMA, Writing – Review and Editing: AI and AMA, Visualization: AMA, Supervision: AI, Project Administration: AI, Funding Acquisition: AI.

## Supplementary Materials and Methods

### Dataset downloading information

Human datasets were downloaded from the following places:

- Ahlers et al. (2022): fastq files were downloaded from SRA archive (accession number PRJNA754272) and were processed using kallisto, embedded in loompy pipeline.
- Billi et al. (2022): files were downloaded in 10X mtx form (unprocessed filtered file) from GEO database (accession numbers GSE186476).
- Boothby et al. (2021): files were downloaded in h5 form (unprocessed filtered file) from GEO database (accession number GSE183031).
- Burja et al. (2022): fastq files were downloaded from SRA archive (accession number ERR9121019) and were processed using kallisto, embedded in loompy pipeline.
- Deng et al. (2021): files were downloaded in 10X mtx form (unprocessed filtered file) from GEO database (accession numbers GSM4994382, GSM4994383 and GSM4994384 for scar and GSM4994379, GSM4994380 and GSM4994381 for scar).
- Gao et al. (2021): files were downloaded as a unprocessed filtered loom from GEO database (accession number GSE162183).
- Gaydosik et al. (2019): files were downloaded in csv form (unprocessed filtered file) from GEO database (accession numbers GSM3679038, GSM3679039, GSM3679040, GSM3679041).
- Gur et al. (2022): files were downloaded in txt form (unprocessed filtered file) from GEO database (general accession number GSE195452).
- He et al. (2020): fastq files were downloaded from SRA archive (accession numbers SRR11396162, SRR11396164, SRR11396166, SRR11396167, SRR11396168, SRR11396170, SRR11396171, SRR11396175) and were processed using kallisto, embedded in loompy pipeline.
- Hughes et al. (2020): files were downloaded in csv form (unprocessed filtered file) from GEO database (accession number GSE150672). Metadata files indicating sample and disease were downloaded from this website (https://tinyurl.com/4e345da2).
- Kim et al. (2020a):fastq files were downloaded from SRA archive (accession numbers SRR9307706, SRR9307707, SRR9307708, SRR9307709, SRR93077010, SRR9307711) and were processed using kallisto, embedded in loompy pipeline.
- Liu et al. (2021): fastq file processing produced a faulty count matrix and, therefore, the count matrix was extracted from the Seurat object provided under personal request to the authors, which can be downloaded from the following link (https://tinyurl.com/3bvzajkn).
- Mariottoni et al. (2021): files were downloaded in tsv form (unprocessed filtered file) from GEO database (accession number GSM5352395).
- Mirizio et al. (2020): fastq files were downloaded from SRA archive (accession numbers SRR12955136 to SRR12955159), were trimmed using seqtk (arguments -e 124 -e 52) and were processed using kallisto, embedded in loompy pipeline.
- Reynolds et al. (2021): fastq files were downloaded from EBI repository E-MATB-8142, and were were processed using the loompy fromfq pipeline. For other annotations and analysis of other populations, the processed h5ad adata from Reynolds et al. (2021) was downloaded from the Zenodo repository (ID: 4536165).
- Rindler et al. (2021) Rindler et al. (2021): files were downloaded in tsv form (unprocessed filtered file) from GEO database (accession numbers GSM5534590 to GSM5534593).
- Solé-Boldo et al. (2020): fastq files were downloaded from SRA archive (accession numbers SRR9036396 and SRR9036397) and were processed using kallisto, embedded in loompy pipeline.
- Tabib et al. (2018): data file in csv form was downloaded from Pittsburgh University website (link) as well as the metadata file (link).
- Tabib et al. (2021): files were downloaded in h5 form (unprocessed filtered file) from GEO database (accession numbers GSM4115868, GSM4115870, GSM4115872, GSM4115874, GSM4115875, GSM4115876, GSM4115878, GSM4115880, GSM4115885, GSM4115886).
- Jones et al. (2022): files were downloaded in fastq form from Amazon Web Services platform (under previous access request). Fastqs from samples *TSP10_Skin*, *TSP14_Abdome*, *TSP14_Chest* were downloaded and processed using kallisto, embedded in loompy pipeline.
- Theocharidis et al. (2020): 10X mtx raw filtered files were obtained by personal request, and downloaded from their Dropbox service. Sample IDs from controls were *H1_080717*, *H2_091117*, *H3_091117*, *H4_100317*.
- Theocharidis et al. (2022): files were downloaded in csv form (unprocessed filtered file) from GEO database (accession numbers GSM5050521, GSM5050534, GSM5050538, GSM5050540, GSM5050542, GSM5050548, GSM5050552, GSM5050553, GSM5050555, GSM5050556, GSM5050560, GSM5050564, GSM5050567, GSM5050568, GSM5050574, GSM5050522, GSM5050524, GSM5050525, GSM5050526, GSM5050528, GSM5050529, GSM5050562, GSM5050565, GSM5050570, GSM5050572, GSM5050523, GSM5050527, GSM5050531, GSM5050532, GSM5050536, GSM5050537, GSM5050539, GSM5050541, GSM5050547, GSM5050566, GSM5050569, GSM5050573, GSM5050530, GSM5050533, GSM5050535, GSM5050557, GSM5050558, GSM5050559, GSM5050563).
- Vorstandlechner et al. (2020): processed loom file was obtained under personal request.
- Vorstandlechner et al. (2021): files were downloaded in tsv form (unprocessed filtered file) from GEO database (accession numbers GSM4729097 to GSM4729102).
- Xu et al. (2021) Xu et al. (2021): files were downloaded in 10X mtx form (unprocessed filtered file) from OMIX database (accession number: OMIX691).

Mouse datasets were downloaded from the following places:

- Abbasi et al. (2020): files were downloaded in 10X mtx form (unprocessed filtered file) from GEO database (accession number: GSM2910020).
- Boothby et al. (2021): files were downloaded in 10X mtx form (unprocessed filtered file) from GEO database (accession numbers: GSM5549901 and GSM5549902).
- Buechler et al. (2021): fastq files were downloaded from ArrayExpress database (accession number E-MTAB-10315) and were processed using kallisto, embedded in loompy pipeline.
- Haensel et al. (2020): files were downloaded in 10X mtx form (unprocessed filtered file) from GEO database (accession numbers: GSM4230076 and GSM4230077).
- Joost et al. (2020): files were downloaded in 10X mtx form (unprocessed filtered file) from GEO database (accession numbers: GSM4186888 to GSM4186893).
- Phan et al. (2020): files were downloaded in loom format (unprocessed filtered file) from GEO database (accession numbers: GSM4647788 to GSM4647790).
- Shin et al. (2020): files were downloaded in mtx form (unprocessed filtered file) from GEO database (accession numbers: GSM3177991, GSM3177992, GSM4155928, GSM4155929).
- Shook et al. (2020): fastq files were downloaded from SRA database (accession numbers SRR10480641 and SRR10480643 to SRR10480646) and were processed using kallisto, embedded in loompy pipeline.
- Vorstandlechner et al. (2021): unprocessed filtered mtx files were provided under personal request.

### Dataset metadata

Datset metadata contains the following entries:

- Author: Surname of the first author of the publication.
- Year: Registered year of last publication date.
- DOI: DOI of last registered publication.
- Accession (General): Accession number of all the samples.
- Accession (Sample): Accession number of each sample.
- Accession (SRR): If the file is a FASTQ, SRR number of the SRA archive, if existing.
- Data origin: Type of raw data–e.g. FASTQ, Raw count matrix–.
- Aligner: Alignment software used for preprocessing.
- Genome: Genome used during the alignment.
- Donor identifier: Identification number for the donor. If none, natural numbers are used.
- Sample identifier: Identification tag of the sample, as of the dataset repository.
- Internal sample identifier: Sample identifier used within the anndata file. Sometimes, the internal sample identifier differs from the sample identifier because the sample identifies is too long or missing.
- Downloaded: Raw data files were downloaded (boolean).
- Analysed: Raw data files were processed (boolean).
- Exclusion reason: If data were not downloaded or processed, indicate the reason if necessary.
- Library preparation: Library preparation method.
- Sequencer: Sequencer used after library preparation.
- Organism: mouse or human.
- Age: Individual age–in days, weeks, months or years depending on the organism–.
- Age (mean): Sometimes, due to a lack of information on individual ages, an age range is provided. For calculation measures, the mean age is computed.
- Age format (y/m): Format of the age stated before.
- Gender: Gender of the individual.
- Race: Race of individual.
- Ethnicity: Ethnicity of the individual.
- Sample location: Body location where the sample was extracted from.
- Condition: Condition of individual, either healthy/control, wounding, or any disease.
- Condition (other): Notes on the condition–e.g. severity of the disease, or days after wounding–.

### Human and mouse markers used to characterise major skin populations

The list of human populations and markers is the following:

- fibro: *LUM*, *PDGFRA*, *COL1A1*, *DCN*, *SFRP2*, *APOE*, *APOD*, *FN1*
- fibro - ANGPTL7: *ANGPTL7*, *ENTPD2*, *ETV1*, *C2orf40*, *SCN7A*, *SOX8*
- F: *B4GALT1*, *TMSB4X*, *PPP1CB*, *WTAP*, *PTPRS*, *CTNNB1*, *INSR*, *BICC1*, *CTNNB1*
- melanocyte: *MLANA*, *PMEL*, *TRIM63*, *QPCT*, *PLP1*, *TYRP1*
- neuro: *GPM6B*, *PLP1*, *S100B*, *SCN7A*, *NRXN1*, *GFRA3*, *MPZ*
- secretory cell: *KRT7*, *KRT8*, *KRT18*, *KRT19*, *DCD*, *SCGB2A2*, *PPP1R1B*, *MUCL1*, *AZGP1*, *SCGB1D2*, *PDCD4*, *TSPAN8*
- muscle: *TAGLN*, *DES*, *PCP4*, *ACTG2*, *CNN1*, *CSRP1*, *TPM1*, *SYNPO2*, *PRUNE2*, *SORBS1*, *P2RX1*
- peri - CYCS: *TAGLN*, *CRISPLD2*, *CYCS*, *VDAC1*, *RHOB*, *SORBS2*, *PLEKHO1*, *CNN1*, *DNAJB9*, *CSRP2*
- peri - RERGL: *TAGLN*, *ACTA2*, *CRISPLD2*, *RERGL*, *BCAM*, *ADIRF*, *NET1*, *ARPC1A*, *PLN*
- peri - RGS5: *ACTA2*, *RGS5*, *ABCC9*, *HOPX*, *ARHGDIB*, *KCNJ8*, *FXYD6*
- peri - ZFP36: *RGS16*, *NR2F2*, *TGFBI*, *CCL8*, *RERG*, *HOPX*
- endo artery: *PLVAP*, *CLDN5*, *PECAM1*, *IGFBP3*, *SRGN*, *SEMA3G*, *RHOB*, *HEY1*
- endo capillary: *PLVAP*, *CLDN5*, *PECAM1*, *SELE*, *SOCS3*, *CDKN1A*, *NFKBIA*, *DNAJB1*, *ATF3*
- endo venule: *PLVAP*, *CLDN5*, *PECAM1*, *CYP1B1*, *CLU*, *PERP*, *VWF*, *IER3*, *TSC22D3*
- lymph: *CCL21*, *LYVE1*, *CLDN5*, *TFF3*, *MMRN1*, *EFEMP1*, *FGL2*, *TFPI*, *MAF*
- krt basal: *KRT14*, *COL17A1*, *KRT5*, *KRT15*, *DST*, *PDLIM1*
- krt channel: *KRT23*, *GJB6*, *GJB2*, *CDA*, *MMP7*, *PNLIPRP3*
- krt spinous: *KRT1*, *KRT10*, *DMKN*, *KRTDAP*, *CHP2*, *LYPD3*
- krt gran: *FLG*, *NCCRP1*, *CNFN*, *TGM1*, *CST6*, *KLK7*
- immune: *TPSB2*, *TPSAB1*, *HLA-DRA*, *FCER1G*, *CD74*
- T CD4+: *CD52*, *CD3D*, *TRAC*, *TCF7*, *CD4*, *IL7R*, *CD40LG*
- T CD8+: *CD52*, *CD3D*, *TRAC*, *CD8B*, *THEMIS*, *CD8A*, *FOXP3*, *CCR4*, *RORC*, *TIGIT*
- plasma cell: *SDC1*, *SLAMF7*, *TNFRSF17*, *PTPRC*, *CXCR4*, *MYH9*, *PRDM1*, *CD38*, *CD27*, *IGHG1*
- dendritic cell: *GZMB*, *MRC1*, *XCR1*, *CLEC9A*, *IRF8*, *EPCAM*, *CD1B*, *STMN1*, *IDO1*
- APC: *HLA-DQA1*, *HLA-DRB6*, *TYROBP*, *FCER1G*, *AIF1*
- mast cell: *IL1RL1*, *CPA3*, *HPGDS*, *TPSB2*, *HPGD*, *RGS13*, *CTSG*, *TPSAB1*, *GATA2*
- NK cell: *NCAM1*, *XCL1*, *CD38*, *CD7*, *IL18R1*, *KLRF1*, *KLRK1*
- mt: *MTND2P28*, *MTND4P12*, *MTCO1P40*, *ADAM33*, *RN7SL2*, *MTRNR2L6*,
- eritro: *HBB*, *HBA2*, *HBA1*, *HBD*

The list of mouse populations and markers is the following:

- peri: *Rgs5*, *Myl9*, *Ndufa4l2*, *Nrip2*, *Mylk*, *Rgs4*, *Acta2*, *Sncg*, *Tagln*, *Des*, *Ptp4a3*, *Myh11*
- endo: *Pecam1*, *Cdh5*, *Egfl7*, *Cd36*, *Srgn*, *Adgrf5*, *Ptprb*, *Scarb1*, *Plvap*, *Grrp1*, *C1qtnf9*, *Mmrn2*, *Flt1*
- kerato: *Krt14*, *Krt15*, *Perp*, *S100a14*, *Ccl27a*, *Gata3*, *Dapl1*, *Rab25*, *Ckmt1*, *Col17a1*, *Serpinb5*
- kerato *Gjb2*: *Ucp2*, *Krt71*, *Gjb2*, *Ahcy*, *Acaa2*, *Cbs*, *Slc3a2*, *Serpina11*, *Lap3*, *Gss*, *Basp1*
- fibro: *Dcn*, *Pdgfra*, *Lum*, *Col1a1*, *Col1a2*
- fibro_2: *Ncam1*, *Ptch1*, *Trps1*, *Col11a1*, *Wif1*
- fibro_acan: *Acan*, *Col2a1*, *Col11a1*, *Col9a1*, *Snorc*, *Col9a3*, *Mia*, *Cnmd*, *Ucma*, *Chad*
- T cell: *Rac2*, *Ptprcap*, *Il2rg*, *Cd3g*, *Skap1*, *Hcst*, *Ctsw*, *Ets1*, *Cd3d*, *Ctla2a*, *Cd2*
- APC: *Tyrobp*, *Cd74*, *H2-Aa*, *H2-Eb1*, *Ctss*, *Spi1*, *Napsa*, *Cd68*, *Lyz2*, *Csf2ra*
- lymph: *Ccl21a*, *Egfl7*, *Mmrn1*, *Nsg1*, *Meox1*, *Gimap6*, *Kdr*
- melano /schwann: *Syngr1*, *Pmel*, *Mlana*
- myo: *Tnnt1*, *Tnnt2*, *Tnnt3*, *Tnnc2*, *Acta1*, *Myl1*, *Tnni2*, *Tcap*, *Eno3*, *Myoz1*
- neural: *Itgb8*, *Plp1*, *Ptn*, *Egfl8*, *Chl1*, *Cadm4*, *Sox10*, *Cdh19*, *Snca*

### Confirming the expression of genes in fibroblast subpopulations and general cell types

After running the robust population and marker pipelines, we produced a table of relevant information for each gene so that it can be useful to determine the functions of each of the populations based on their gene signature. To construct this table, we need to assign the populations in which that gene is expressed first.

The chosen plots to this step are a combined dot plot and UMAP plot that show the expression pattern of each gene across datasets. The size of the dot in the dot plot represents the proportion of cells within each subpopulation that expresses that gene, and the colour represents the level of expression. To better visualise the proportion, the value that is represented in the dot plot is scaled as 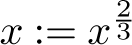, which allows a better discrimination of genes expressed in fewer cells. The aim of this dot plot is to have an overall view of the expression patterns of each gene, especially when the pattern of expression is diffuse or complex.

As a complement to the dot plot, the UMAP plot with the expression of a gene in all dataset supports the findings from the dot plot, and allows to find differences of expressions across datasets.

All this information will be used to create the table with gene information, in which we will include the following information:

- Gene name
- DEG-based location: to create this column, genes appearing as the first 30-50 markers in the dictionary obtained from the pipeline described above are included in this column. If a gene is a marker of more than one population, this information is reflected. For instance, a gene with the label “A1 | B2 | C1” is a marker or A1, B2 and C1 populations at the same time, whereas a gene with the label “D1” is only a marker of the D1 population.
- UMAP-based fibroblast location: this column represents the pattern of expression of each gene within different fibroblast populations. If two populations share the same level of expression, the symbol *∼* is used; whereas if one population has a greater, or clearer, expression than the other, the symbol > is used. For instance “A1 *∼* B2 > C1 *∼* B4” indicates that the gene is more expressed in A1 and B2 than in C1 and B4; but it is expressed at a similar level in A1 and B2, and in C1 and B4. If there is a thorough expression of the gene, but it is apparent in one or more populations, “rest” is added to the comparison. For instance, “A1 > B2 *∼* B3 > rest” indicates that the gene is clearly expressed at A1, then at B2 and B3, and there is a minor, basal expression in the rest of the cells.
- UMAP-based general location: similar to the previous column, a combination of dot plot- and UMAP-based visualisation are used to determine the expression patterns of each gene within the major cell types of the skin.
- Location: location of the gene product within the cell (extracellular, endoplasmic reticulum, plasma membrane, Golgi, nucleus, etc.). This information is obtained from GeneCards website (Safran et al., 2021).
- Basic gene information: in order to have a quick view of the putative functions of the gene, summaries from Entrez gene, GeneCards and Uniprot are included. All the summaries are joined, and extracted from, the GeneCards website (Safran et al., 2021).

It is important to remark that the expressions used in the UMAP-based fibroblast and general location columns are arbitrarily depicted based on the dot plot and UMAP plots. Unfortunately, this level of annotation across such heterogeneous datasets requires a subjective and arbitrary classification. Nonetheless, this notation is really helpful to have a quick view of the gene expression patterns without constantly needing to resort to visualisation methods.

### PAGA and population-relationship graphs

One important step in the secondary analysis is to detect relationships between individual populations. This is important because fibroblasts, like any other cell type, are not independent entities, but rather interact within the tissue, either with other fibroblasts or other cell types. Therefore, fibroblast subpopulations, as well as independent axes will likely interact with each other, or share more similarities between them.

To analyse the similarities between fibroblast populations, PAGA algorithm is used (Wolf et al., 2019). The objective of the algorithm is to extend the idea behind the neighbouring concept that is applied cell-wise to clusters. Using the kNN information computed during the processing steps, and for a given set of clusters, PAGA produces an adjacency matrix indicating relationships between each cluster. With this information a graph between clusters is created where, the greater the connection between two nodes, the greater the similarity between two clusters.

To simplify the relationships within the PAGA graph a secondary tree graph is constructed, which explains the major relationships, and forces interactions to be in that tree structure. This graph comes with the limitation of “breaking” cliques or looped cluster relationships, but allows for a clearer lecture of an otherwise sometimes complex graph. Additionally, this graph tends to preserve relationships that are clearly meaningful.

Thus, for each dataset these two PAGA graphs are computed. This allows to analyse the heterogeneity of populations and interactions between populations across datasets.

In order to integrate all the information, a joined graph is built. To build this graph, to the PAGA adjacency matrix from the tree graph of each dataset, we remove interactions with values smaller than a threshold–0.6 by default–, and then compute a power of that number–between 1 and 2, depending on the graph–. The first step removes loose relationships that tend to saturate the joint graph, and the second step exacerbates differences between clearly meaningful cluster relationships and less-relevant relationships, making them stand out in the figures.

### Marker-to-population algorithm

The aim of this algorithm is to, given a dictionary of populations and their respective markers, generate a mapping between the clusters of the dataset and the populations. This correspondence is not univocal: it is possible to assign one population to several clusters, but it is not possible to assign several populations to one cluster. Additionally, it is possible to obtain clusters that are not assigned, that is, there is no population whose markers are not sufficiently representative of that cluster. These populations are usually relevant at the initial iterations, because they might be novel populations that were unnoticed in previous datasets.

The algorithm follows these steps:

- For each category, a matrix of shape the number of cells in the AnnData by the number of markers of that category, M_0_, is created.

For each gene in M_0_, the gene expression array is multiplied by the neighbour matrix, to produce a matrix of cumulative counts in the kNNs. The aim of this product is to reinforce the expression of local genes. This kNN matrix is divided by the number of neighbours, and the result is stored in the corresponding column of M_0_.

- Once M_0_ has been completed, the mean is extracted, column-wise. This produces a column matrix with the mean kNN values across genes, m_1_. We create the matrix M_1_, resulting of concatenating m_1_ matrices with all populations. Therefore, the M_1_ shape is the number of cells in the AnnData by the number of populations.
- The cluster labelling from *leiden* is added to M_1_, and this information is collapsed into a M_2_ matrix, whose dimension is the number of clusters by the number of populations. To collapse the information of several cells from one cluster in M_1_ into one number in M_2_, either the percentile defined in the quantile_gene_sel argument, the CV, or the maximum value are computed.
- For the M_2_ matrix with the selected collapsing function, the assigned population is the one with the highest value across clusters.

If none of the populations achieves a minimum value established in min_score the population assigned to the cluster is undefined.

### Population-to-marker algorithm

The aim of this algorithm is to, given a set of datasets with labelled populations, produce a list of markers that are commonly expressed in all datasets for each population. This algorithm arises because of the scenario of certain genes being markers of a specific dataset, and not in others, due to specific characteristics derived from each dataset. In order to avoid these types of genes, this algorithm achieves robustly-expressed sets of markers across all datasets instead.

For each of the populations, the algorithm follows these steps:

- First, a common set of markers across all datasets is extracted. For that, the DEGs of that population in each dataset are obtained, following a threshold–by default, p-value < 0.01–. Then, genes that follow unwanted patterns–ribosomal genes, mitochondrial genes, *MALAT1*, *S100* genes, etc.– are excluded. The non-excluded set is the set of genes that will be used for evaluation.
- For each gene and each dataset, two values are stored: the scoring value used in the DEG calculation–by default, the score value from sc.tl.rank_genes function–, and the sum of the expression of that gene in the cells within that population. This will produce two matrices of shape the number of genes from the first step by the number of datasets that will store these values.
- For each of these two matrices (M)–the one based on the DEGs, and the one based on the expression–, a mean value is computed across datasets, so that we have a value per gene. This mean is pondered (µ*^∗^*) using the sigmoid function of the value, so that expression values of datasets with greater quality–overall, genes with more localised expression and, thus, higher scores–are dampened and thus the selected markers are not completely biased towards one or two datasets:

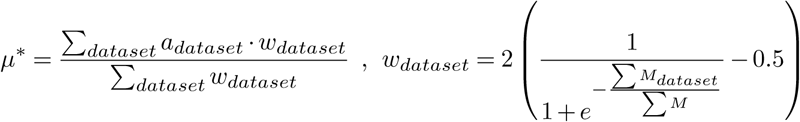

Where a*_dataset_* is the scoring value is in the DEG calculation, and w*_dataset_* is the weight computed for the pondered mean, which is based on a*_dataset_*.
  - Once mean values for genes with score based on DEGs (µ⃗*^∗^_DEG_*) and based on the expression (µ⃗*^∗^_expr_*) are computed, a score value is obtained by applying the following expression:

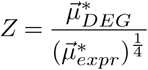

That is, mean score values are divided by mean expression values to avoid selecting for genes that have higher thorough expression–which makes the DEG score higher–but which are not specifically expressed in the population of interest. Finally, in order to avoid genes selected as DEGs but with a extremely small expression, only Z values with µ*^∗^_expr_ ≥* 0.035 are chosen. This allows, therefore, to choose from genes with middle expression values.

In the end, the population-to-marker algorithm produces a dictionary with the representative populations and a data frame of the markers–with scores and Z–.

### Selection of best N for Jaccard index in mouse-human marker overlap analysis

The first step to do the comparisons is to select the number of markers from each population that are going to be compared. To do that we applied the following algorithm:

- For each pair of populations, given a number of markers N, calculate the Jaccard index between the first N genes in one population and N genes in the other. All Jaccard index values are stored in a matrix of indexes, M*_N_* .
- Compute the maximum trace of M*_N_*. To do so, we use the Hungarian method from the *munkres* package (Munkres, 1957). This algorithm finds the permutations of a matrix to satisfy that its trace will have the minimum or maximum value. Therefore, obtaining the trace of M*_N_* after applying the Hungarian method will provide a value of wellness of overlap between the most relevant pairs of populations. This trace value is stored alongside the N value. The higher the trace, the more general overlap between the sets of markers across different populations occurs.
- The previous two steps are applied with a range of N values (10 to 300). To select the best value, we plotted N values and their respective traces and analysed the trend line. Generally, there is a knee point where trace values plateau. This was the selected N for the analysis.
- Once the best N was selected, the matrix M*_N_* was plotted and correspondences were found. In human-human and mouse-mouse comparisons, a superficial analysis was done to assert that comparisons are logical, and in line with the combined PAGA graph.

### Preprocessing of adatas for CellxGene

In each collection, a combined UMAP of all datasets is shown. The UMAP can display information about discrete categories such as age, author, and other categories described in the following paragraph. Expression of individual genes can be displayed, and DEGs between manually selected subpopulations can be performed. The expression values are based on the *processed* matrix counts. However, to unify value expression across datasets, mean-variance scaling from *scanpy* sc.pp.scale() function is performed. The arctan function is applied afterwards to remove outlier weight, whose range is [-1.5, 1.5]. This double-step transformation allows for the comparison of gene expression across all datasets.

## Supplementary Results

### Human-human comparisons

Figure S15 shows the heatmap of Jaccard index values with the top 125 markers of each population. Most of the conclusions from the heatmap are shared based on PAGA graphs, and UMAPs commented before. These results are expected and confirmatory to some extent since both UMAP and PAGA information is based on the transcriptomic profiles of the populations.

- A axis is highly isolated from the rest of the clusters, except for A2 population, which shares some markers with C and E axis (*∼*0.05), indicating a possible interacting functionality between them. However, its relatively low similarity profile with C1, C3 and E1 makes it a somewhat “independent” population.
  – A3 population is a bridge between A1 and A2, being more *similar* to A1 (0.27) than to A2 (0.05). The most relevant overlap markers between A1 and A3 are *AOX1*, *ARFGEF3*, *COL14A1*, *CORIN*, *GLRB*, *OMD* and *SVEP1*.
  – A4 overlaps both with A1 and A3, but the overlap is more apparent with A1 (0.19) than with A4 (0.08). The most relevant markers between A1 and A4 are *ACKR3*, *CD12*, *CD55*, *CD70*, *CLEC3B*, *DBN1*, *FBN1*, *GPX3*, *IGFBP6*, *ISLR*, *LGR5*, *SLPI*.
  – No overlap between A1 and A2 is found, and some overlap between A2 and A3 is found (0.05).
  – Relevant markers overlaping between A2 and C1/C3 are *COL21A1*, *COL6A1/3*, *COL7A1*, *COMP LAMC3*, *LOXL2*, *PTK7*, and *SPON1*.
- B axis, similar to A axis, looks independent from most of the other axes.
  – It is clear again that B3 acts as a bridge cluster between B1 and B2, with a more apparent overlap with B2 (0.36) than with B1 (0.08). Markers shared between B2 and B3 are *ADRA2A*, *C3*, *CCL19*, *CTSC*, *CX3CL1*, *ICAM2*, *IGFBP3*, *IL34*, *NLGN4X*, *PTGDS*, *RBP5*, and *TNFSF13B*; and markers shared between B1 and B3 are *CCL2*, *CXCL1/2/3*, *GEM*, *HAS2*, and *TNFAIP3*.
  – B1 and B2 clusters are not related–similarly to A1 with A2–, and B2 is related to B4 as well (0.11): *ABCA8/10*, *APOC1*, *C7*, *EPHX1*, *GGT5*.
  – Despite the relative independence of the B axis, B1 has a partial overlap with D1 (0.04), and B4 has partial overlaps with D1 (0.07), D2 (0.06) and E1 (0.07)
- Regarding C, D and E axes, there are instances of a high correlation–e.g. C1 and C3, D1 and D2–, and, as a whole, there is an underlying basal correlation between all of these populations. It is likely that all three axes have a common interaction due to a similar location or a similar function.
  – C1 and C3 are specifically related (0.19), and also C2 with C3 (0.16). This may indicate the partial relevance of C3 as a bridge cluster between C1 and C2.
  – C5 cluster is related to C1 (0.14).
  – As observed from UMAPs and graphs too, D1 and D2 are highly related (0.23).
  – Similar with D axis, E1 population overlaps with several populations (C and D axes, as well as A2 and B4), which makes it a good candidate for further analysis.

### Mouse-mouse comparisons

We calculated the Jaccard index with the top 150 markers to perform this comparison, similar to human-human populations. Figure S16 shows the heatmap of Jaccard index values. Overall, there is a high overlap of clusters with each axis and a low overlap of clusters between axes. Also, as expected, bridge clusters show some overlap between their respective clusters.

- z2 and z1 are highly similar (0.25), as expected.
- Surprisingly, contrary to what was observed in UMAPs and PAGA graphs, there is no relevant marker overlap between z1 and y1.
- Populations from axis x have a certain degree of similarity–x2 with x1 (0.15) and x1 with x/y (0.13)–. The bridge axis x/y overlaps with y3 (0.10) and y1 (0.08).
- y axis shows a general overlap across all populations in that axis. The population interactions with a higher degree are between y2 and y3 (0.12), y4 (0.15) and y5 (0.12); and between y4 and y5 (0.11). These observations align with what was previously commented before, based on UMAPs and PAGA graphs.
- Within w axis, there are two pairs of interactions with a high overlap–between w1 and w1 (0.39) and between w3 and w4 (0.17)–. Interestingly, and as it was previously stated, w3 also interacts with other populations, mainly with w/x (0.08) and v1 (0.05). w/x interacts with w3, w4, and with x2 (0.06), but not with w1, which is observed in UMAPs and PAGA graphs.

### Markers of human-mouse comparisons

For each pair of populations, we will show their overlapping markers, divided into four categories: ***markers relevant in human and mouse***, **markers relevant in human**, *markers relevant in mouse* and the rest of markers. We consider markers relevant if they are expressed mainly in that population. This criterion is arbitrary, so we try to make it conservative; we select relevant markers that are only expressed visibly in that population or, at most, in closely similar populations–like A1 and A4–. Genes expressed in more populations or have mild expression are not considered.

- A2: although it shares some markers with some populations from axes y and w, the highest overlap is with population x2. Interestingly, the interaction of A2 with w axis populations (w1 especially), is something that we observe within human-human and mouse-mouse comparisons–A2 with C1/C3 and x2 with w/x–, assuming that C and w are homologous populations. Although there are slight differences between populations, this pattern is recurrent, and could be of interest for further analysis.
  – A2 - x2: CD109, COL3A1, COL5A1, LOXL2, NFATC2, NOTUM, PAM, RSPO3, SCARF2, TCF4, ZNF608, **COL13A1**, **CYP26B1**, **DAAM2**, **ISM1**, **NKD1**, **NKD2**, **PAPPA**, **PREX1**, **PTGS1**, **TGFBI**, **THBD**, **THSD4**, *C10orf105*, *CAV1*, *CCBE1*, *EMX2*, *ENHO*, *LSAMP*, *MAMDC2*, *SPRY1*, *TWIST2*, *WNT5A*, ***AHRR***, ***AXIN2***, ***CD9***, ***COL7A1***, ***F13A1***, ***GREM2***, ***IGFBP2***, ***KCNK2***, ***PTK7***, ***PTPRE***, ***RSPO1***, ***SMIM3***, ***STC1***, ***TNFRSF19*** [11, **12**, *10*, ***14***] - 47
  – A2 - w1: C4orf48, EMB, ETV1, PAPPA, PLCB1, SFRP2, WIF1, WNT5A, **APCDD1**, **AXIN2**, **FGFR2**, **NKD1**, **RAMP3**, **RGS2**, **SMIM3**, *LAMC3*, *NOTUM*, *NRN1*, *RSPO1*, *SPON1*, *TFAP2C*, ***APELA***, ***COL13A1***, ***COL23A1***, ***DAAM2***, ***F5***, ***NKD2***, ***PTK7***, ***RSPO3*** [8, **7**, *6*, ***8***] - 29

Some of these markers, such as *DAAM2*, *NKD2*, *PTK7* or *RSPO3* are coexpressed between A2 and C2 or other C members.

- A1: this population shows a relevant overlap in several populations, although we will focus on x1, x/y, z2 and y3. Among the combinations, the mouse population that best overlaps A1 is z2, with 12/40 relevant markers in mouse and human. Interestingly, none of these markers overlap with relevant markers in the A1-x1 and A1-y3 comparisons, which may imply that A1 population establishes functions that might be of separate fibroblast types in mice.
  – A1 - x1: C1QTNF3, CD34, CHPF, CLEC3B, COL1A1, COL1A2, COL3A1, CREB5, CTSK, DCN, ELN, FKBP9, HTRA1, IGFBP5, ITGBL1, KDELR3, LOXL4, MFAP4, MMP14, P4HA2, PCOLCE2, PDGFRL, PLPP1, RCN3, SLC38A10, SPARC, TSPAN4, VKORC1, **AEBP1**, **ANGPTL1**, **CADM3**, **GDF15**, **GPNMB**, **HPGD**, **MMP2**, **PODN**, **SCARA5**, **TIMP2**, **TNXB**, *ADAMTS2*, *PPIC*, ***CGREF1***, ***CPZ***, ***CYBRD1***, ***CYP4B1***, ***MMP27***, ***SEMA3B*** [28, **11**, *2*, ***6***] - 47
  – A1 - x/y: ADAMTSL1, ANGPTL1, BASP1, CCDC80, CD34, CILP, COL12A1, COL1A1, COL1A2, CTSB, CTSK, DCN, FKBP9, IGSF10, ITGBL1, NUCB2, OGN, P4HA2, MGST1, PCOLCE2, PPIB, RCN3, SERF2, SERPINF1, VKORC1, **ABCC9**, **ADGRD1**, **AEBP1**, **AGTR1**, **CADM3**, **CYP4B1**, **GALNT15**, **HPGD**, **LOX**, **PCOLCE**, **PDGFRL**, **SEMA3B**, **SMOC2**, **SVEP1**, **THBS2**, **THBS3**, **TNXB**, *C1QTNF3*, *GPX3*, *GREB1L*, *PLTP*, ***CGREF1***, ***CPZ***, ***LGR5***, ***MMP27*** [25, **17**, *4*, ***4***] - 50
  – A1 - z2: CREB5, ECM1, FBN1, IGFBP5, IGFBP6, METRNL, PAMR1, PLXDC2, SEMA3C, TIMP3, VGLL3, **CHRDL1**, **CLEC3B**, **MFAP5**, **QPCT**, **SCARA5**, **TIMP2**, *ADAMTSL4*, *BASP1*, *CD248*, *CD34*, *COL14A1*, *DBN1*, *EMILIN2*, *LRRN4CL*, *PTGIS*, *SFRP2*, *UCHL1*, ***ACKR3***, ***ADGRD1***, ***CD55***, ***DPP4***, ***ISLR***, ***LIMS2***, ***MTCL1***, ***NPR1***, ***PI16***, ***PRKG2***, ***SEMA3E***, ***TUBB4A*** [11, **6**, *11*, ***12***] - 40
  – A1 - y3: ANGPTL1, COL12A1, COL14A1, CRYAB, CYB5R3, DHRS3, ELN, FGL2, GAS1, GPX3, IGFBP6, ITGBL1, MFAP4, OGN, PRELP, SERPINF1, SFRP2, TIMP3, **CLU**, **FBLN1**, **FBLN2**, **GALNT15**, **LGR5**, **LOX**, **CD151**, **PCOLCE**, **PDGFRL**, *C1QTNF3*, *CILP*, *MGP*, *PTGIS*, ***ABCC9***, ***OMD***, ***PODN***, ***SMOC2*** [18, **9**, *4*, ***4***] - 35
- A4: the most overlapping mouse populations in mouse are z1 and z2. Interestingly, although y1 shows a degree of overlap comparable to z1 (0.09), it does not have relevant enough markers to be focusing on it. Between z1 and z2, there seems to be more overlap and relevant markers from z2 (0.12) than from z1 (0.09), although many of them are shared–e.g. *AIFL*, *CD248*, *CD55*, *NPR1*, *SEMA3C*, *SEMA3E*, *SFRP4* or *WNT2*–.
  – A4 - z1: ADAMTS5, FSTL1, GFPT2, MGLL, RAMP2, SDK1, TIMP2, TNFAIP6, VASN, ZYX, **ACE**, **CLEC3B**, **GPX3**, **IGFBP6**, **LOXL1**, **MFAP5**, **PTGIS**, **RAB32**, **SCARA5**, *ACKR3*, *ADGRD1*, *AXL*, *CHRDL1*, *FLNC*, *GAP43*, *HAS1*, *HEG1*, *PROCR*, *PRSS23*, *UGDH*, ***AIF1L***, ***CD248***, ***CD55***, ***DBN1***, ***DPP4***, ***EMILIN2***, ***NPR1***, ***SEMA3C***, ***SEMA3E***, ***SFRP4***, ***WNT2*** [10, **9**, *11*, ***11***] - 41
  – A4 - z2: BMP7, CD34, DDAH2, GFPT2, IGFBP5, MGLL, NHSL1, PCSK6, PPP1R14B, PXN, RAMP2, SCARA3, SDK1, SMURF2, TRIO, ZNF385A, ZYX, **ACE**, **ACKR3**, **CLEC3B**, **DBN1**, **FBN1**, **IGFBP6**, **LOXL1**, **MFAP5**, **SCARA5**, **TIMP2**, **TIMP3**, *ACKR2*, *ADAMTSL4*, *ADGRG2*, *AXL*, *CHRDL1*, *DACT2*, *GAP43*, *HEG1*, *PROCR*, *PRSS23*, *UGDH*, ***ADGRD1***, ***AIF1L***, ***CD248***, ***CD55***, ***DPP4***, ***EMILIN2***, ***ISLR***, ***LIMS2***, ***NPR1***, ***PAMR1***, ***PTGIS***, ***RAB32***, ***SEMA3C***, ***SEMA3E***, ***SFRP4***, ***WNT2*** [17, **11**, *11*, ***16***] - 55
  – A4 - y1: ADAMTS5, ADGRD1, AHNAK2, APBB1IP, AXL, CAPG, CDH13, FAP, FSTL1, GFPT2, GLIPR2, HEG1, LGR4, LSP1, MEDAG, MGST1, MMP2, PRSS23, RAMP2, TAGLN2, THBS3, TMSB10, TNFAIP6, UGDH, VCAN, ZNF385A, **CD248**, **CD55**, **CLEC3B**, **EMILIN2**, **FBN1**, **GPX3**, **IGFBP6**, **LOXL1**, **MFAP5**, **TNXB**, **TPPP3**, *HMCN2*, *NOVA1*, *PLTP*, *SSC5D*, ***ACE***, ***CTHRC1*** [26, **11**, *4*, ***2***] - 43
- B1: B1 interacts mainly with z1 population (0.08). Interestingly, z1 also interacts with A4, which might imply that they have similar functions, split in human.
  – z1 - B1: ADAMTS1, BCL3, C3, CSRNP1, CSRP2, GFPT2, HMOX1, IFI16, MYC, NFE2L2, NFKBIZ, NOCT, PHLDA1, PIM1, PLSCR1, PNP, SOD2, **ERRFI1**, **KDM6B**, **MAFF**, **NFKB1**, **NFKBIA**, **NR4A3**, **REL**, **TIPARP**, **TNFAIP3**, **TNFAIP6**, **UAP1**, **ZC3H12A**, *HAS2*, *PTGS2*, *PTX3*, *TNFAIP2*, ***CCL2***, ***CXCL2***, ***FOSL1***, ***GCH1***, ***IL6*** [17, **12**, *4*, ***5***]- 38
- B2 and B3: these two populations interact mainly with y4–0.07 for B3 and 0.08 for B2–. Interestingly, y4 also interacts with B4, and with a higher overlap (0.14), but it is more likely that B2/B3 show relevant markers–3/61 from B4 and 7/38 from B2–.
  – y4 - B2: ABCA8, ABI3BP, C1S, FRMD6, HGF, ID4, IL11RA, LDB2, NRP1, OSMR, P2RY14, PLCXD3, TRIM47, TSPAN11, VEGFA, **APOC1**, **C7**, **CXCL12**, **CYGB**, **GGT5**, **IGFBP3**, **IGFBP7**, *ADCYAP1R1*, *AVPR1A*, *COL4A4*, *NDRG2*, *PTCH2*, *RBP1*, *SNED1*, *TMEM176A*, *TMEM176B*, ***APOE***, ***C3***, ***IL33***, ***MGP***, ***NFIB***, ***SLCO2B1***, ***TNFSF13B*** [15, **7**, *9*, ***7*** ] - 38
  – y4 - B4: ARHGDIB, COL4A2, CYGB, DACT1, EGFR, EPHA3, EPS8, F3, FAM13A, FOXP1, GSN, HGF, IGFBP3, IGFBP7, ITM2B, MGST1, NGFR, NID2, OLFML2B, SERPING1, SRPX, TMEM176B, TNFSF13B, TSHZ2, **ABCA8**, **APOC1**, **APOD**, **C7**, **CXCL12**, **FGF7**, **FMO1**, **FZD4**, **GGT5**, **GPX3**, **HHIP**, **IGF1**, **NFIB**, **NTRK2**, **PODN**, **VIT**, **ZFHX4**, *ABCC9*, *ADAMTSL3*, *APOE*, *BMPER*, *BNC2*, *C3*, *COL4A4*, *FMO2*, *IL33*, *INMT*, *NR2F2*, *NRP1*, *PEAR1*, *PPL*, *PTCH2*, *SFRP4*, *TMEM176A*, ***CHRDL1***, ***GDF10***, ***MGP*** [24, **17**, *17*, ***3***] - 61
- B4: B4 shows a diffuse pattern of overlap throughout y axis (y1 to y5). Although the overlap grade is enough to be considered relevant (0.07 to 0.14), none of the overlaps have enough relevant genes (2 or 3 in most cases). Most of the genes are relevant in human or mouse only, but not in both. Therefore, it is likely that this population is not present in mouse and its putative functions might be shared across the existing mouse populations. Additionally, we observe that y2 and y3 populations also map to populations from A, C, D and E axes, so it is likely that these populations directly cannot be comparable to any human fibroblast population.
  – B4 - y1: APBB1IP, ARPC1B, COL4A2, EBF1, EGFR, GPC3, HIC1, IGFBP3, LGALS3BP, LGMN, MEDAG, NFIB, NR1H3, PRSS23, SLIT2, TGFBR2, THY1, TMEM135, **CXCL12**, **CYGB**, **EFEMP1**, **FMO1**, **FZD4**, **GGT5**, **GPX3**, **IGFBP6**, **ITM2A**, **LSP1**, **MGST1**, **RARRES2**, **ZFHX4**, *BMPER*, *FABP4*, *NOVA1*, *PLAT*, ***PPARG*** [18, **13**, *4*, ***1***] - 36
  – B4 - y2: ARPC1B, CAPN6, COL15A1, COL4A2, COL4A4, CRLF1, EGFR, EPS8, F3, FHL2, HIC1, HSPG2, HTRA3, IGFBP7, ITM2A, ITM2B, NID2, SPRY1, VWA1, **ABCA9**, **APOD**, **CYGB**, **FMO1**, **FZD4**, **GGT5**, **GPX3**, **GSN**, **IGF1**, **LSP1**, **MGP**, **MYOC**, **NFIB**, **PODN**, **TSHZ2**, **ZFHX4**, *ADAMTSL3*, *BMPER*, *INMT*, *PEAR1*, ***ABCA8*** [19, **16**, *4*, ***1***] - 40
  – B4 - y3: ARHGDIB, DEPTOR, DHRS3, EPHA3, F3, FHL2, GHR, GPC3, IGFBP3, IGFBP6, IGFBP7, KCNJ8, NPY1R, PEAR1, SFRP4, SLIT2, SUSD2, TIMP3, **ABCA8**, **APOD**, **CYGB**, **FGF7**, **FMO1**, **GPX3**, **GSN**, **IGF1**, **ITM2A**, **MGP**, **MYOC**, **NFIB**, **PHLDA3**, **PODN**, **TXNIP**, **ZFHX4**, *ABCC9*, *FMO2* [18, 16, 2, 0] - 36
  – B4 - y4: ARHGDIB, COL4A2, CYGB, DACT1, EGFR, EPHA3, EPS8, F3, FAM13A, FOXP1, GSN, HGF, IGFBP3, IGFBP7, ITM2B, MGST1, NGFR, NID2, OLFML2B, SERPING1, SRPX, TMEM176B, TNFSF13B, TSHZ2, **ABCA8**, **APOC1**, **APOD**, **C7**, **CXCL12**, **FGF7**, **FMO1**, **FZD4**, **GGT5**, **GPX3**, **HHIP**, **IGF1**, **NFIB**, **NTRK2**, **PODN**, **VIT**, **ZFHX4**, *ABCC9*, *ADAMTSL3*, *APOE*, *BMPER*, *BNC2*, *C3*, COL4A4, *FMO2*, *IL33*, *INMT*, *NR2F2*, *NRP1*, *PEAR1*, *PPL*, *PTCH2*, *SFRP4*, *TMEM176A*, ***CHRDL1***, ***GDF10***, ***MGP*** [24, **17**, *17*, ***3***] - 61
  – B4 - y5: ABCA8, F3, GPC3, HSPG2, IL33, ITM2A, LXN, MGP, RARRES1, **COL15A1**, **GSN**, **NTRK2**, *ABCA6*, *APOD*, *FOXS1*, *NR2F2*, *PHGDH*, *VIT*, *VWA1*, ***MYOC***, ***TSHZ2*** [9, **3**, *7*, ***2***] - 21
- C2 population is related to several mouse populations, but mainly w/x, with whom it shares 12/42 relevant markers–based on an overlap value of 0.09–. Secondly, w1/w2 populations share 6/34 relevant markers–overlap of 0.06–. Lastly, y3 only shows overlap with 2 relevant markers, despite showing a similar overlap with w1/w2–0.05–. Therefore, we do not consider the C2-y3 interaction as useful.
  – C2 - w/x: ADAMTS9, CBFA2T3, COL11A1, EDNRA, ENHO, HTRA1, SLC48A1, SRPX, TBXA2R, TENM3, **CPNE5**, **CRABP1**, **FZD1**, **GPM6B**, **MEOX2**, **NCAM1**, **NFATC2**, **PLXDC1**, **PTH1R**, **RSPO4**, **SLC40A1**, **TBX15**, **TCF4**, **TRIB2**, **TRPS1**, *HS3ST6*, *KIF26B*, *MEGF6*, *NR2F1*, *TSHZ3*, ***CHST15***, ***COCH***, ***CYP1B1***, ***DKK2***, ***EMID1***, ***FIBIN***, ***FMOD***, ***MAFB***, ***MKX***, ***NRP2***, ***PTGFR***, ***TNMD*** [10, **15**, *5*, ***12***] - 42
  – C2 - w1|w2: CNTN1, CTTNBP2, PRLR, PTGER3, PTPRD, SFRP1, TSPAN7, **ATP1B1**, **BTBD11**, **CLEC14A**, **DKK2**, **EMB**, **GPM6B**, **IGFBP5**, **MEIS2**, **NCAM1**, **SPARCL1**, *ALX4*, *CRABP2*, *DUSP10*, *LEF1*, *NDP*, *PDE1A*, *PTK7*, *RSPO4*, *RUNX3*, *SDC1*, *WNT5A*, ***CRABP1***, ***DAAM2***, ***NDNF***, ***NOTUM***, ***TRPM3***, ***TRPS1*** [7, **10**, *11*, ***6***] - 34
  – C2 - y3: CALM2, MDFIC, OGN, PDE4B, PMEPA1, PRELP, RERG, SERTAD4, SUSD2, **EMB**, **FIBIN**, **FMOD**, **FZD1**, **KCNAB1**, **PTPRD**, **PTPRK**, **TBX15**, *CAVIN2*, *F3*, *FGF9*, *LTBP4*, *PAQR6*, *SFRP1*, ***ASPN***, ***CADM1*** [9, **8**, *6*, ***2***] - 25
- C1 population interacts mainly with w4 and w3 mouse populations–0.12 and 0.11 overlap respectively–. When looking at the population marker overlap, we observe a much larger overlap of relevant markers with w4–18/53– than with w3–2/48–.
  – C1 - w4: ARHGAP28, ATP10A, CDH11, COL27A1, COL4A1, KIAA1217, KLF5, LMO7, MGLL, NPNT, NUAK1, NXN, PALLD, PMEPA1, POSTN, RNF152, SPARC, **CNN2**, **MME**, **SEMA5A**, *ADAMTS6*, *CD200*, *COL7A1*, *FOXD2*, *LMO4*, *LRRC15*, *NID2*, *NTRK3*, *PARD6G*, *PAWR*, *PRR5L*, *SATB2*, *TMEM119*, *TNMD*, *TPM2*, ***ACAN***, ***ACTA2***, ***ADAMTS18***, ***ADAMTS9***, ***BCL11B***, ***CALD1***, ***CCND1***, ***COL11A1***, ***COL12A1***, ***COL8A2***, ***CPXM2***, ***EDNRA***, ***EDNRB***, ***EGFL6***, ***MEF2C***, ***RAMP1***, ***TAGLN***, ***TENM3*** [17, **3**, *15*, ***18***] - 53
  – C1 - w3: AFAP1L2, COL6A3, FNBP1L, FOXD2, JAG1, KIF26B, LMO4, MDK, MGLL, MICAL2, NPM1, PALLD, PARD6G, PAWR, PMEPA1, SHOX2, SOX18, SOX4, TMEM119, TNS3, TPM2, **ACAN**, **ACTA2**, **ADAMTS18**, **ALX4**, **BCL11B**, **CALD1**, **CD200**, **CDH11**, **CNN2**, **COL11A1**, **DKK3**, **EDNRA**, **EVA1A**, **F2R**, **KIAA1217**, **LAMC3**, **MEF2C**, **PTCH1**, **ROBO2**, **RUNX2**, **STMN1**, *LRRC15*, *MMP11*, *NTRK3*, *TAGLN*, ***CCDC3***, ***EGFL6*** [21, **21**, *4*, ***2***] - 48
- C3 population, being established as a putative “bridge” between C1 and C2, is expected to overlap with mouse populations with C1 and C2. As such, C3 overlaps with w/x–0.11–and w4–0.10–populations, as expected. Looking at the significative markers, both w/x and w4 show a slight overlap–4/51 and 5/44– with C3. Of note, the number of relevant markers of C3 is reduced; therefore, we would expect this lower level of relevant marker overlap between human and mouse. Additionally, C3 also shows some degree of overlap with x1–0.08–, but there is only 1 relevant marker between human and mouse, so this comparison should not be considered.
  – C3 - w/x: ADAMTS9, AQP1, CDH11, CHCHD10, CNN2, COL11A1, COL16A1, CRABP1, DKK3, EDNRA, EGFLAM, EMP2, FIBIN, FZD1, GPM6B, HMCN1, MEF2C, MFAP4, MMP11, MXRA8, PLXDC1, PTMA, RSPO4, SESN3, SLC40A1, TBX15, TCF4, THY1, TMEM119, TMEM204, TPM2, TRPS1, **F2R**, **HTRA1**, **MFAP2**, **POSTN**, **TENM3**, *CPXM2*, *DKK2*, *EMID1*, *KIF26B*, *MEGF6*, *NR2F1*, *NREP*, *NRP2*, *RFLNB*, *TSHZ3*, ***COL7A1***, ***MAFB***, ***MMP16***, ***RASL11B*** [32, **5**, *10*, ***4***] - 51
  – C3 - w4: BGN, BHLHE41, CALD1, CDH11, CNN2, EMP2, FABP5, ITGB1, ITGBL1, MGLL, MYH9, NPNT, PALLD, PPIC, PRELP, SORCS2, SPARC, **KIAA1217**, **LMO7**, **PMEPA1**, **SEMA5A**, *ACAN*, *ACTN1*, *ADAMTS9*, *AQP1*, *CGNL1*, *COL11A1*, *COL12A1*, *CPXM2*, *EDNRA*, *EGFLAM*, *LBH*, *MEF2C*, *MME*, *SLC40A1*, *TAGLN*, *THBS4*, *TMEM119*, *TPM2*, ***COL7A1***, ***COL8A2***, ***LRRC15***, ***POSTN***, ***TENM3*** [17, **4**, *18*, ***5***] - 44
  – C3 - x1: BMP1, CHPF, COL3A1, COPZ2, GLT8D2, ITGBL1, LUM, MARVELD1, MFAP4, MMP14, MMP2, P4HA2, PYCR1, RCN3, RRBP1, SLC16A3, SPARC, SPHK1, SPON2, **BGN**, **C1QTNF6**, **COL5A1**, **COL5A2**, **ELN**, **HTRA1**, *ADAMTS2*, *AEBP1*, *COL16A1*, *COL1A1*, *COL1A2*, *EMID1*, *PPIC*, *PPP1R14A*, *SULF2*, **LTBP2** [19, **6**, *9*, ***1***] - 35
- C5 population shows some overlap with population w3–0.08–and w1|w2–0.07–. When looking at the relevant markers, we observe a better overlap with w1|w2 than with w3–8/29 vs 2/26–. Interestingly, the amount of markers overlapping between C2 and w1|w2 is similar to the ones with C5 and w1|w2, although higher in proportion–6/34 vs 8/29–; and the markers that are relevant in human and mouse between the two comparisons do not overlap between them. Therefore, C2 and C5 functions may be shared between w1 and w2, without a clear pattern.
  – C5 - w3: CFL1, COL11A1, DKK3, EDNRA, EEF1A1, F2R, PFN2, PMEPA1, PTMA, SMS, SOX4, STMN1, **ALX4**, **CXCR4**, **HEY2**, **JAG1**, **LMO4**, **MARCKSL1**, **PRDM1**, **PREX2**, **PTCH1**, **SOX18**, *KIF26B*, *LAMC3*, ***CDH11***, ***ROBO2*** [12, **10**, *2*, ***2***] – 26 C5 - w1|w2: C4orf48, FST, LAMC3, MARCKSL1, PLK2, PTCH1, PTMA, RAB34, SPARCL1, **CXCR4**, **IGFBP3**, **PDE3A**, **PRDM1**, **PREX2**, **ROBO1**, *EDN3*, *HEY2*, *SDC1*, *SLC26A7*, *SPON1*, *TRPS1*, ***ALX4***, ***BMP7***, ***INHBA***, ***LEF1***, ***PGM2L1***, ***SOX18***, ***TFAP2A***, ***WNT5A*** [9, **6**, *6*, ***8***] - 29
- D1 vs D2 - y5
  – D1 - y5: ARL4A, CNN3, CSPG4, ITM2A, LUM, NID1, **EBF2**, **ENTPD2**, **ETV1**, **GPC3**, **PHLDA1**, **SPARCL1**, **TM4SF1**, *CDKN2B*, *MATN2*, ***ABCA8***, ***APOD***, ***COL8A1***, ***FOXS1***, ***NR2F2***, ***P2RY14***, ***SOX9***, ***TGFBI***, ***VIT*** [6, **7**, *2*, ***9***] - 24
  – D2 - y5: ABCA8, CNN3, EBF2, ETV1, ITM2A, NR2F2, SMOC2, TGFBI, TM4SF1, VIT, *FOXS1*, ***MATN2***, ***P2RY14*** [10, **0**, *1*, ***2***] - 13
- D1 vs D2 - v1
  – D1 - v1: CAVIN2, CSPG4, CSRP1, CTNNAL1, DOCK9, FRMD4B, JAG1, MTUS1, NDRG1, PEAR1, PLEKHA4, PTCH1, STMN1, SYNE2, TLN2, TUBA4A, **CD200**, **DUSP5**, **EFNA1**, **EGR3**, **ETV1**, **MRAS**, **NR2F2**, **OLFML2A**, **SLC12A2**, **SOX9**, **TENM2**, **TGFBI**, **VIT**, *BNC2*, *EZR*, *ITGA6*, *ITGB4*, *KLF5*, *SBSPON*, *SLC2A1*, ***AKAP12***, ***CLDN1***, ***EBF2***, ***EFNB1***, ***ETV4***, ***MTSS1***, ***NDRG2*** [16, **13**, *7*, ***7*** ] - 43
  – D2 - v1: v1 - D2: BHLHE40, DMD, ENDOD1, ETV1, FRMD4B, JAG1, KTN1, MDFIC, MFAP5, NDRG1, NR2F2, PHLDA3, PLSCR4, PTCH1, SFRP1, STMN1, STXBP6, SYNE2, TLN2, **ADAMTSL3**, **AKAP12**, **CAVIN2**, **CSRP1**, **DACT1**, **DUSP5**, **IGFBP6**, **ISYNA1**, **MRAS**, **OLFML2A**, **PEAR1**, **PLEKHA4**, **SORBS1**, **TGFBI**, **TJP1**, **VIT**, *CCDC3*, *DDIT4*, *EZR*, *PERP*, *TPD52*, *TRIB2*, ***AQP3***, ***BNC2***, ***CAV1***, ***CAV2***, ***CLDN1***, ***DOCK9***, ***EBF2***, ***EFNB1***, ***GAB1***, ***GPC1***, ***ITGA6***, ***ITGB4***, ***KLF5***, ***KRT19***, ***MTSS1***, ***NDRG2***, ***SBSPON***, ***SLC2A1***, ***TENM2*** [19, **16**, *6*, ***19***] - 60
- D1 vs D2 - y4
  – D1 - y4: APOD, BNC2, EBF2, EGFR, EPS8, FMO1, IGFBP7, ITGA6, KLF5, LTBP4, MEOX2, NDRG2, NGFR, NR2F2, PEAR1, S100B, SCN7A, VIT, **CYP1B1**, **P2RY14**, **PHLDA1**, **SPARCL1**, **WFDC1**, *ABCA8*, *CYGB*, *INMT*, *SFRP4*, ***ENTPD2***, ***FMO2*** [18, **5**, *4*, ***2***] - 29
  – D2 - y4: y4 - D2: ABCA8, ADAMTSL3, CCDC3, COL4A2, EBF2, EGFR, CYP1B1, IGFBP7, ITM2B, LTBP4, MEOX2, MFAP5, NDRG2, NR2F1, NR2F2, P2RY14, SCN7A, VIT, **BNC2**, **CAV1**, **DACT1**, **GPC6**, **ITGA6**, **KLF5**, **NGFR**, **PEAR1**, *CHRDL1*, *SFRP4*, ***INMT*** [18, **8**, *2*, ***1***] - 29
- E1 population shows more reduced levels of overlap–0.05 to 0.07– with several mouse populations: y2, y3, y4, w3, w4, and v1. Some of these populations, like w3, w4, and v1, have a high chance of being related to other human populations like C1 or D2, and it is less likely, based on the smaller overlaps of relevant markers, to be associated with E1. On the other hand, based on the heatmap from Figure 4, and on the overlapping markers, populations y2, y3 and y4 are not one-to-one correspondence candidates, since they have marker overlap with a wide range of human populations. Therefore, based on this analysis, we cannot assign a clear human-to-mouse comparison candidate to E1.
  – E1 - y2: APOD, CAPG, COL15A1, FHL2, G0S2, GSN, IGF1, ITGA11, ITM2A, ITM2B, JUP, LAMA2, LPL, LSP1, MFAP5, SCN7A, SFRP1, SPRY1, TMEM204, **CMKLR1**, **MGP**, **PLEKHA6**, **RGMA**, *COL14A1*, *CRLF1*, *GPM6B*, *INMT*, *PEAR1*, *TGFBI*, *TSHZ2*, *VWA1*, ***MEOX1*** [19, **4**, *8*, ***1***] - 32
  – E1 - w3|w4: ACOT7, ALX4, BHLHE40, BHLHE41, C1QTNF7, CALD1, CCDC34, CDC42EP3, CDK6, DKK3, EFNB1, EMP2, EVA1A, FHL2, HES1, HEY1, ITGB1, JUP, KLF5, LAMC3, LMO7, MAF, MFGE8, MSI2, RASGRP2, RGCC, RUNX3, SEMA3G, SEMA5A, SHOX2, TBX18, **EGR2**, **KIAA1217**, **OLFML2A**, **RAMP1**, **TCF7L2**, *AQP1*, *ARHGDIB*, *CDH11*, *COL8A2*, *CSRP1*, *EDNRA*, *F2R*, *FOXD2*, *PARD6G*, *RUNX2*, *STMN2*, ***IGFBP2***, ***NTRK3*** [31, **5**, *11*, ***2***] - 49

### Supplementary Discussion - Putative function of fibroblasts

In order to elaborate on the putative functions of the fibroblasts, we used markers from previous results, as well as markers manually curated derived from DEGs. Then most of the markers were elaborated using information from GeneCards, and certain selected markers, further described throughout this section, were studied by searching information in the literature.

To perform this search, the following formulas were used in Pubmed and Google Scholar: GENE AND (fibro*) AND (skin OR derm*). Additional search terms including immune, nerve, vasc*, TGF, Wnt, ECM were used depending on the scope. All the information derived from the bibliographic research is available at the following spreadsheet.

### C axis

***C1.*** C1 population can be described by three main markers used in other single-cell publications: *ACTA2*, *COL11A1* and *DPEP1*. This population, which shares these markers with mouse w4 population, as well as *EDNRA* and *TENM3*, is described by Ahlers et al. (2022)–*COL11A1*^+^ *DPEP1*^+^–and Tabib et al. (2021)–*COL11A1*^+^ *ACTA2*^+^–as DS cells. Some of these markers, as well as markers described below, are represented in Figure S17.

One of the most relevant markers studied from C1 populations is *ACTA2*, the gene encoding for α-SMA. *ACTA2* is already described as a marker of DS cells (Mistriotis and Andreadis, 2013), and is traditionally described as a myofibroblast marker. In fact, during wound healing, *ACTA2*^+^ fibroblasts also express *COL1A1* and *ITGB1*, although neither of them is strictly required for proper wound closure (McAndrews et al., 2022). Another report indicates that, although *ACTA2*^+^ myofibroblasts contribute to wound closure, they are not strictly necessary, and other actins (*ACTB*, *ACTG1*) may also be involved in the process (Ibrahim et al., 2015). Additionally, at least in myocardial fibroblasts, they may still differentiate into myofibroblasts regardless of ACTA2 expression (Li et al., 2022b). All these studies may be relevant to postulate whether *ACTA2*^+^ DS fibroblasts already exist in a primed pro-fibrotic state, or if myofibroblasts already originate from another population.

A set of markers relevant to DS expression are related to its location. Some markers descriptive of this population are also expressed in the BM surrounding the CTS, such as *KRT17* (Antonini et al., 2013), *COL5A1/2* (Chanoki et al., 1988), MME (Morisaki et al., 2013), and *MDK* (Woo et al., 2022). Interestingly, some of these markers are also detected in DP as well as in DS: *ADAMTS18* (Hagner et al., 2020), *COL7A1* (Tsutsui et al., 2021), *COL12A1* (Sasaki and Enami, 1996), *LEF1* (Sun et al., 2022), *MDK* (Rendl et al., 2005).

C1 has a clear link to ECM. For instance, *TNMD* is expressed probably to control the width of collagen fibres (Docheva et al., 2005); and *ACAN*, *ASPN*, *COL11A1* and also *TNMD* are usually expressed in cartilage, tendon and ligaments (Roughley and Mort, 2014; Kalamajski et al., 2009; Luo et al., 2019; Sun et al., 2019).

C1 may also be related to the blood vessels surrounding the CTS, due to the expression of COL15A1 (Manon-Jensen et al., 2019) and COL21A1 (Kehlet et al., 2019a), which are usually found surrounding blood vessels. Additionally, C1 also expresses *EDNRA*, which regulates blood pressure and blood vessel constriction (Liu et al., 2019), and which, in individuals with mutations in this gene, may lead to alopecia (Gordon et al., 2015); indicating the putative relevance of this gene, and C1 population in general, in HF vasculature. This is schematised in Figure S19 .

Also related to HF homeostasis, C1 expresses genes that are related to hair cycle regulation. For instance, *MME* is more active in early anagen (Morisaki et al., 2013); KRT17 binds CDKN1B to prolong the anagen phase (Tong and Coulombe, 2006; Panteleyev et al., 1997); and *LEF1* is associated with lengthening of anagen and shortening of telogen and catagen (Zhang et al., 2013).

Therefore C1 DS cells that are active in the anagen phase express several genes contributing to that state; also express HF-related specific ECM genes, and have a wide control of CTS microvessels, probably to supply the IRS and ORS cells within the HF.

***C2.*** C2 population shares common robust markers both with mouse w/x–*CHST15*, *COCH*, *DKK2*, *FMOD*, *POSTN*–and w1/w2–*CRABP1*, *DAAM2*, *NOTUM*, *TRPM3*, *TRPS1*–populations. One of these markers, *CRABP1*, is a classical DP marker described by Collins and Watt (2008). Additionally, Ahlers et al. (2022) described a DP population expressing both *COCH* and *CRABP1*, C2 population exclusive markers. Some of these markers, as well as markers described below, are represented in Figure S17.

One of the most relevant markers of C2 is COL24A1, an specific collagen of this population. Although its function is not well known, it has been observed to be highly expressed in bone and has 3 non-collagenous domains that may confer additional, *non-traditional*, functions (Nielsen and Karsdal, 2019). The relationship of this collagen with bone is relevant since we observe that C2 shows highly active participation within the ECM homeostasis, and expressed other markers, like *ASPN*, which has been shown to actively participate in calcium binding of collagen to induce ECM mineralisation (Kalamajski et al., 2009).

Other relevant ECM regulatory components are *CHADL*, which regulates collagen and aggrecan (*ACAN*) fibrillogenesis (Tillgren et al., 2015), *F13A1*, which binds fibronectin and vitronectin (Winnemöller et al., 1991; Huttenlocher et al., 1996); *P3H2*, which binds collagens I and IV and is necessary for their maturation (Tiainen et al., 2008); and *SDC1*, a promiscuous HSPG that binds collagens I, II and IV, fibrin, fibrillin, TNC and vitronectin, as well as MMPs, ADAMTSs, BMPs, EGFs, cytokines and complement proteins (Stepp et al., 2015). Other relevant ECM modulators expressed by C2 are *ADAMTS9* and *CHST15*, which degrade CSPGs such as aggrecan (*ACAN*) and versican (*VCAN*) (Kelwick et al., 2015; Kai et al., 2017).

This ECM regulatory activity, as well as the active expression of ECM components that are more “cartilage-like”, similar to markers from C1 DS population, may confer C2 a unique ECM microenvironment necessary for DP cells to dwell within the HF.

One of the hallmarks of HF signaling is the Wnt pathway. From the analysis of markers, we observe that C2 shows a high Wnt activity. Interestingly, this signaling is *mixed*, that is, both pro- and inhibitory markers are expressed. For instance, *DAAM2* enhances Wnt activity by receptor aggregation, which potentiates the downstream cascade (Lee and Deneen, 2012; Lee et al., 2015), but also expresses *NOTUM*, which inhibits this aggregation (Zhang et al., 2015). C1 population also expresses *FZD1* and *RSPO4*, receptor and co-activator of canonical signaling (Gazit et al., 1999; Szenker-Ravi et al., 2018); together with *DKK2* and *PTK7*, which inhibit canonic signaling (Ahn et al., 2011; Lhoumeau et al., 2011). PTK7, additionally, is a non-canonic signaling co-receptor, which may interact with C1-expressed WNT10, a non-canonic ligand (Chen et al., 2012).

Wnt signaling in DP was observed to be mainly pro-anagenic since many of these markers are not expressed during telogen phase (Reddy et al., 2001; Lim and Nusse, 2012). Nonetheless, some markers like *SFRP1*, also observed in cuboidal ORS at the base of the HF, are potentially active in regressing telogenic HFs (Geyfman et al., 2014).

***C3.*** C3 population, based on the PAGA graphs and other population analyses, is highly related to C1 and C2, since many markers expressed by C3 are also expressed by C1 and C2, but not vice-versa. Based on human-mouse population similarity, C3 is most related with w/x and w4, with shared markers such as *COL7A1*, *MMP16*, *LRRC15* and *POSTN*. w4 is also similar to C1, so probably C3 is more related to C1 than C2. Some of these markers, as well as markers described below, are represented in Figure S18.

*TNMD*, expressed by all C populations–Ahlers et al. (2022) classifies it as a DS marker–, reduces the amount of *BGN* (biglycan), *COMP* and *FN1* (fibronectin). Therefore, C3 may be, at least ECM-wise, a spatially complementary population to C1 and C2, with a separate ECM niche (Lin et al., 2017).

Despite being a “mixture” of C1 and C2, C3 expresses certain genes more than the other two populations. Some of these genes, such as *MMP11*, *BGN*, or *POSTN*, are tightly related to the functioning of the ECM. *BGN* is an SLRP that binds collagens I and VI, chondroitin and dermatan sulfate (Roughley and White, 1989; Schönherr et al., 1995); and *POSTN*, localised around the HF, binds several components, including collagen I, fibronectin and tenascin C (Yamaguchi, 2014). Interestingly, Deng et al. (2021) observed a *POSTN*^+^ population in keloid samples, which increases collagen production (Zhang et al., 2014); and the authors show that blockade of POSTN function led to a decrease in collagen production. Therefore, C3 might be probably related to ECM homeostasis of HF cells.

Regarding its location, one of the few exclusive markers expressed by C3 is *KLK4*, a member of the kallikrein serine proteases family. It does not have a clearly defined function, although it may participate in certain immune interactions (Filippou et al., 2020), androgens may regulate its expression in certain cancers (Korkmaz et al., 2001), and may also regulate the degradation of ECM components (Obiezu et al., 2006). Regarding its location, KLK4 has been found in IRS, but also in ORS, SG and eccrine glands; similar to other KLKs such as KLK6/10/11 (Komatsu et al., 2003), which are described in the literature but are not expressed in our analysis.

Based on the expression of these markers, the population C3 may be involved in both the secretion of HF-specific ECM, as well as in its degradation, based on the expression of specific metalloproteases and kallikreins.

***C5.*** Similar to C3, C5 function is not a cell type categorised based on bibliographic records. Moreover, the fact that, compared to the rest of C axis populations, has comparatively fewer cells, indicates that it is probably a cell population not present in all HF stages, or that it represents a specific cell type with a reduced number of cells, like stem cells. Some of these markers, as well as markers described below, are represented in Figure S18.

Looking at the PAGA graphs in human, C5 shares a higher transcriptomic similarity with C1 DS compared to C2 DP, coexpressing genes such as *CDH11*, *INHBA*, *KRT17*, *LEF1*, *MME*, *SOX18*. However, it also shares specific key genes with C2 DP population–*CRABP1*, *PTK7*, *DAAM2*, *CYP26B1*–. Regarding mouse-human similarity, C5 is transcriptomically more similar to w1/w2 populations, with shared genes such as *BMP7*, *INHBA*, *LEF1*, *SOX18*, *WNT5A*; mostly expressed by C1.

Some of these markers, apart from being associated by similarity, are reported to be expressed by DP cells, like *WNT5A* (Lim and Nusse, 2012), *CRABP1* (Collins and Watt, 2008), *TPD52* (Rubin et al., 2004), *CXCR4* (Zheng et al., 2022), *LEF1* (Sun et al., 2022), or *SOX18* (Villani et al., 2017). Another relevant marker, *RSPO3*, has been described in DP cells to induce proliferation of HF stem cells and DS (Hagner et al., 2020). In fact, considering that some of these markers are also expressed in other parts of the HF, like *LEF1* in DS, *WNT5A* in IRS (Lim and Nusse, 2012), *CXCR4* in ORS (Zheng et al., 2022), *KRT17* in ORS and SG (Antonini et al., 2013) and *MME* in bulge DS (Morisaki et al., 2013), it is possible that the expression of certain DP markers in additional cell types is due to C5 population being part of these cell types.

Despite differences and similarities with other HF populations, C5 expresses a few exclusive markers, such as *TPD52*, *KRT9*, *WNT5A*. There are other markers which do not have a clear role, either in general or in HF, such as *LMO3*, *LUZP2* or *PKP4*. This last one is expressed in desmosomal plaques (Berika and Garrod, 2014).

Similarly to C1, C5 shows a high and, at the same time, diffuse Wnt activity. For instance, *APCDD1* and *NKD1* expression is linked to negative Wnt signaling (Mazzoni et al., 2017; Angonin and Raay, 2013), whereas *DAAM2* is an activator (Lee and Deneen, 2012), and *RSPO3* and *DKK2* are canonical Wnt activators and inhibitors respectively (Szenker-Ravi et al., 2018; Ahn et al., 2011). Lastly, *WNT5A* is a non-canonical Wnt ligand (Romanowska et al., 2009), which can be expressed by *SOX18*–another marker of C5–(Villani et al., 2017), and WNT5A induces the expression of *KRT9* (Rinn et al., 2008); all 3 markers are expressed by C5. Surprisingly, *KRT9* is a classical palmoplantar keratin (Fu et al., 2014), with no bibliographic records indicating its expression if HF and other skin appendages. This process is schematised in Figure S19.

Similar to C1, there are certain markers involved in cell cycle regulation, directly or indirectly, such as *IRF1*, which regulates *CDKN1A* (Dornan et al., 2004); as well as in HF cycle, like *MME*, which shows increased activity in anagen (Morisaki et al., 2013), or *CXCR4* which, besides its classic CXCL12 receptor involved in chemotaxis, delays the transition from telogen to anagen (Zheng et al., 2022).

Another function of C5, not described previously in C1 or C2, is TGF-β signaling. Two of the markers of C5 are BMP7, a key ligand of the non-canonical TGF-β pathway (Meng et al., 2016); and INHBA, a common TGF-β inhibitor that binds to the ligand and inhibits receptor coupling (Ferdous et al., 2007). It is possible that both genes act synergistically to promote only non-canonical TGF-β signaling.

Although it might not be its main function, C5 expresses *S100B* and *SLC5A3*, two key genes in neurogenesis and vasculogenesis. *S100B*, also expressed by D1/D2, is a classical neuron marker, although it is also present in melanocytes, adipocytes, DCs and lymphocytes (Donato et al., 2009). S100B has been found to participate in neuron myelinisation (Fujiwara et al., 2014), neurite growth (Donato et al., 2009), satellite cell activation (Sorci, 2013) and production of iNOS and VEGF by endothelial cells (Donato et al., 2009). On the other hand, *SLC5A3*, an inositol-Na^+^ transporter expressed in nerve, pancreas, lung and muscle (Ma et al., 2012); besides being a key cellular and intercellular osmolarity regulator (Pizzagalli et al., 2020), it forms complexes with other channels in nerve and blood vessels to sense neuronal excitability (Dai et al., 2016) and vessel contractibility (Barrese et al., 2020).

Lastly, an interesting remark about C5 population is the expression of certain B axis markers, like *CD74* (B3), *IER1* (B1) and *IRF1* (B1 and B3). The functions of these genes will be discussed later but, at first sight, it is possible that either the C5 population carries some level of immune response or these genes have additional functions apart from the ones exerted by immune cells and immune-related fibroblasts.

Therefore, based on transcriptomic similarity, markers and putative functions, there are several hypotheses regarding the cell type that C5 population could be assigned to. It is possible that C5 represents a population that shares its location with the DS, but also lies within the DP environment. Firstly, C5 cells could either be matrix cells adjacent to DP, implicated in HF proliferation, or precortex cells, implicated in differentiation. However, both cell types have a marked expression of *KRT15* o *KRT6* (Geyfman et al., 2014), which are not specifically expressed within the fibroblast cells in our analysis. Secondly, C5 cells could also be dermal cup cells, that is, DS cells located below the dermal papilla. Hagner et al. (2020) analyse dermal cup cells in mice and show that these cells express specifically *Epha3*, *Hic1*, *Igfbp2* and *Tnnt1*, *Mcam1* and *Pcp4* come of which are expressed by E1 or C1 sparsely–*Epha3*, *Igfbp2*, *Tnnt1*–and the rest of genes do not show any expression pattern. This effect is already expected due to the gene expression differences that are commonly observed across species. Lastly, C5 population is a candidate of telogenic DP or DS cells. Considering that the HF cycle is asynchronous in humans and the fact that around 5% of HF cells are found to telogen (Serrano-Falcón et al., 2013), the reduced amount of C5 cells may be an indicator. Additionally, despite telogen being popularly considered as an inactive HF state, in reality, telogen DP is very active and necessary to maintain the club hair properly (Geyfman et al., 2014). The main drawback to this assumption is that C5 population expresses both pro-anagenic and pro-telogenic markers, and the number of markers expressed is reduced, which decreases the conclusiveness of the results.

#### Consistency of C axis populations

Results from this analysis, as well as other studies that characterise HF cells using scRNAseq show that the HF is an incredibly heterogeneous structure and, focusing on fibroblasts, this analysis shows that, besides DP and DS cells, other new fibroblastic populations are relevant for HF biology, and are yet to be characterised (Chovatiya et al., 2021; Wu et al., 2022). Even more, the characterisation of DP and DS, at least based on single-cell and other studies, is yet to be 100% defined as well.

In fact, not only differences between human and mouse markers are to blame, but also the differences across studies. For instance, the comparison of DS and DP populations described by Shin et al. (2020) and Joost et al. (2020) described in Table S3 shows that the overlap of markers is still extremely small. For instance, we see that CTS1 from Shin et al. (2020) is more similar to DS2 from Joost et al. (2020), although the Jaccard index is only 11.11% (10 shared out of 99 compared between both datasets); and CTS2 is more similar to DS1, but the similarity to DS2 cannot be discarded–with Jaccard indexes of 12.36 (11/89) and 7.53 (7/93)–. Regarding DP populations, the only “relevant” overlap is between aDP and DP3–Jaccard index of 12.36 (11/89)–.

The same effect occurs comparing Shin et al. (2020) or Joost et al. (2020) markers to the merged populations from this analysis. Both tDP and aDP populations reported by Joost et al. (2020) match w1 and w2 indistinctively, and DS1 and DS2 populations match w3 and w4. However, the differences between w1/w2 and w3/w4 populations, despite being highly similar, are consistent across datasets, which implies that the characterisation of individual datasets may not be consistent and that, at least in mouse, annotation of w axis does not reflect differences between anagen and telogen states. However, considering the small number of cells within Joost et al. (2020) population, it is possible that anagen/telogen differentiation was not relevant enough to be considered for the marker-to-population algorithm. If further studies including more anagen/telogen cells are introduced in the analysis, probably new w populations reflecting this heterogeneity will appear.

A similar phenomenon can be described for the mouse population w5. This mouse population, characterised by its increased expression of cell cycle markers–e.g. *Mki67*, *Cdk1*, *Cdkn3*–is described by Shin et al. (2020) as HF dermal stem cells (hfDSCs). However, although this population is reported to exist in human HF (Wu et al., 2022), based on the expression of *Col17a1* and *Krt15*, it is not located within the fibroblasts clusters. Therefore, it is possible that hfDSCs do not share the same lineage in both organisms.

With all this in mind, even a *proper* mouse HF fibroblast characterisation may not lead to new human populations, firstly, due to the fundamental limitations between human-mouse comparisons; and secondly, due to the fact that, even within the same organism, fundamental differences between mRNA and protein levels arise. Tsutsui et al. (2021) comment that several genes, including *Col4a1*, *Tnc* or *Smoc1* show significative differences between expression levels of mRNA and protein. They state that these differences may be due to posttranslational mechanisms or due to protein expression by adjacent populations affecting the measurement levels.

This last effect is also sound in human samples. In fact, many of the fibroblast markers are also expressed by many cell types, including keratinocytes, endothelial cells, perivascular cells and immune cells; so the putative fibroblast functions inferred during this discussion section may be affected by additional interactions with adjacent populations.

### A axis

#### A2

A2 population is one of the key archetypes of the A axis, together with A1, and shows both a differentiated marker profile and location. A2 population is highly likely to be associated with the papillary dermis, which we will extend on this section.

#### ECM production and modulation

A2 population shows a rich expression of collagens that are known to be bound to DEJ structures, located in BM around vessels or other structures, or are located in the papillary dermis. The most relevant collagens in this aspect are the following: (1) *COL7A1* is a component of the anchoring fibrils in the BM, which bind collagen I and III fibres (Barbieri et al., 2014); (2) *COL6A1/2/3*, which can be found in the DEJ, but also around blood vessels, reticular dermis and hypodermis (Sabatelli et al., 2011; Theocharidis and Connelly, 2017), in DEJ it assembles anchor cells to the ECM (Kaur and Reinhardt, 2015); similarly, (3) *COL6A5* is located in the papillary area, and attaches several growth factors and MMPs (Freise et al., 2009); (4) *COL13A1* and *COL18A1* are also found in the DEJ, as well as in vascular and epithelial BMs (Peltonen et al., 1999; Bonnet et al., 2017; Pehrsson et al., 2019); (5) *COL4A2/4* is similarly found in the lamina densa of BMs (Barbieri et al., 2014). There are three additional collagens: *COL14A1* and *COL21A1* FACIT collagens; and transmembrane *COL23A1*, which is mainly located in epithelia (Kehlet et al., 2019b). Interestingly, some collagens, like *COL14A1* or *COL6A5*, are also found in reticular dermis (Nauroy et al., 2017; Theocharidis and Connelly, 2017). This may be because these genes are also expressed by other populations that are likely to be reticular–e.g. *COL14A1* is also expressed by A1 and A3, and *COL6A5* is also expressed by B3–or because this population, aside from being located in the papillary area and DEJ, is also necessary for proper endothelial BM functioning, regardless of the location of the blood vessels.

Other relevant ECM components expressed by A2 are *COMP*, which binds collagens I, II, IX, XII and XIV (Farina et al., 2006; Agarwal et al., 2012); and *POSTN*, which binds collagen I, fibronectin and tenascin C, and is also found at the DEJ (Yamaguchi, 2014). Specifically with collagens XII and XIV, COMP acts as a bridge between these collagens located in the anchoring plaques and collagen I, to stabilise the structure (Agarwal et al., 2012). Regarding *POSTN*, it is interesting to recall that it binds tenascin C, expressed by A2, C and B populations, in contraposition to tenascin XB, expressed by A1/A3/A4 populations. Some of the ECM components previously mentioned, as well as components secreted by other A populations, are described in Figure S20.

Besides ECM components, A2 population also expresses a large array of ECM modulatory molecules, that are known to either bind ECM components (1) without knowing their function, (2) to change the properties of the ECM, or (3) to have a secondary function not involved with ECM modification. For instance, *APCDD1* acts like *PCOLCE*, expressed by A1, which cleaves pro-collagen molecules into their active form, and also binds BMP1, another similar procollagen proteinase, to enhance its activity (Baicu et al., 2012); and *LOXL2*, a member of the lysyl oxidase family, crosslinks collagen chains to make them more resistant to insults (Sarrias et al., 2004). Interestingly, *APCDD1* is reported to be expressed by reticular dermis too (Solé-Boldo et al., 2020), possibly due to being expressed by C2 and C5 populations. On the other hand, *ADAMTS9* and *MMP11* are known to degrade, respectively CSPG such as ACAN or VCAN (Kelwick et al., 2015), collagens IV, V, IX, X, XI, vitronectin, fibronectin and laminins, as well as activate several proMMPs (Kahari and Saarialho-Kere, 1997). Some of this modulatory components are described in Figure S20.

Additionally, A2 population also expresses (1) *CD44*, a pleiotropic molecule involved in fibrosis and TGF-β signaling, immune activation and cell proliferation, which binds HA, fibronectin, proteoglycans, and collagens I, IV and XIV (Jalkanen and Jalkanen, 1992; Fujimoto et al., 2001; Kawashima et al., 2000; Bennett et al., 1995; Ishii et al., 1993). It may maintain the levels of collagen I, N-cadherin and fibronectin in homeostasis (Tsuneki and Madri, 2015) and wounding, avoiding its accumulation (Govindaraju et al., 2019).(2) *TGFBI*, which codifies a multidomain protein that may bind integrin αvβ3, fibronectin, vitronectin, collagens–including XII–, fibrinogen and VWF (Ruoslahti and Pierschbacher, 1987; Runager et al., 2013). (3) *F13A1*, the A subunit of the coagulation factor XIII, binds fibrin, fibronectin, vitronectin, collagen VI, and other ECM components, crosslinking them (Muszbek et al., 2011).

Lastly, *ANTXR1* can bind collagen VI (Nanda et al., 2004), and might be a link between ECM and cytoskeleton, necessary for ECM-sensing and migration (Gu et al., 2010). A similar function may be exerted by sarcoglycans *SGCA* and *SGCG*–commonly expressed by muscle, and also expressed by A1/A3–, which act as a link between the cytoskeleton and the ECM (Noguchi et al., 1995).

#### Nerve and vascular

The A2 population, together with other populations like C3, or D axis, expresses several markers related to nerve and/or blood vessel physiology. For instance, *KCNQ3*, which encodes a potassium channel protein (Singh, 2003), is observed at the endings of the lanceolate complex in mice (Schütze et al., 2016). Similarly, *SEMA5A* is observed to act both as an attractor and a repulsor in neural guidance (Hilario et al., 2009); and probably also in vasculogenesis (Purohit et al., 2014). *TGFBI*, besides binding ECM components, is shown to have antiangiogenic properties (Son et al., 2013); similarly to *EDN3*, an endothelial cell vasoconstrictor (Perrine, 2007). Interestingly, *EDN3* was shown to induce pigmentation in melanocyte *in vitro* (Garcia et al., 2008), and hair plucking led to melanocytic stem cell activation via *EDN3*/*EDNRB* activation (Li et al., 2017).

#### HF

Despite several genes expressed by A2 showing evidence that it may be expressed in the papillary dermis, certain exclusive markers show that the presence of A2 population may be also extended to HF. We previously mentioned the expression of *COL7A1* and *COL13A1*, related to DEJ and BMs. Additionally, these collagens are located at the BMs surrounding DP and ORS (Tsutsui et al., 2021). We also mentioned *KCNQ3*, a K^+^ channel observed at the endings of the lanceolate complex, which surrounds and innervates the HF. Additionally, *TNFRSF19*, which in humans is located at the stratum basale, also marks IFE and infundibulum in mice HFs in telogen and hair bulb in anagen. *EDN3* is also expressed by DP cells in mouse, although not in human (Hagner et al., 2020). *WNT11* is expressed in DS and ORS cells (Lim and Nusse, 2012). Lastly, *FGFR2*–which binds FGF7 and FGF10 expressed by B4–is observed to be expressed by keratinocytes, SG (Katoh, 2009) and, in mouse HF, by matrix cells near DP (Rosenquist and Martin, 1996). Some of these components are depicted in Figure S22.

#### Wnt and TGF-*β* signaling

Linking to the putative association of the A2 population with HF, another classical pathway involved with HF homeostasis and hair cycle is Wnt signaling. A2 expresses more than 10 Wnt-related markers, which are also expressed by HF populations, mainly C2, C3 and C5. Although this evidence of Wnt signaling by A2 does not directly imply that A2 fibroblasts have to belong in HF, it is highly likely that Wnt signaling is not only focused on HF cells but also by surrounding populations–including A2, B4, or D axis–by paracrine signaling.

Among the Wnt pathway markers, we observe a minority of markers related to canonical signaling, specifically *RSPO1/3/4* canonical activators (Szenker-Ravi et al., 2018) and *DAAM2* potentiator, which aggregates Fzr/Axin/Dvl complexes (Lee and Deneen, 2012; Lee et al., 2015). The rest of the markers are involved with non-canonical signaling, either by (1) inhibition of canonical signaling by *AXIN2*, a member of APC|Axin|GSK3β complex that inhibits canonical Wnt through β-catenin inhibition (hoon Jho et al., 2002); *DKK2/3*, *PTK7* and *TNFRSF19*, which sequester LRP5/6 co-receptors and promote their internalisation (Ahn et al., 2011; Lhoumeau et al., 2011); and *NKD1/2*, inhibitors of the translocation of β-catenin to the nucleus (Angonin and Raay, 2013; Zhao et al., 2015); (2) direct Wnt inhibition by *WIF1*, which sequesters several Wnt ligands (Angonin and Raay, 2013), or (3) non-canonical markers, like *WNT11* and *PTK7*.

Interestingly, although the vast majority of markers are either (1) inhibitors of the canonical pathway, (2) general Wnt inhibitors, or (3) ligand or co-receptors of either the canonical or non-canonical pathways, we do not see any receptor being expressed by A2 population. Therefore, although some markers are located in the cell membrane, the lack of proper receptors may imply that A2 is a candidate to be a paracrine Wnt canonical inhibitor.

Regarding TGF-β signaling, A2 only expresses two related markers, also expressed by C populations: *MFAP2*, which sequesters TGF-β1 in the ECM microfibrils (Weinbaum et al., 2008), and *NOG*, a BMP7 inhibitor (Blazquez-Medela et al., 2019).

#### Immune and lipid signaling

The A2 population also shows the expression of certain markers involved in side, but relevant functions, including immune, and lipid metabolism.

We have previously mentioned the expression of *FGFR2*, which binds B4-expressed *FGF7* and *FGF10*. The binding of these two factors induces the synthesis of IL-1α (Melnik and Schmitz, 2008). IL-1α is highly relevant, acting synergistically with TNFα and also acting in a positive feedback loop to activate NF-kB signaling (Laberge et al., 2015). Another gene, *CLEC2A*, is involved in NK cell-mediated cytotoxicity by binding to its receptor (Steinle et al., 2009). *ANTXR1* is a transmembrane protein that, besides from binding collagen VI (Nanda et al., 2004), may have an immunosuppressive role, modulating M2 polarisation and T cell exhaustion (Huang et al., 2020).

Another interesting protein that binds the immune with lipid signaling is *OSBP2*. As we will see in B axis signaling, oxysterols–cholesterol-derived molecules–are relevant to mediate oxidative stress and inflammatory processes (Gargiulo et al., 2016), but also in physiological aspects of cholesterol metabolism and membrane homeostasis (Liu and Huang, 2020). In that sense, *OSBP2* may bind oxysterols and be related in the regulation of their synthesis and toxicity (Gargiulo et al., 2016).

Other two genes related to lipid signaling are *PTGS1* and *AHRR*. *PTGS1* is involved in the synthesis of thromboxanes from arachidonic acid, such as TXA2 or PGI2, involved in homeostatic and inflammatory functions (van der Heide et al., 2006). *AHRR*, on the other hand, is a TF that binds a wide list of ligands, which includes eicosanoids, bilirubin, indoles, as well as external pollutants and dietary ligands; and is related to the transcription of (1) several metabolism-related genes, including many members of the CYP450 family; (2) endocrine disruptors, and (3) proinflammatory factors such as IL-1β, TNFα, IL8, MMP1 or TNFSF13B (Vogel and Haarmann-Stemmann, 2017; Larigot et al., 2022).

Lastly, A2 population also expresses two genes involved in vitamin A transport and metabolism: *TTR*, which binds RBPs to co-transport retinol into the cells, and *CYP26B1*, which degrades all-trans retinoic acid by hydroxylation (Isoherranen and Zhong, 2019). In HF, retinoic acid contributes to the refractory telogen phase, and CYP26B1 may take part in the modulation of this step (Hovland et al., 2020).

#### A1/A3

In this section, we will discuss the main functions of A1 markers. However, we remind that many of these markers are also shared by A3 and A4. Thus, these populations, although not explicitly mentioned, may also be active in some of these functions.

#### ECM production and modulation

A1 is one of the key fibroblast subpopulations due to its collagen and ECM component-producing capacity. The most important collagen genes expressed by this population are *COL1A2* and *COL3A1*, the main chains from collagens I and III, the most abundant collagens in dermis (Henriksen and Karsdal, 2019; Nielsen et al., 2019). Interestingly, although collagen III is said to be more predominant in the papillary dermis (Barbieri et al., 2014; Stunova and Vistejnova, 2018), A2 population does not express it, so it is likely, and also expected, that either A1/A3/A4 or C axis populations, fullfil this task in the papillary dermis. The other two relevant collagens in skin are *COL12A1* and *COL14A1*, which belong to the FACIT family. These collagens show interruptions in their chains that allow the interaction and binding of other collagens, GAG chains, DCN, COMP, TNC and other ECM components (Mortensen et al., 2019). Interestingly, and similarly to collagen III, *COL12A1* is more expressed in the papillary dermis (Nauroy et al., 2017), but it is not expressed by A2.

Compared to other populations, like A2 or C axis, which express a wide range of collagens, A1 only expresses the aforementioned 4 collagens. Even nucleating collagens, relevant for the proper structure of the ECM, are not expressed by A1. However, despite that, these are the most abundant, and therefore, functionally relevant collagen components of the ECM.

Besides collagens, the A1 population also expresses other relevant ECM components like the SLRPs *PODN*, *DCN* and *OGN*, which bind fibronectin and other elements (Winnemöller et al., 1991). *DCN* also regulates the expression of collagenases (Huttenlocher et al., 1996), which play an active part in ECM regulation. Other two relevant ECM components are *HSPG2* and *TNXB*. *HSPG2*–perlecan–, binds a large array of targets such as laminin, collagen IV, V, VI, XI, elastin and FBLN2; and also binds lipids with an LDLR-like domain, as well as Wnt morphogens (Hayes et al., 2022). *TNXB*, on the other hand, interacts with type I, III, V, XII and XIV collagens, as well as decorin and integrins, and is antagonistic to the location of *TNC*, mainly in the papillary dermis and HF (Lethias et al., 2006), as shown in Figure S23.

Other proteins belonging to the matrisome, and with high binding capacity are *WISP1/2* and *MFAP5*. WISP proteins contain several domains, including IGFBP, von Willebrand type C repeats, thrombospondin type 1 repeat and cysteine knot motif. *WISP1* in particular has been observed to bind DCN and BGN (Desnoyers et al., 2001). *WISP2* is known to inhibit the binding of fibrinogen to integrin receptors (Janjanam et al., 2021). Additionally, it is upregulated in hypertrophic scars (Chaudet et al., 2020) and is involved in *PPARG* activation in adipocytes. *PPARG* despite its *clasical* role in lipid metabolism, suppresses the expression of TNFα, IL-1β and IL6 as well as MMP9 in many immune cells (Jiang et al., 1998), an activity shared by certain A1 markers such as *PI16*, or *SLPI*. Lastly, *MFAP5* is a key component in the organisation of elastic fibres by interacting with fibrillin-1/2 (Penner et al., 2002).

Certain genes are involved in the maintenance, protection and maturation of the ECM fibres. One of these genes is *CA12*, a carbonic anhydrase that catalyses the production of HCO3*^−^* and H^+^, thus regulating pH and CO2 homeostasis (Supuran, 2008). This anhydrase is also present in eccrine sweat glands, regulating Cl*^−^* levels (Na et al., 2019). The other two families of ECM regulators are LOX and PCOLCE. LOX family, composed of *LOX* and *LOXL1*–although *LOXL2* is expressed by A2 and C axis–is necessary to crosslink collagen and elastin chains–*LOXL1* favours elastin (Liu et al., 2004)–a process necessary for the stabilisation of the ECM. However, excessive activity of *LOX* may be related to stiffened ECM in ageing (Langton et al., 2012). On the other hand, *PCOLCE* and *PCOLCE2* are necessary to cleave pro-collagen molecules, activating them and allowing them to polymerise into the final helical fibres (Takahara et al., 1994; Steiglitz et al., 2002).

Another aspect of ECM homeostasis is not only its synthesis but also its degradation. ECM degradation is carried out by MMPs, like *MMP2*, which can degrade collagens I, IV, V, VII, X, XI, fibronectin, elastin, laminin, vitronectin and activate proMMPs (Cabral-Pacheco et al., 2020); and *CTSK*, a cathepsin that activates MMP9 (Christensen and Shastri, 2015), decreases *COL1A1* expression (Soundararajan et al., 2021) and catabolises collagens and elastin (Kondo et al., 2022).

This effect is primarily reverted by TIMPs, such as *TIMP2* or *TIMP3*, which inhibit different MMPs and ADAMs (Cabral-Pacheco et al., 2020). Interestingly, another inhibitor of MMP2 is *PI16*, a peptidase that is activated under ECM stress or inflammation (Hazell et al., 2016). *PI16* also cleaves B2-produced RARRES2 into chemerin, which binds to CMKLR1 (E1) to exert its functions, including immunomodulation, adipogenesis and angiogenesis (Regn et al., 2016).

A similar function is exerted by *SLPI*, an inhibitor of serine proteases, to control that ECM is not over-degraded (Nugteren and Samsom, 2021). *DPP4* is a pleiotropic dipeptidase that may regulate CXCL12, MMP1, and MMP3 levels (Ospelt et al., 2010) and its binding to certain ECM components may activate or inhibit fibroblast proliferation. For instance, binding to fibronectin induces mobilisation (Cheng et al., 2003), whereas binding to glypican 3 leads to DPP4 inhibition and reduced cell proliferation (Davoodi et al., 2007). Moreover, *DPP4* is often complexed with *ADA*, also expressed by A1, which may show an antifibrotic potential (Fernández et al., 2013) and regulates the collagen production (Marucci et al., 2022).

*DPP4* has been used in several studies as a marker for different fibroblasts, even with contradictory findings. For instance, Philippeos et al. (2018) used *Cd26/Dpp4* to isolate papillary cells, and Korosec et al. (2019) also stated its presence in the papillary dermis. On the other hand, *Dpp4* is a clear marker of z1/z2 populations from adventitia cells, as stated by Joost et al. (2020); and Tabib et al. (2018) located an SFRP2^+^DPP4^+^ population throughout the dermis.

#### Immunomodulation

*SLPI* is an inhibitor of monocyte-produced MMP1 and MMP9 (Zhang et al., 1997) and inhibits the activity of IL-1β (Zakrzewicz et al., 2019). This anti-inflammatory potential from *SLPI* and *DPP4-ADA* is also shared by markers involved in the negative regulation of the complement pathway. A1, among other populations, expresses *CD55*, which inhibits the complement by binding to C4 and C3b fragments and inhibiting the creation of C2aC4b and C3bBb complex respectively (Dho et al., 2018). Additionally, *CLU* directly inhibits the formation of the MAC (Shinjyo et al., 2021); and *DCN*, aside from its SLRP component function, binds to C1q to inhibit the initiation of the classical pathway (Krumdieck et al., 1992). The complement modulation by axis A populations is depicted in Figure S24.

Related to immune signaling, with a possible role in immune regulation, are the chemokine receptors *ACKR3* and *ACKR4*. These receptors bind, respectively, CXCL12–which binds CXCR4–and CCL19/CCL21–CCL19 binds CCR7–, ligands produced not only by immune cells but also by B2/B3 populations. Both ACKRs act as scavenger receptors for these chemokines. One of the proposed and studied functions of this scavenging is to create a gradient that favours immune cell migration (Donà et al., 2013; Berahovich et al., 2013; Lipfert et al., 2013). For instance, in lymphatic vessels, *ACKR4* is expressed in afferent lymphatic collectors. There, T cells migrate by adhesion from CCL21^+^ lymphatic capillaries into CCL21*^dim^*ACKR4^+^ collectors, where they lift and go with the lymph flow (Friess et al., 2022). However, considering the additional functions of other immune-related markers, *ACKR3* and *ACKR4* may also simply just scavenge these chemokines from the ECM to avoid unnecessary immune infiltration. This regulatory ability of ACKR molecules is depicted in Figure S25.

Two last markers that may be related to immune and vascular function are *AGTR1* and *CD248*. *AGTR1* is a receptor for angiotensin II, a potent vasoconstrictor (Bergsma et al., 1992; Barnes et al., 2005). However, in cardiac fibroblasts, *AGTR1* was observed to regulate fibroblast proliferation and collagen expression (Tadevosyan et al., 2017). On the other hand, *CD248*–endosialin–is a transmembrane glycoprotein related to stromal cell proliferation and microvascularisation during tissue remodelling (Wu et al., 2021), and may also act as a receptor of CCL17–expressed by macrophages–to induce collagen expression in fibrotic environments (Pai et al., 2020). It should be noted that the two genes show a secondary function related to proliferation, in line with the ECM production function of these fibroblast populations.

#### Metabolism

Fibroblasts are not only in charge of synthesising ECM but are also active scavengers of hormones, xenobiotics and other small molecules that may be necessary for paracrine or endocrine signaling, or which are harmful to the environment.

*SLC47A2* is a H^+^/organic cation antiporter expressed in the kidney for excretion of toxic endo or exocompounds (Yonezawa and ichi Inui, 2011). In skin, it may have a role in skin drug absorption (Alriquet et al., 2015). Among the molecules that it transports there are endogenous molecules such as creatinine, guanidine or thiamine, and exogenous molecules such as tetraethylammonium–TEA, a potent channel inhibitor–, acyclovir, ganciclovir, paraquat, metformin or DAPI (Yonezawa and ichi Inui, 2011; Avsar, 2022).

*CES1* metabolises a wide range of molecules–heroin, cocain, etc.–into their active forms; and also converts monoacylglycerides to free fatty acids and glycerol, and cholesterol ester to free cholesterol (Markey, 2010). Similarly, *MGST1* belongs to the MAPEG family, involved in eicosanoid and glutathione metabolism, and catalyses the reduction of many reactive intermediates and drugs, and aids them to be further metabolised (Morgenstern et al., 2011).

Belonging the CYP450 family, the A1 population also expresses *CYP4B1*, a member that oxidises a large set of substrates, including fatty acids, arachidonic acid and cholesterol, which is activated by hypoxia, androgens, and other factors (Röder et al., 2023). Lastly, we have *HPGD*, a member of the alcohol dehydrogenases which, similar to *CYP4B1*, metabolises a wide range of prostaglandins, mainly to regulate their levels (Yan et al., 2004; Cho et al., 2006). For instance, it degrades the IL1-derived proinflammatory cytokine PGE2 (Arai et al., 2014). It also activates resolvins, enzymes that promote restoration of normal cellular function following the inflammation (Arita et al., 2006). The functions of some of these *metabolic* genes–e.g. *CYP4B1*, *MGST1*–are compatible with the immunoregulatory function of this fibroblast population.

#### Wnt and TGF-*β* signaling

Wnt signaling is partially present in A1 population, with the expression of *FZD6*, a receptor of canonical and non-canonical signaling (Corda and Sala, 2017); *LGR5*, a co-receptor of RSPOs to sequester ZNFR3 and allow Wnt binding to the receptor; and *CTHRC1*, a pleiotropic protein that binds FZRs and induces their activation (Mei et al., 2020). On the other hand, we also observe the expression of *DKK1*, a canonical Wnt inhibitor (Ahn et al., 2011), and *WIF1*, a Wnt regulator that sequesters Wnt ligands in the ECM (Gajos-Michniewicz and Czyz, 2020).

Contrary to the possible non-canonical Wnt signaling in this population, genes involved in TGF-β signaling expressed by A1 show a marked inhibitory effect, in line with the anti-fibrotic potential from *DPP4* and *ADA*. This effect could be contrary to the proliferative effect of *AGTR1* and *CD248*, although this effect could be pro-fibrogenic and thus compatible with *ADA* and *DPP4*, or could not be active in this fibroblast population.

*CTHRC1* expression does not only control Wnt signaling but also attenuates TGF-β pathway by induction of proteasomal degradation of Smad2/3 complex (Myngbay et al., 2021). *CILP* shows a similar inhibition mechanism by impeding the phosphorilation of Smad2/3, and also by binding to TGFBR (Liu et al., 2020). Similarly, *SLPI* has an additional function of suppressing TGF-β (Ashcroft et al., 2000).

This effect is mainly associated with canonical TGF-β signaling. Regarding non-canonical pathways, we see that A1 expressed *GDF10/15*, which act similarly to BMP proteins (Wang et al., 2021a). However, the A3 population expresses *SOSTDC3*, an antagonist of BMP protein (Faraahi et al., 2019), which may counteract GDF proteins.

#### A4

In this section, we will refer to the markers of A4 and their main functions. However, most of these markers are already described in A1 so that we will focus on the new markers. Regarding these markers, most of them are only more expressed in the A4 population that in the A1 population, and only a few of them are exclusive of this population. Thus, newly attributed functions in A4 may also likely be performed by the A1 population.

#### ECM production and modulation

Most ECM components expressed by A4 have already been described: *COL1A2*, *COL3A1*, *COL12A1*, *DCN*, and *MFAP5*. There are additional elements expressed that are relevant for the ECM, such as *EFEMP1*, a matrisome protein containing several EGF-like repeats and a fibulin-type C-terminal domain (Sun et al., 1998); or *PRG4*, a proteoglycan usually found in cartilage due to its high viscosity, which binds to hyaluronan and fibronectin (Rhee et al., 2005; Eguiluz et al., 2015).

Three additional genes are related to the synthesis of elastic fibres and microfibrils: *FBN1*, *ELN* and *EMILIN2*. *FBN1*–fibrillin 1–, is a glicoprotein that constitutes a structural component of microfibrils (Lee et al., 2004). These fibres can be associated with elastic fibres and elastin-independent networks (Jensen and Handford, 2016), and can also bind other matrisome proteins, including BMPs or LTBPs, playing an indirect role in TGF-β response (Jensen and Handford, 2016). *ELN*–elastin–is one of the key components of elastic fibres, which also needs the binding of GAGs, fibrillin, heparan sulfate and other ECM components (Gheduzzi et al., 2005). Elastin, being more than 1000 times more flexible than collagen, is key to the flexibility of the ECM (Kristensen and Karsdal, 2016). Lastly, *EMILIN2* is another component of the microfibrils, which binds fibrillin to other structures (Doliana et al., 2001; Schiavinato et al., 2016).

In order for fibroblasts to secrete ECM and, more generally, move alongside the ECM, they express certain markers involved in cellular motility. Some of these markers, like *SGCA/G* are also key markers of the rest of A axis populations. In this case, A1 and A4 express additional markers related to sensing and motility: *PIEZO2*, *DBN1* and *TPPP3*. *PIEZO2*, a mechanosensory channel located in Merkel cells (Wu et al., 2017), could potentially sense variations of pressure and tension in ECM to modify its structure. *DBN1* and *TPPP3* bind to F-actin in growth cones and filopodia and to microtubules, respectively (Butkevich et al., 2015; Vincze et al., 2006) Finally, regarding ECM modulators, all the genes expressed by A4 have already been described for A1.

#### Metabolism

Metabolism of complex molecules is a hallmark of A1 population. In the A4 population, in addition to *MGST1* and *HPGD*, two more genes are expressed: *PLA2G2A* and *PTGIS*. *PLA2G2A* is a phospholipase that transforms different phospholipids into arachidonic acid, which can later be transformed into eicosanoids by enzymes like *HPGD*. Another enzyme is *PTGIS*, which transforms PGG2 into PGI2–prostacyclin–, which is involved in the regulation of HF regeneration. In fact, higher levels of PGI2 and related to androgenic alopecia (Chovarda et al., 2021), and minoxidil inhibits the action of PGI2 (Messenger and Rundegren, 2004). Thus, A1 and A4 fibroblasts are active members of arachidonic acid metabolism due to the expression of *MGST1*, *CYP4B1*, *HPGD*, *PLA2G2A* and *PITGIS*.

#### Wnt signaling

Compared to A1, TGF-β signaling is not present as the expression of specific markers, whereas Wnt signaling is apparent. Besides the expression of *CTHRC1* and *DKK1* regulators, A4 population also expresses *APCDD1L*, which is predicted to be involved in the negative regulation of Wnt pathway (Otsuki, 2005); *SFRP4*, which acts similarly by sequestering Wnt ligands; and Wnt canonical ligands *WNT2* and *WNT10B*. *WNT2* may promote fibre deposition by fibroblasts (Cai et al., 2017), and *WNT10B* has been studied in the context of DP proliferation and maintenance (Ouji et al., 2012).

#### Vasculature

Although A1 population was indirectly related to vascular homeostasis with the expression of genes such as *AGTR1* of *CD248*, the A4 population may show a greater implication in this matter. PGI2, synthesised by *PTGIS*, has been shown to activate *PPARD* and induce VEGF production (Wang et al., 2013). Another relevant marker is *MGP*, a protein with affinity for Ca^2+^ cations, which, aside from decalcifying elastic fibres, is also necessary to avoid the calcification of vascular cells.

Two other markers largely studied in the vasculature and cell migration context are *SEMA3C* and *SEMA3E*, belonging to the semaphorin family. SEMA3C binds NRP1 (A1/A4), NRP2 (D and C) and Plexin D1 molecules, and shows a mixed angiogenic profile, which is pro or antiangiogenic dependent on the tumour type (Neufeld and Kessler, 2008). Similarly, SEMA3E binds Plexin D1 only, with a similar mixed profile (Neufeld and Kessler, 2008).

#### Immune response

Although certain genes with some links to immune signaling have been discussed for A1 population–complement inhibitors *CD55* and *CLU*–a large set of similar markers is expressed by the A4 population. One of the mentioned markers, *SLPI*, aside from its preotease function associated with ECM, also regulates immune signaling by inhibiting NF-kB and TLR signaling (Nugteren and Samsom, 2021; Greene et al., 2004), and also inhibits the maturation of IL-1β (Zakrzewicz et al., 2019) and IL6 (Zakrzewicz et al., 2019). *PRG4* proteoglycan expression can also target to immune-related CD44. Binding of PRG4 to CD44 internalises it and targets the inflammasome, reducing IL1β expression, and affecting TLR-mediated cascades, including NF-kB (Richendrfer and Jay, 2020). Similarly, DPP4 may truncate T chemoattractant CXCL12–produced by B2/B3–into an inactive form that only binds ACKR3, and not CXCR4 (Elmansi et al., 2022).

Four additional molecules with pleiotropic functions involved in immune signaling are *C1QTNF3*, *PDPN*, *HSD3B7* and *MGP*. (1) *C1QTFN3* inhibits IL1 and TNFα (Guo et al., 2020); but also TLR4, IL6, VCAM1, ICAM1, and E/P selectins (Schmid et al., 2021); (2) *PDPN* may favour the motility of NK cells, neutrophils and DCs (Sobanov et al., 2001; Kerrigan et al., 2009; Seymour et al., 2016), and a subset of PDPN^+^ cells form a reticular network near lymph nodes was observed to facilitate leukocyte migration and antigen presentation via CCL19, CCL21 and IL7 (Fletcher et al., 2015); (3) *HSD3B7* participates in the degradation of the proinflammatory molecule 7α,25-dihydroxycholesterol–oxysterol–synthesised by CH25H and CYP7B1 from cholesterol by B2/B3 fibroblasts; and (4) *MGP* has additionally been involved in the downregulation of TNFα, IL-1β and NF-kB; as well as inhibition of Ca^2+^-dependent inflammatory processes (Viegas et al., 2017).

Lastly, there are two markers that are related with T cells: *CD70* and *TRAC*. *CD70* is a cytokine that belongs to the TNF ligand family. It induces the proliferation of costimulated T cells, enhances the generation of cytolytic T cells, and contributes to T cell activation (Han et al., 2016). For instance, in the skin, LCs present CD70 to augment CD8^+^ T cell presence in the epidermis (Polak et al., 2012). On the other hand, *TRAC* encodes part of the TCR molecule, which binds to MHC molecules and activates the adaptive immune response in the T cell (Brownlie and Zamoyska, 2013). The presence of TRAC in fibroblasts is surprising since the presence of the TCR is supposedly restricted to T cells, and its expression in fibroblasts has not been recorded in the literature. Interestingly, *TRAC* only encodes the α subunit of the receptor, and since the β subunit is not expressed in fibroblasts, it is likely that TRAC is constitutively expressed without exerting a relevant function, or that it exerts an unknown function in the A4 population. The putative relationship of A4 fibroblasts with T cells, as well as the modulation of vasculature, are depicted in Figure S26.

### B axis

#### B1

B1, together with B2, is one of the initially discovered immune-related fibroblast populations. There is a marked difference between B1 and B2 markers and therefore, it is likely that immune response in the dermis, as with many other organs, requires different types of responses.

B1 shares some relevant markers with the z1 population, including *CCL2*, *CXCL2*, *FOSL1*, *IL6*, or *GCH1*. Interestingly, this mouse population also shares relevant markers with A4 population, such as *CD248*, *DPP4*, *EMILIN2*, *NPR1*, *PTGIS*, *SEMA3C/E*, *SFRP4* and *WNT2*. Thus, the z1 population, due to its restricted location in mice, may serve the double function of these two populations to act as an ECM-producing and immune-related fibroblast. This is not surprising considering the array of immune-related markers expressed by the A4 population.

#### Immediate Early Genes

Immediate Early Genes are genes that are transcribed within minutes after stimulation in response to both intrinsic and extrinsic signals. Many of these responses involve cell differentiation, signaling related to the immune system and stress response (Bahrami and Drabløs, 2016). Identification of IEGs tends to be complex and depends on the system of study. However, some “universal” lists of these genes have been obtained for mammalian cells (Tullai et al., 2007), which were used during this analysis.

Among the immune populations, B1 is the one expressing more IEGs (8) compared to others like B2/B3 (4), which B1 also expresses. These genes are *IER3*, *JUNB*, *SLC2A3*, *TNFAIP3*, *ZFP36*, *IL6*, *FOSL1*, and *NFKBIA*. Some of these genes, like *FOSL1* and *JUN* belong to the AP-1 TF complex, which binds JUN family proteins (Baines and Renaud, 2017). Some of these markers belong to the acute phase immune response, which involves the production of cytokines secreted to attract the innate immune response, among other roles. These markers are discussed below and are depicted in Figures S27 and S28.

#### Acute phase chemoattraction (chemokines)

There is a family of chemoattractants expressed by B1, and to a lesser extent by B3: *CXCL1*, *CXCL2* and *CXCL3*. All of these CXCLs bind to CXCR2, and the most studied one is CXCL1 (Bautista-Hernandez et al., 2017). This chemokine, and probably CXCL2 and CXCL3 as well, binds to heparan, dermatan and chondroitin sulfate GAGs to function, after being liberated by MMPs (Wang et al., 2003). Also, GAGs seem necessary for full activation of CXCR2 as well (Wang et al., 2003). CXCL1 has a marked angiogenic potential (Murphy, 2007), which may facilitate the chemoattraction of eosinophils, basophils, macrophages, immature DCs and naïve T cells; and neutrophils, towards which shows the most chemoattractant activity (Bautista-Hernandez et al., 2017; Murphy, 2007). CXCL2 and CXCL3 both act similarly to CXCL1 (Murphy, 2007), although we note that binding of CXCL2 to ACKR1 in blood vessels is necessary for neutrophils to perform diapedesis (Girbl et al., 2018). Another factor necessary for immune cell chemotaxis is *ICAM1*, expressed by endothelial cells to induce leukocyte trafficking (Bui et al., 2020). Although this effect is studied in endothelial cells, a similar process may be necessary in fibroblasts to facilitate leukocyte motility within the ECM. For instance, certain HF fibroblasts express *ICAM1* to attract perifollicular macrophages, which seem necessary for proper HF functioning since *ICAM1* KO mice suffer from hair regression (Müller-Röver et al., 2000).

*ICAM1* expression is induced by *NFKB1*, one of the effector members of the NF-kB pathway, highly active in immune processes, and also expressed by B1 (Liu et al., 2017). Key chemoattractant targets induced by *NFKB1* are *ICAM1* and *VCAM1* adhesion molecules, MMPs, *COX2/PTGS2* and *NOS2* (iNOS), these last two genes involved in angiogenesis (Liu et al., 2017). Additionally, *NFKB1* expression has been observed to correlate with M1 (pro-inflammatory) macrophage polarisation and neutrophil recruitment (Liu et al., 2017). Interestingly, B1 primarily also expresses *NFKBIA* the IkBa inhibitor. This protein sequesters NFKB1 in the cytoplasm until a pro-inflammatory signal–e.g., TNFs, TCR–targets it for degradation and liberates NFKB1 (Yu et al., 2020). Therefore, the B1 population seems to be the first to react by being prepared to activate NF-kB signaling whenever a stimulatory signal is triggered.

#### Acute phase cytokines

Another key aspect of immune response is the production of cytokines that can mediate the inflammatory response. The cytokines produced by the B1 population, or induced by genes expressed by it, are mainly acute phase cytokines, that is, cytokines that induce the chemotaxis of innate immune cells, during early immune responses.

Most of these cytokines, like IL-1β, IL6, TNFα, TGFβ, are induced by many genes expressed by B1, including *NFKB1*, *IL32*, *TNFSF14*, *CD44*, and *FOSL1* (Liu et al., 2017; Alsaleh et al., 2010; Pierer et al., 2007; He et al., 2022). Unsurprisingly, these genes were also expressed as primary chemoattractants. Interestingly, one of the target chemokine genes, *IL6*, is also expressed by B1. Its expression is induced by LPS and other external agents that bind TLRs, as well as IL1, TNFα, TGFβ and other factors. In fact, as an IEG, it is one of the most readily inducible cytokines and activates the differentiation of B cells to plasma cells, peripheral T cells–Th2, Th17, Treg–, as well as ECM remodelling–production of collagen and GAGs–and fibrosis (Paquet and Pierard, 1996; West, 2019; Duncan and Berman, 1991). It also induces the secretion of more IL6, as well as different chemokines (CCL2, CCL11) and adhesion molecules (ICAM-1, VCAM-1), which mediate its chemotactic properties (West, 2019).

#### ECM regulation

One family of target genes of NF-kB signaling are MMPs. ECM plays a key role in immune response, which is observed by the expression of other genes whose targets are also ECM regulatory components. For instance, *CD44*–previously mentioned in A2 population to maintain collagen and fibronectin levels and to bind many ECM molecules–plays an active role in immune activation too. For instance, low molecular weight hyaluronic acid mediates CD44-induced IL6, CXCL1 and CXCL2 expression in fibroblasts (Vistejnova et al., 2014). Another member involved in ECM production under immune responses is *FOSL1* which, besides regulating Th17 commitment (Shetty et al., 2022), induces the expression of *TGFB1*, *FN1*, *VIM* and *MMP1/9/14* (Sobolev et al., 2022), shown in Figure S29.

Some of these MMPs, like MMP1 and MMP3 are also expressed by B1 population. MMP1 degrades mostly collagen III, but also I, II and BM collagen IV, as well as aggrecan and entactin (Cabral-Pacheco et al., 2020). It also activates proMMP2 (Kahari and Saarialho-Kere, 1997). MMP3 can degrade collagens IV, V, IX, X, and XI, aggrecan, vitronectin, fibronectin and laminins; and also activates proMMPs 1, 8, 9, and 13 (Cabral-Pacheco et al., 2020), (Kahari and Saarialho-Kere, 1997). Similarly, *ADAMTS4* metalloprotease is also expressed and can degrade CSPGs such as aggregan or versican (Kelwick et al., 2015).

The last member of the ECM regulators is *TNFSF14*, a member of the lymphotoxin system, a network of LR pairs expressed by T cells activated during viral infection (Dostert et al., 2019). TNFSF14 induced the expression of MMP9 and IL6 in synovial fibroblasts in rheumatoid arthritis (Pierer et al., 2007), which are also expressed in the skin.

#### TGF-*β* signaling

Pro-inflammatory immune signaling goes unsurprisingly hand in hand with the pro-fibrotic TGF-β pathway. However, contrary to expected association, pro-inflammatory signaling induces an antifibrotic response, as observed by B1 markers. In fact, this signaling is not only unique to the B1 population but also to B2/B3 populations, with shared and independent markers.

One of the previously mentioned markers, *CXCL1*, is also related to TGF-β signaling. It is observed that TGF-β reduces its expression (Fang et al., 2015); although the inverse might not be true, since CXCR2 KO mice resulted in decreased TGF-β1 and collagen I expression (Zhang et al., 2020), and this reduction might be a result of indirect processes. Another previously mentioned marker is *CD44*, which is observed to act as a negative regulator of TGF-β and PDGFRB (Porsch et al., 2014).

Lastly, B1 expresses *PPP1R15A*, which belongs to a family of regulatory phosphatases expressed under stress conditions. PPP1R15A is recruited to prevent the excessive phosphorylation of the translation initiation factor eIF-2A/EIF2S1. This, in turn, reverses the shut-off of protein synthesis that is initiated by stress-inducible kinases and facilitates the recovery of cells from stress (Choy et al., 2015; Santos et al., 2016). Additionally, PP1R15A down-regulates the TGF-β signaling pathway by promoting the dephosphorylation of TGFB1 (Santos et al., 2016), and also by recruiting Smad7 inhibitor and by dephosphorylating TGFBR (Shi et al., 2004).

#### Other functions

B1 expresses additional markers with mixed functions that are not directly classified into the previous categories. One example of a marker is *PTGS2* (COX2), mentioned to be induced by *NFKB1*. In fact, *PTGS2* is induced by upstream IL1 and IL6. PTGS2 belongs to the family of arachidonic acid metabolism and is involved in the synthesis of PGE2, related to fever and pain signaling, as well as other immune signaling processes (van der Heide et al., 2006).

Another marker is *GCH1*, the rate-limiting enzyme in the synthesis of tetrahydrobiopterin (BH4) (Zhang et al., 2007), a cofactor necessary for NOSs and tyrosine hydroxylase–e.g., to make compounds such as dopamine–(Lewthwaite et al., 2015). Additionally, BH4 is a potent, diffusable antioxidant that resists oxidative stress and enables cancer cell survival (Kraft et al., 2019).

A similar oxidative stress protector is *SOD2*, a protective enzyme against ROS that transforms superoxide from the mitochondrial electron transport chain into H_2_O_2_ and O_2_ (Pias et al., 2003). In skin–and presumably other tissues–SOD2 is necessary for age-related damage protection and, in fact, SOD2 expression decreases with age (Treiber et al., 2012); and SOD2 deficiency promotes aged phenotype in mice (Weyemi et al., 2012). However, despite its protective role, SOD2 is necessary for an active innate response. Lack of SOD2 impairs IFN-I and cytokine production, as well as dysregulation of NF-kB; all of which are related to active ROS production (Wang et al., 2017). Additionally, at least in lung cancers, *SOD2* expression is associated with MMP upregulation (Yi et al., 2017).

#### B2/B3

B2 and B3 populations are what would traditionally be understood as adaptive response immune fibroblasts, that is, their main function is to attract immune cells focused on the adaptive response against specific antigens, or becoming cells that produce responses against such antigens for future infections. This response is usually preceded by the innate/acute inflammatory response although both responses may coexist and be complementary.

Comparing it with mouse populations, B2 shares the highest homology with the y4 population, based on the co-expression of some of the following markers: *APOE*, *C3*, *IL33*, *MGP*, *SLCO2B1*, and *TNFSF13B*; most of which have defined functions that will be discussed throughout this section. Similarly to the B1 population, B2 also expresses IEGs such as *JUNB*, *ZFP36*, and *TNFAIP*; as well as more specific IEGs such as *SLC2A3* and *CCL2*. For instance, *SLC2A3*–a.k.a. *GLUT3*–is a monosaccharide transporter, including glucose and galactose, to supply the cell with metabolic precursors for glycolysis (Seatter et al., 1998; Deng et al., 2015).

#### Complement

B2/B3 populations and, to some extent B4, expressed key components of the complement pathway, namely C3, C6 and C7. C3 is a cornerstone of the classical and alternative pathways; and can initiate the alternative cascade on its own. It activates into C3a anaphylatoxin and C3b, which binds to C2a and C4b to activate C5. (Rutkowski et al., 2010). C5 is also activated by CTSH, produced by B2/B3 (Bhakdi et al., 2004). Additionally, C3 can also be an immune primer: it is observed that in the synovial space, C3 exposure primes fibroblasts for future immune responses, which may lead to chronic inflammation (Afzali and Kemper, 2021).

The other two members of the complement cascade expressed by dermal fibroblasts are C6 and C7. These proteins, together with the end-of-cascade protein C5b, as well as C8 and C9 form the MAC (Rutkowski et al., 2010). Interestingly, some components, like the pore protein C9, are not expressed by these fibroblasts. In fact, C9–and other complement terminal proteins–is produced by monocytes and mature DCs as well as hepatocytes (Lubbers et al., 2017), a target suppressed by hepatitis C virus to impair MAC formation and hinder immune response in the host (Kim et al., 2013a).

Therefore, B2/B3 fibroblasts express complement cascade initiators and some terminal components as part of the innate response, but a proper response is reliant on C9-expressing immune populations.

#### Chemoattractant

One of the main functions of B2/B3 populations is the chemotaxis of adaptive and innate immune cells, shown in Figure S30. A clear marker is *CCL19*, a chemoattractant of CCR7^+^ cells, such as DCs, B cells, NK cells and several types of T cells (Robbiani et al., 2000; Reif et al., 2002; Ohl et al., 2004; Laufer et al., 2019). In APCs, it is observed that migration occurs after CCR7 phosphorylation and internalisation (Tian et al., 2013; Anderson et al., 2015). Moreover, CCL19-induced chemoattraction is also observed in certain tumors, which favours T CD8^+^ infiltration and antitumorigenic response (Cheng et al., 2018).

Another member of the CCL family expressed by B2/B3 is *CCL2*, which binds CCR2 and CCR4, expressed by bone-marrow-derived monocytes (Vanbervliet et al., 2002; Craig and Loberg, 2006). Together with CCL5 it is involved in epidermal-to-dermal migration of LCs after injury (Ouwehand et al., 2010). Additionally *CCL2* is also expressed as a regenerative signal in the context of HF. It was observed that hair plucking led to the release of CCL2 to recruit macrophages that help in adjacent hair creation by activation of regenerative signals (Chen et al., 2015; Rahmani et al., 2020).

Two additional expressed chemokines are CX3CL1 and CXCL12. CX3CL1 binds CX3CR1 to attract T cells, NK cells, monocytes and DCs (Limatola and Ransohoff, 2014; Johnson and Jackson, 2013). In fact, during wound healing, *CX3CL1* leads to the promotion of macrophage and fibroblast accumulation (Ishida et al., 2008); and in atopic dermatitis, it induces the retention of T cells in the skin (Staumont-Salle et al., 2014). Interestingly, *CX3CL1* promotes leukocyte attachment in physiological conditions in its anchored form, whereas cleavage by ADAMTS10 and ADAMTS17 in inflammatory conditions leads to its soluble form, which increases its activity (Thelen and Uguccioni, 2016). Regarding CXCL12, it binds CXCR4 expressed by T cells and Langerhans cells (Bautista-Hernandez et al., 2017; Ouwehand et al., 2008).

B2/B3 populations also produce several chemotactic interleukins, namely, IL15, IL32, IL33 and IL34; of which IL15 works as a chemokine. IL15 shows roles in innate and adaptive responses, mainly activating B, T and NK cells (Lodolce et al., 2002), and also induces the production of IL8 and CCL2–in monocytes and fibroblasts–to attract neutrophils and monocytes (Badolato et al., 1997). IL32 has been observed to switch between pro- and anti-inflammatory programs (Heinhuis et al., 2015) and, in the pro-inflammatory program, it induces expression of pro-inflammatory IL1β, IL18 and TNFα cytokines (Alsaleh et al., 2010).

A relevant chemokine that has been previously mentioned is RARRES2, that binds CMKLR1 in the context of angiogenesis. Additionally, RARRES2 shows pro- and anti-inflammatory properties depending on cleavage by proteases (Mattern et al., 2014). For instance, it was observed to enhance the chemotaxis of immature DCs and monocytes but also reduce the recruitment of neutrophils and macrophages into inflamed tissues (Cash et al., 2008; Luangsay et al., 2009).

Other relevant molecules for proper immune response are adhesion molecules. B2/B3 populations express *ICAM1*, *ICAM2* and *VCAM1*. *ICAM2* works similarly to *ICAM1*, commented on B1 population section, although it may be necessary for neutrophil crawling between endothelial cells during diapedesis (Halai et al., 2013). Additionally, and related to this previous role, CXCL2 regulates endothelial barrier function and permeability (Amsellem et al., 2014).

Regarding *VCAM1*, it is usually expressed in endothelial cells after cytokine stimulation–i.e. IL1, TNFα, IL4, IL3–, which promotes the adhesion of lymphocytes, monocytes, eosinophils and basophils (Mantovani and Dejana, 1998; Broide and Sriramarao, 2014). In other contexts, like muscle, VCAM1 is necessary for satellite cells to communicate with other satellite cells as well as immune cells and may affect myofibril growth (Choo et al., 2017). Therefore, adhesion molecules may also be used by B axis fibroblasts to gather other fibroblasts into the inflammation site.

Lastly, *HAS2* may also work as an indirect chemoattractant, since large hyaluronic chains are necessary for protection against stressors and tissue repair, but also for proper leukocyte homing (Sussmann et al., 2004), possibly by binding chemoattractants like CXCL1 in the ECM.

#### Immunomodulation

Immunomodulation refers to the processes of induction of changes in the immune cells, either to secrete specific factors, to repress or activate them, or to induce their maturation. All B axis populations show a certain degree of immunomodulation, although B2/B3 show the greatest capacity. This immunomodulatory capacity is depicted in Figure S31

Some of the most relevant immunomodulators are the members of the MHC, necessary for the activation of T cells. B2/B3 fibroblasts, and B1 to a lesser extent, express *HLA-B* and *HLA-F*, belonging to the MHC-I family; and HLA-DRB, associated with the MHC-II family. *HLA-B* and *HLA-F* present both autoantigens and exogenous antigens produced after infection to T CD8^+^ cells (Neefjes et al., 2011); and *HLA-B* is constitutively expressed in all cells, whereas *HLA-F* expression is restricted to B cells and activated lymphocytes (Ishitani et al., 2003). Compared to the rest of the members of the MHC-I family, HLA-B loads the peptide and transports it to the membrane quicker, and shows a broader range of adaptability to peptides (Neefjes and Ploegh, 1988; Peh et al., 1998), which justifies its increased expression in immune fibroblasts. Additionally, HLA-F can act as a regulator by binding to activating and inhibitory receptors in NK and T cells (Lin and Yan, 2019). On the other hand, *HLA-DRB1* is expressed mainly in APCs and captures exogenous antigens to process them and present their peptides to CD4+ T cells (Neefjes et al., 2011). Therefore, fibroblasts may also act as APCs.

Additionally, these fibroblasts also express *CD74*, a molecule that serves two main functions. On the one hand, it is an intermediary element of the MHC-II antigen protein. A specific segment of CD74, the class II–associated li chain peptide (CLIP), binds to the MHC-II antigen binding site and prevents its premature binding to antigenic peptides (Rosenzweig, 2018). On the other hand, *CD74* can also be expressed as a surface receptor, where it forms a dimer with CD44 to activate NF-kB and related pathways and induce the expression of IL6, TNFα, TLR4 and the antiapoptotic protein Bcl-2 (Williams et al., 2022).

One key molecule for immune maturation is CD40. CD40 interacts with CD40L, expressed as a surface receptor in several immune types, such as DCs or T cells, inducing their maturation (Smith, 2005). For instance, activation in macrophages leads to the expression of TNF surface receptor expression, and activation in B cells leads to their differentiation into plasma cells (Kawabe et al., 1994). A protein similar to CD40 is TNFSF13B, which actually belongs to the same family, and is also expressed by B2/B3 (So and Croft, 2013). The binding of TNFRSF13B to a TRAF subfamily receptor induces DC activation and B cell maturation (Rickert et al., 2011).

A second family of proteins involved are the immunomodulatory interleukins IL15, IL33 and IL34–which were also chemotactic–; and the soluble form of IL11 receptor, IL11RA. IL11RA binds gp130, a transmembrane protein associated with IL6 receptors, and induces IL11 signaling, even on cells with no IL11 receptor (Lamertz et al., 2018; Lokau et al., 2016). IL11RA is involved in the activation of several downstream pathways, including MAPK, JAK-STAT, NF-kB or Akt (Balakrishnan et al., 2013). IL15 acts as an activator of adaptive responses, especially T, B and NK cells (Lodolce et al., 2002). In T cells, this process is also mediated by suppression of apoptosis, which enhances its function (Malamut et al., 2010). Regarding IL33, it is stored in the nucleus and released during injury, working as an alarmin (Haraldsen et al., 2009) that is activated by tryptases and chymases released by other immune cells, like mast cells (Eissmann et al., 2020). IL33 activates T, NK, DC and mast cells, as well as basophils, eosinophils and macrophages to produce other cytokines and chemokines (Bonilla et al., 2012; Moro et al., 2009; Price et al., 2010; Pecaric-Petkovic et al., 2009). Lastly, IL34 induces the differentiation of monocytes and macrophages through binding to CSF1R (Lin et al., 2008) and SDC1 (Segaliny et al., 2015).

A similar macrophage activator is CSF1, which also binds CSF1R and regulates macrophage differentiation. However, its range of action is systemic, compared to IL34, which is focused on the central nervous system and skin (Nakamichi et al., 2013; Greter et al., 2012). Other macrophage activators are SLCO2B1 and C3. Interestingly, C3 is secreted in blood vessels by adventitial fibroblasts in vesicles containing other components and induces macrophage reprogramming (Kumar et al., 2021).

Another set of markers indirectly related to immune processes are *CH25H* and *CYP7B1*. Both proteins are related in the production and metabolisation of oxysterols, a family of lipids related to cholesterol that is highly reactive and induces cytotoxic and pro-apoptotic responses by interfering with membrane lipids (Olkkonen et al., 2012). Additionally, oxysterol itself and metabolising enzymes indirectly are highly related to immune function. *CYP7B1* expression is induced by innate cells and is related to the regulation of immunoglobulins by B cells (Dulos et al., 2005) and, together with *CH25H* the activation of macrophages and DCs (Olkkonen et al., 2012). Also, 25-hydroxycholesterol, a product of CH25H, is involved in the induction of TNF and IL6, and dermal Tγδ17 cells require it for homing in that area (Frascoli et al., 2023).

Other markers related to the immune signaling mediated by lipids are *APOE* and *APOC1*. Aside from their *traditional* role in cholesterol and lipid transport and metabolism, these apolipoproteins show a relevant immune activity. For instance, (Fuior and Gafencu, 2019) reported a significant effect of APOC1 in immune modulatory processes, and APOE is observed to suppress T cell proliferation, neutrophil activation, and regulate macrophage function (Zhang et al., 2010). Additionally, it is likely that the lipid transport function associated with the lipoproteins APOC1–present in VLDL and HDL–and APOE–present in VLDL and IDL–is necessary for the synthesis and metabolism of oxysterols by CH25H and CYP7B1.

Lastly, two additional markers show an immunomodulatory function: *IRF1* and *SOCS3*. On the one hand, *IRF1* TF is associated with the expression genes related to (1) IFN signaling (*IFNA/B*, *TNFSF10*, *ZBP1*), (2) regulation of cell cycle and proliferation (*TP53*, *CDKN1A*), (3) antibacterial response and angiogenesis (*NOS2*), (4) apoptosis (*CASP1/7/8*), (5) immune response (*IL12/15/17*, *PTGS2*), and (6) MHC expression (*B2M*, *PSME1*, *CIITA*) (Oshima et al., 2004; Dornan et al., 2004; Su et al., 2007; Park et al., 2007; Bowie et al., 2008; Huang et al., 2009; Gao et al., 2009). *SOCS3*, on the other hand, is expressed as a response to the binding of pro-inflammatory molecules–e.g. IL6, IL12, LPS, TNFα or IFNs–inhibiting the effector pathways activated by these molecules by binding to several cascade elements, like STATs or IL6R-bound JAK (Yin et al., 2015; Carow and Rottenberg, 2014). Activation of *SOCS3* may serve a double purpose: first, it may reduce the overall immune response and act as a negative feedback regulator; and may also reduce the innate, acute response to favour a secondary, adaptive response.

#### TGF-*β* signaling

Like B1, B2/B3 populations are related to the negative regulation of TGF-β signaling. There are 6 genes expressed by these populations involved in this pathway: *IL11RA*, *IL33*, *ACHE*, *PPP1R15A*, *SOCS3* and *HAS2*.

It has been observed that IL11 may induce IL33 expression in fibroblasts (Widjaja et al., 2022), and restrain them from switching to a myofibroblastic state (Gatti et al., 2021). However, IL11 by itself may show the potential to be either responsive to TGF-β signaling (Schafer et al., 2017) and also antifibrotic, at least in endothelial cells (Allanki et al., 2021). *ACHE*, acetylcholinesterase, has been shown to be indirectly related to TGF-β response inhibition (Stegemann and Böhm, 2020).

Interestingly, regarding SOCS3, TGF-β induces the suppression of this gene in fibroblasts (Dees et al., 2020). Although the contrary phenomenon–SOCS3 inhibiting TGF-β signaling–has not been described, it is highly likely that it might be true. Lastly, it is observed that *HAS2* expression in myofibroblasts is increased, which also increases CD44 ECM levels and results in fibrosis (Li et al., 2011). However, in this scenario, *HAS2* expression is already linked to the myofibroblastic state, so we cannot directly assume that HAS2 is pro-fibrotic.

#### Vasculature and ECM modulation

Vasculature and ECM modulation are necessary processes during immune responses, and although a bit diffuse, the expression of certain genes by B2/B3 populations may be involved in some of these processes.

One of the genes implicated is *ACHE*, acetylcholinesterase. This enzyme *traditionally* hydrolyses the acetylcholine neurotransmitter in neuromuscular junctions and cholinergic synapses. However, acetylcholine is also necessary for vasodilation–it is involved in NO production–, and thus ACHE may modulate blood pressure (Amezcua et al., 1988). In fact, this process is used for the migration of ACHE^+^ T cells (Fujii et al., 2017). Additionally, it is also highly likely that ACHE is involved in other functions due to the upregulation of acetylcholine-mediated response in several skin diseases like scleroderma and atopic dermatitis (Stegemann and Böhm, 2020). Similarly, *IGFBP7*, a ligand of IGF receptors, is observed to diminish angiogenesis by diminishing the activity of COX2 and PGE2 secretion, which affects VEGF production (Tamura et al., 2009).

Regarding ECM modulation, 3 genes are putatively responsible for this activity: *COL6A5*, *CTSH* and *MGP*. *COL6A5* may be indirectly associated with ECM modulation since it attaches certain MMPs, including inflammatory MMP1 and MMP9 (Freise et al., 2009). Interestingly, this gene is expressed by B3 population as well as A2, which may indicate that it possesses a certain immunogenic degree. *CTSH*, a cystein protease from the same family as *CTSK* expressed by A1/A4, is known for degrading lysosomal proteins. However, it also degrades the ECM for lymphocyte infiltration (Li et al., 2010). Lastly, MGP, which has already been mentioned to be involved in ECM and blood vessel decalcification, may be relevant for immune infiltration. Calcified ECM may be more complicated for immune cells to traverse, and therefore, its decalcification may facilitate this process.

#### B4

B4 is the last of the immune populations. It shows a clear, distinct, transcriptomic profile with markers such as *PPARG*, *FGF10*, *MYOC* or *ITM2A*. Interestingly, no immune population in mice was sufficiently matched to this population, probably indicating a unique function in humans. This population shows a diverse range of actions, including immunomodulatory actions, ECM and fibrosis response, lipid and compound metabolism, stress protection, vascular homeostasis, and, interestingly, it shows a relationship with HF.

#### Immune function

Regarding immune function, B4 shows a mixed profile, with a predominance for immunomodulation and pro-regeneratory signals, as shown in Figure S32. One of its key markers, *FGF7*, for instance, is observed to participate in reparatory processes in gut inflammation, as well as the reduction of apoptosis, and neurite reparation (Marega et al., 2021; Chen et al., 2017); and the similar family marker *FGF10*, induces the chemotaxis of immune cells in lung by secretion of IL33 to activate repairing processes (Marega et al., 2021). However, in skin, besides its alarmin and chemotactic function described in B2/B3 fibroblasts, it may have a pro-inflammatory role by inducing IL-1 secretion by keratinocytes, and vice-versa (Russo et al., 2020). A similar function is observed for IGF1, which is observed to induce M2 polarisation in macrophages, related to anti-inflammatory and pro-regenerative signals (Yunna et al., 2020).

Regarding negative immunomodulation, several genes are involved. Two of the most relevant markers of B4 are *ITM2A* and *PPARG*. *ITM2A* is involved in contributing to T helper response (Kirchner and Bevan, 1999; Tai et al., 2014), and is also involved in the induction of PD-L1, an immunosuppressive ligand (Zhou et al., 2019; Zhang et al., 2021). On the other hand, *PPARG*, a widely studied adipose tissue TF, is known to suppress the expression of *TNF*, *IL1B*, *IL6*, *MMP9* innate response genes (Jiang et al., 1998), and Th1-related *IL12*, *CD80*, *CXCL10* and *RANTES* (Szatmari et al., 2004; Nencioni et al., 2002). It is also involved in the upregulation of lipid transport and metabolism genes such as CD36 (involved in the uptake of oxLDL), FABP4 or CD1d, involved in the presentation of lipids to T cells (Tontonoz et al., 1998; Chawla et al., 2001; Szatmari et al., 2006). Thus, its immune profile might be mixed, restricting acute and innate responses, and regulating more adaptative ones.

Other two markers involved in immune response downregulation are *SLPI* and *MGP*, mentioned in A4 and A1 populations. *SLPI* inhibits the maturation of IL-1β (Zakrzewicz et al., 2019), and reduces the expression of *IL6* in mouse adipocytes (Adapala et al., 2011). Similarly, MGP contains several Glu residues that can be carboxylated by vitamin K, attracting Ca^2+^ cations (O’Shaughnessy et al., 2018; Bashir et al., 2015) that confer them the ability to downregulate *TNF*, *IL1B* and NF-kB genes, as well as inhibition of Ca^2+^-dependent inflammatory processes (Viegas et al., 2017). Lastly, the immune maturation molecule CD40 is not only expressed by B2/B3 fibroblasts, but also by B4 fibroblasts.

Besides immunomodulatory functions, B4 population is also observed to express certain chemoattractant molecules such as *CXCL12*, which attracts T cells in general (Bautista-Hernandez et al., 2017), although it has higher attractive activity for T regs (Lin et al., 2009) and Langerhans cells (Ouwehand et al., 2008).

Lastly, regarding the complement system, B4 population expresses the components also expressed by B2/B3–C3, C6, C7–but it also actively expresses *CFH*, the inhibitor of the C3bBb convertase (Jozsi, 2017).

#### HF

Interestingly, and contrary to B2/B3, B4 population expresses 10 genes involved in HF homeostasis, or expressed in this skin appendage. Some of these markers are *EFEMP1*, expressed in the bulge area (Takahashi et al., 2020); *WNT11* ligand is expressed in the dermal condensate at E14.5, and is expressed in adult HF DP and ORS (Lim and Nusse, 2012); *HSPG2* is located in the BM adjacent to DP (Tsutsui et al., 2021); *IGF1* is expressed by DP cells (Panchaprateep and Asawanonda, 2014); *FGF10* is located in DP and ORS (Zhang et al., 2018); and *MGST1* is a marker of mature sebocytes (Kobayashi et al., 2019).

Additionally, *IGF1* may be expressed in the HF regeneration context, since lower levels of IGF1 are correlated with androgenic alopecia (Panchaprateep and Asawanonda, 2014), similarly to *FGF7/10*, which also induce new HF cycles by up-regulation of β-catenin (Greco et al., 2009; Zhang et al., 2018). Similarly to FGFs, *MYOC* also intervenes in Wnt signaling as it interacts with FZDs, SFRPs and WIF, inhibiting their function, and activating Wnt pathway (Kwon et al., 2009).

The last two markers are *PLA2G2A* and *PPARG*. *PLA2G2A* overexpression is related to adnexal hyperplasia, which results in alopecia (Grass et al., 1996), and *PPARG* is involved in HF morphogenesis and maturation, and its downregulation induces a delay in HF cycles (Islam and Garza, 2018).

#### ECM

B4 population is actively involved in secreting factors that bind several ECM components. For instance, (1) HSPG2 perlecan binds heparan and chondroitin sulfate GAGs, laminins, collagens IV, V, VI, XI, elastin and FBLN2. It also binds, lipid with a LDLR-like domain, as well as Wnt morphogens (Hayes et al., 2022); (2) PODN podocan binds collagen I (Shimizu-Hirota et al., 2004); (3) FGF7/10 bind to HSPG (Marega et al., 2021), and may bind TNC (Jones et al., 2021), and (4) *MGP* binds elastic fibres.

Lastly, although it is not an ECM-component binder, *SLPI* is involved in ECM homeostasis. SLPI is an inhibitor of serine proteases–such as trypsin and cathepsin–, and MMP1/9 produced by immune cells (Nugteren and Samsom, 2021; Zhang et al., 1997). Therefore, as done by A1/A4, it may regulate and avoid excess ECM degradation. Some of these processes are depicted in Figure S29

#### Fibrosis/regeneration

B4 population, similar to the rest of the immune fibroblasts, and in line with findings from previous markers from B4 and the rest of B populations, shows a pro-regenerative and antifibrotic response. *ADA*, apart from being necessary for immune signaling, is observed to regulate collagen production (Marucci et al., 2022) and may have an antifibrotic potential (Fernández et al., 2013). Similarly, *SLPI* suppresses TGF-β (Ashcroft et al., 2000); *IGF1* shows modulation of *ACTA* expression *in vitro* (Culley et al., 2021); and *GPX3* is involved in cardiac fibroblast regulation into a pro-regenerative fate under stress conditions (Li et al., 2022a).

Interestingly, this population also expresses *GDF10*, a TGF-β ligand from the BMP7 family. Therefore it is likely that this population has either a reduced TGF-β signaling or it is linked with non-canonical signaling.

#### Protection against stress

B4 is a highly active population, and due to this activity and the involvement with other immune populations, on last relevant function is derived from the expression of “protective” markers. Similar to B1, which expressed *SOD2* to reduce H_2_O_2_, B4 expresses *GPX3*, an extracellular enzyme that reduces H_2_O_2_, hydroxyperoxides and other oxidised species (Brigelius-Flohé, 2006; Maiorino et al., 1995), and its expression is upregulated by PPARG (Reddy et al., 2018; Chung et al., 2009). Similarly, B4 fibroblasts express *MGST1*, a metabolic enzyme involved in eicosanoid and glutathione metabolism. This factor, glutathione, is a highly active component against oxidative stress too (Pompella et al., 2003). Additionally, B4 expresses two markers with an associated chaperon function: *MYOC* and *ITM2A*. *MYOC* is a modulator of the actin cytoskeleton, but it also acts as a chaperone against ER stress (Anderssohn et al., 2011); and *ITM2A* contains a BRICHOS domain that is known to act probably as a chaperone in different scenarios (Hedlund et al., 2009). Lastly, *APOD* is observed to have a protective action against oxidative stress preventing lipid oxidation in different organisms (Ganfornina et al., 2008).

### D axis

#### D1

D1 population is, together with D2 and E1, a small population that was previously unidentified but with very particular markers. Human-mouse comparisons revealed that D1 was transcriptomically similar to either y5 or v1 populations. However, at a closer examination, more genes from D1-y5 are selected markers based on their function. Some of these markers are *ABCA8*, *APOD*, *COL8A1*, *SOX9*, *TGFBI*, and *VIT* ; of which *SOX9* is exclusively expressed by D1. There are other markers expressed exclusively by D1, such as the similar SOX gene, *SOX8*, *BAMBI*, *CDH19*, *FMO2*, or *ATP1A2*. Some of their functions will be mentioned later.

The most relevant functions of the D1 population, and D2, to a lesser extent, are related to the expression of nerve and blood vessel markers that are either expressed by or observed in these structures or whose function is associated with the activities performed by these cells.

Focusing on the nerve aspect of the D axis populations, specially of D1 but also of D2, is the increased number of markers linked to that structure may be a sign of not only association to nerve, but even to be part of nerve structures, like endoneural and epineural fibroblasts.

In classical literature on these fibroblasts, such as Richard et al. (2012, 2014), common fibroblast and/or perivascular markers are used, including *CD34*, *NG2*, *PDGFRB* or *NES*; which are not sufficiently specific to discriminate any fibroblast population from this analysis. A recent paper published by (Chen et al., 2021) uses single-cell to determine the heterogeneity of peripheral nerve cells and differentiates epineural fibroblasts, expressing *Sfrp2*, *Dpt*, *Pcolce2*, *Adamts5*, *Sfrp4*, *Prrx1*, *Comp*, *Ly6c1*, from endoneural fibroblasts, which express *Sox9*, *Osr2*, *Wif1*, *Cdkn2a/b*, *Abca9*, *Plxdc1*, *Apod*. A more in depth analysis on DEGs of these populations reveals that endoneural fibroblast express a large set of markers that are also markers of D1 population, as well as D2: *Apod*, *Col15a1*, *Crispld2*, *Abca9*, *Angptl7*, *Matn2*, *Sox9* and *Gfra1*, among others. Interestingly, looking at the markers of perineural fibroblasts, some of them are A or C ECM or ECM modulatory markers–*Pi16*, *Bgn*, *Aspn*, *Gpx3*, or *Col14a1*–and others are more expressed in A4 population–*Clec3b*, *Pcolce2*, *Ccdc80*, *Sfrp4*, *Fbln1*, *Cd248*, *Gpx3*, *Sema3c*–.

Therefore, D1 fibroblasts–and A4 possibly–are likely to be related to the endoneural fibroblasts, however, further research should be performed to ensure that the location of these fibroblasts match the one putatively reported in the literature.

#### Nerve function

The D1 population expresses several markers that are also markers of other cell populations. For instance, (1) *ABCA8* is highly expressed by oligodendrocytes and may play a role in sphingomyelin production (Kim et al., 2013b), (2) *SOX8* TF is involved in the development of astrocyte, oligodendrocyte and Schwann cell precursors (Stolt et al., 2004; Takouda et al., 2021; Hutton and Pevny, 2009), and also plays a role in sphingomyelin production (Turnescu et al., 2017), (3) *CDH19* cadherin is expressed by Schwann cell precursors (Kim et al., 2020b), (4) *S100B* is also a marker of Schwann cells, although it is expressed in melanocytes, chondrocytes, adipocytes and immune cells (Donato et al., 2009); (5) *PIEZO2* is a mechanosensory Ca^2+^ channel involved in propriocepcion (Woo et al., 2015), located in Merkel cells and afferent terminals (Wu et al., 2017); and (6) *COL28A1*, although not being related with a clear neural function, it is almost-exclusively detected by terminally-differentiated Schwann cells and Merkel cells (Grimal et al., 2010).

Some of these and other D1 markers have additional relevant functions in neural environments. For instance, *ATP1A2* is a Na^+^/K^+^ channel that osmoregulates the cation gradients for proper neuronal excitability in nerves and muscles (Friedrich et al., 2016); *OGN* and *VIT*, usually involved in fibroblast proliferation, participate in neural development and neurite growth (Deckx et al., 2016; Whittaker and Hynes, 2002); and *SCN7A*, another Na^+^ channel that acts jointly with *ATP1A2* in glial cells (Dolivo et al., 2021), acts as a Na^+^ concentration regulator in homeostasis and TEWL-related injuries (Watanabe et al., 2000). In this case, coactivation of *ATP1A2* to compensate Na^+^ influx leads to the hydrolysis of ATP and the production of lactate. This, together with an increase in endothelin ligands induced by *SCN7A*, leads to compensatory neural responses like an increase in water intake in the ECM or an increase in blood pressure (Dolivo et al., 2021).

#### Vasculature

Jointly with neural functioning, and as we have observed with *SCN7A*, D1 population expresses more than 10 markers with a direct or indirect function related to vasculature.

Three interesting markers are *COL15A2* and *COL8A1/2*, which are shown to be expressed at the BM of microvessels or at proliferating vessels (Manon-Jensen et al., 2019; Sutmuller et al., 1997). *EFNA1* and *NRP2*, also expressed by D2, act as ligands of receptors of several pathways involved in neural and vascular neogenesis (Hao and Li, 2020; Islam et al., 2022). In fact, NRP2 is involved in vascular permeability and lymphangiogenesis as well (Harman et al., 2020; Favier et al., 2006; Yuan et al., 2002). In a similar fashion, *BAMBI* and *TGFBI*, related to TGF-β signaling, are also indirectly related to these responses (Guillot et al., 2012), although *TGFBI* specifically shows antiangiogenic potential (Son et al., 2013).

Other members that act by inducing the secretion of pro-angiogenic factors are *IGFBP7*, which secrets PGE2, which in turn induces angiogenesis (Tamura et al., 2009); *NFKB1* and *S100B*, which are involved in the secretion of iNOS (Liu et al., 2017; Donato et al., 2009); or *EPHX1* and *CYP4B1* which metabolise multiple substrates involved in angiogenic responses (Gautheron and Jéru, 2020; Tang et al., 2010). Lastly, both D1, D2 and E1 populations express the kallikrein *KLK1*, responsible for the production of bradykinin (Bellis et al., 2020), a vasoactive substance involved in blood vessel dilation by the production of NO and prostacyclin, as well as derived inflammatory processes (Pinheiro et al., 2022).

#### Immune

B axis is the principal immune axis. However, there are certain markers that are related to this family of responses specifically expressed by D1. Some markers, like *NFKB1* and *SOD2*, already described in B1, or *CCL2*, *CFH*, *RARRES* or *SOCS3*, also expressed by B axis population, showcase the putative function of D1 in immune roles; although due to the pleiotropism of some of these genes, their functions in this population could be different. Specific to D1 are three additional markers: *C2orf40/ECRG4*, *CYP4B1* and *ATP1A2*. *ATP1A2*, aside from its function in nerve, is also observed to modulate LPS responses by NF-kB signaling (Leite et al., 2020); a similar activation mechanism of *C2orf40*, which also activates IFN-I mediated signaling (Moriguchi et al., 2016) and neutrophil recruitment (Dorschner et al., 2020). Lastly, *CYP4B1* is upregulated by pro-inflammatory cytokines (Smerdova et al., 2014), so downstream effects of this activation may also be immune-related.

#### TGF-*β*

Regarding TGF-β signaling, D1 also expresses *BMP7* and *INHBA*, a combination postulated for C5 population to inhibit canonical signaling and activate the BMP-mediated alternative pathway. In addition, D1 expresses *GDF10*, which acts as *BMP7*.

In a similar fashion to HF populations, TGFB-β signaling in D1 is complex, with additional expression of markers such as *PPP1R15A*, which recruits Smad7 to inhibit TGF-β (Shi et al., 2004) and *BAMBI*, which is also inhibitory, by binding TGFBRI/TGFBRII complex (Huang and Chen, 2012). On the other hand, *SCN7A* expression linked to increased acidity in ECM–from sensing TEWL after injury–is related to latent TGF-β activation (La et al., 2016). In a similar fashion, *SOX9* TF may lead to fibrosis by induction of *ACTA2*, collagens and *LOXL2* (Gajjala et al., 2021; Scharf et al., 2019); and is related with hfSC niche reestablishment by TGF-β signaling (Berndt, 2014). This stemness state, although regulated by NOTCH in specific environments–e.g. gastric cancer–, and not TGF-β signaling, may be induced by *SPON2* (Badarinath et al., 2022).

#### Wnt

Wnt signaling, similar to C axis populations, is active in D axis populations. In D1, it is probable that inhibitory signaling is more dominant, with the expression of *DKK3*–sequesters LRP5/6 (Ahn et al., 2011)–, *SFRP4*–binds Wnt ligands and inhibits their action (Liang et al., 2019)–, *SHISA3*–impedes the translocation of Wnt receptors to the membrane (Hedge and Mason, 2008; Furushima et al., 2007)– or *NDRG2*–activates GSK-3β, which signals β-catenin degradation (Lee et al., 2022)–. Thus, the expression of *FZD2* non-canonical receptor (Gujral et al., 2014) may not be sufficient for this signaling.

#### Collagens and ECM regulation

Interestingly, D1–and D2 to some extent–shows increased expression of 5 collagen genes: *COL8A1/2*, found in proliferating vessels (Sutmuller et al., 1997); microvessel-associated *COL15A1*; and nucleating *COL9A3* and *COL28A1*, found around Merkel and Schwann cells (Grimal et al., 2010). Another related ECM component related to collagen is *DCN*, not only expressed by A axis, but also by D1.

Additionally, D1 population expresses *CYP4B1*, which is related to *POSTN* expression (Zhao et al., 2013); or *VIT* –vitrin–, acting similar to COCH in cell adhesion and neural development (Whittaker and Hynes, 2002). It also expresses several genes whose expression products bind or regulate multiple ECM components, such as *TGFBI*, which binds integrin αvβ3, fibronectin, vitronectin, collagens, fibrinogen and VWF (Ruoslahti and Pierschbacher, 1987); *EGFR*, which binds AREG, TGF-α, or EREG (Normanno et al., 2006; Harris, 2003), and other ligands like TNC, DCN, or laminin-332 (Santra et al., 2002; Iyer et al., 2007; Schenk et al., 2003); and *ANGPTL7*, which is associated with a diminished expression of *FN1*, *COL1A1*, *COL4A1*, *COL5A1*, *MYOC* and *VCAN*, and an increased expression of *MMP1* (Comes et al., 2010), in line with a pro-inflammatory profile.

Although it is complex to establish a clear function of these genes in the context of the D1 population, it is possible that this type of fibroblast transforms “classical” ECM microenvironments into more specialised environments necessary for the correct development of HF, nerve or blood vessels.

#### Vitamin A and lipid metabolism

Regarding lipid and retinol metabolism, three related genes are expressed by D1: *ALDH1A3*, which transforms retinaldehyde into retinoic acid (Kedishvili, 2013); and two lipoprotein-related markers: *APOD* and *LDLR*. *APOD*, despite belonging to the apolipoprotein family, is also highly related to RBPs in function (Munussami et al., 2018); thus, it may be the agent transporting retinol into D1 fibroblasts. Lastly, *LDLR*, which binds VLDL and LDL in plasma (Go and Mani, 2012), may be related to lipid metabolism; or, considering the adipose tissue surrounding anagen HFs, may be an active component in the communication with hypodermis.

#### FMOs

Lastly, the family of flavin mono-oxygenases (FMOs), is highly expressed by B4, D1 and D2 populations; and in the case of D1, is the only population expressing all of the members expressed in the samples from the analysis: *FMO1*, *FMO2* and *FMO3*. In general terms, each FMO metabolites, generally using FAD or NADPH as cofactors, a wide range of molecules including triethylamine and other secondary and tertiary amines, thiols and other xenobiotics (Hisamuddin and Yang, 2007). Although FMOs activity has been linked with fibroblast activation (Yu et al., 2022) and cardiovascular disease (Schugar and Brown, 2015; Shih et al., 2019; Warrier et al., 2015), specific functions of these enzymes are yet unknown.

#### D2

D2 is the second population from the D axis. Although both populations differ in certain markers and possibly in some of their functions, their transcriptomic similarity is extremely high, and the most relevant genes are expressed in both populations. Regarding mouse-human comparisons, D2 shows a clear similarity to v1 population, with exclusive markers, such as *AQP3*, *CAV1/2*, *ITGA6*, *ITGB4*, *KRT19*, which are all exclusively expressed by D2. Additionally, D2 also expresses other exclusive markers, including *DACT1*, *ADAMTSL5*, *AQP3*, *GFRA2* and *NGFR*, many of which will be discussed throughout this section.

#### Nerve and vasculature

Similar to D1, D2 population seems to be associated with nerve and blood vessel microenvironments. Regarding nerve markers, population D2 shows the expression of already mentioned D1 markers such as *ABCA8*, *S100B*, and *VIT* ; but it also expresses specific markers. For instance, (1) *GFRA2*, a member of glial cell-derived neurotrophic factor (GDFN), is necessary for neurotransmitter release of several neuron types (Airaksinen and Saarma, 2002); (2) *KRT19* and (3) *NGFR* are expressed in Merkel cells–as well as fibroblasts, mast cells and rete-ridge keratinocytes–(Botchkarev et al., 2006; Michel et al., 1996), and *NGFR* is necessary for Schwann cell cone formation (Cragnolini and Friedman, 2008); and (4) *SLC22A3*, a channel involved in neurotransmitter transportation (Amphoux et al., 2006), which may be necessary for the regulation of osmolality in the neuron vicinity (Vialou et al., 2004).

Regarding vascular markers, D2 population also expresses D1 markers *NRP2* and *EFNA1*; as well as *CAVIN2*, which will be discused later, and *SEMA3C*, which binds NRP2 and shows a mixed angiogenic profile (Neufeld and Kessler, 2008; Jiao et al., 2021; Valiulytė et al., 2019; Karpus et al., 2019).

#### Immune

D2 population expresses many immunogenic markers, but many of these have already been mentioned in previous populations: *CLU* (complement inhibitor, A1/A4), *CCL2* (B3), *SOCS3* (B3), *ITM2A* (B4) and *C2orf40* (D1). The only relevant new marker is *CCL13*, a pleiotropic chemokine that (1) binds CCR1/2/3/5 and induces chemotaxis of different immune cells, including Th and NK cells, mast cells, basophils, eosinophils and monocyte/macrophages (Mendez-Enriquez and Garcia-Zepeda, 2013); (2) upregulates *TLR2/3/4/5* expression and induces DC maturation (Mendez-Enriquez and Garcia-Zepeda, 2013); and (3) its expression is related to CD40 and MHC-II expression, involved in adaptive responses (Chiu et al., 2004).

Although D axis populations show some expression of regulatory immune modulators such as *SOCS3*, *CLU* and *ITM2A*, the expression of several chemokines and other markers indicate that their immune activity is not negligible.

#### HF and ECM redulation

D2 population also shows a wide range of markers associated with ECM and HF in homeostasis and other conditions.

Some markers that were mentioned to be expressed surrounding nerves, blood vessels and other structures have been shown to mark HF areas such as *KRT19*, which marks ORS in the bulge area (Michel et al., 1996); *SLC22A3*, located in HF, and SG–and may be associated with sebum production–(Takechi et al., 2021); *ADAMTSL5*, also located in ORS (Higgins et al., 2011); *GFRA2*, expressed in ORS, IRS, DP and CTS in anagen, although its CTS expression is maintained throughout the HF cycle (Adly et al., 2008); and *NGFR*, expressed in DP.

In fact, some markers, like *NGFR*, are also associated with the HF cycle or balding processes. For instance, *NGFR* is expressed in the anagen-to-catagen transition (Botchkarev et al., 2000; Enshell-Seijffers et al., 2010); *FGF7*, which signals the induction of new hair cycles (Greco et al., 2009; Geyfman et al., 2014); *CCL13*, whose immune function, if exacerbated, is associated with T cell accumulation in alopecia areata (Wang et al., 2021b); and *PTGDS*, which is highly expressed in androgenic alopecia (Garza et al., 2012).

Regarding ECM regulation, the D2 population expresses three different ADAMTSL members: *ADAMTSL3*, *ADAMTSL4*, *ADAMTSL5*. Their main function is the binding of fibrillin-1 (Sengle et al., 2012), and regulating its biogenesis (Gabriel et al., 2012), although each member has specific functions as well. Additionally, D2 also expresses *TGFBI* and *VIT*, which have been mentioned in D1.

#### Wnt signaling

Similar to D1, D2 population expresses several genes involved in these two pathways. Regarding Wnt signaling, apart from the already mentioned *NDRG2* and *SFRP4*, it expresses *DKK3*, *CAV1* and *DACT1*, all three represorst of Wnt signaling–canonical or overall–(Ahn et al., 2011; Galbiati et al., 2000; Gao et al., 2022; Esposito et al., 2021); and *DAAM1*, which activates PCP non-canonical pathway (Lai et al., 2009).

#### Lipid metabolism

While the D1 population expressed genes related to cholesterol and general compound metabolism; markers expressed by D2 are slightly more related to eicosanoid metabolism. For instance, *EPHX1* metabolises the oxidation of the endocannabinoid 2-AG into arachidonic acid (Nithipatikom et al., 2014); *CYP4B1* CYP450 monooxygenase catalyses arachidonic acid oxidation; and *PTGDS* catalyses the conversion og PGH2 to PGD2 (Zhou et al., 2010).

#### Caveolins

Caveolins are a family of scaffolding proteins associated with caveolae formation. Caveolae are membrane invaginations of lipid rafts that endocytose and are generally involved in signal transduction (Anderson, 1998). D2 population specifically expresses three caveolae-related markers: *CAV1* and *CAV2* caveolins, and *CAVIN2*–Caveolae Associated Protein 2–. Generally, although *CAV1* is able to produce caveolae on its own (Scherer et al., 1997), caveolae are usually created by the interaction of these three proteins (Mora et al., 1999). In some scenarios *CAV1* and *CAV2* may have antagonistic functions (de Almeida, 2017).

The functions of these genes are diverse and context-specific. For instance, *CAV1* is related to TGFBR1 sequestering from membrane rafts into caveolae to reduce TGF-β signaling (Hwangbo et al., 2015); may also sequester β-catetin to regulate Wnt signaling (Galbiati et al., 2000; Gao et al., 2022); may be related to immune responses by inducing T-cell proliferation (Ohnuma et al., 2007), favours neutrophil extravasation (Marmon et al., 2009) or ICAM1-mediated leucocyte adhesion (Bouzin et al., 2007). *CAV2* may be related to keratinocyte proliferation upon KGF action (Gassmann and Werner, 2000), may also act like *CAV1* in TGF-β signaling (Xie et al., 2011), or may be involved in lipid metabolism and receptor trafficking (Sowa, 2011). Lastly, *CAVIN2* may control angiogenesis by regulating eNOS activity (Boopathy et al., 2017) and may also inhibit TNF-mediated NF-kB signaling (Annabi et al., 2017).

The immune profile associated with the expression of caveolin-related genes is in favour of a mature, adaptive response, in line with the immune markers presented in this section.

#### Integrins

Lastly, an interesting heterodimer expressed by D1 principally is the integrin α6β4, formed by the expression of *ITGA6* and *ITGB4*. Similar to caveolae, the functions of this integrin dimer are diverse, but may be extremely relevant for this fibroblast population.

*ITGA6* by itself is found to serve several functions. In mouse HF, ITGA6 is located at the BM surrounding DP and ORS (Tsutsui et al., 2021); and it was observed that HF ITGA6^+^ cells were able to generate hair shaft, ORS and IRS *in vitro* and *in vivo* (Yang et al., 2014, 2020). This activity is not specific to HF, as ITGA6^+^ populations tend to show stem potential across tissues (Yu et al., 2012). Additionally, it participates in Langerhans cell migration from the epidermis into the dermis and lymph nodes (Varlet et al., 1991) and may act as an ECM mechanosensor to induce myofibroblast activation under specific circumstances (Chen et al., 2016). This effect is mediated in keratinocytes by *EGFR* binding to integrin α6β4. Under ECM stiffening, EGF bioavailability increases, activating EGFR and inducing migration and cell activation (Kleiser and Nyström, 2020).

Regarding the heterodimer, it binds several ECM and BM members, including laminins 332 and 511, and colocalises with BM collagen IV, as well as other integrins, in DEJ and HF and SG BM (d’Ovidio et al., 2007; Blok et al., 2015). Additionally, integrin α6β4 is tightly related to cell migration, both by playing a key role in the formation and stabilisation of hemidesmosomes–together with CD151 and COL17–, as well as its disassembly and translocation to lamellipodia (Wilhelmsen et al., 2006; Walko et al., 2015). Another relevant function is the maturation of the microvasculature by playing a suppressive role in angiogenesis (Hiran et al., 2003).

#### E1

Population E1 function, similar to D1 and D2, cannot be assigned based on belonging to any of the “big” axes. Additionally, there is no clear relationship between E1 and any mouse population. Looking at the combined PAGA graphs from Figures 1b and 2, we see that this population shares transcriptomic similarity with B4, D1 and C3 populations. Firstly, all three populations are directly or indirectly linked to HF, so it is also likely that this population will be too. Analysis of markers for this population reveals that, similarly to B4 or D axis, this population has a wide diversity of functions, including TGF-β and Wnt pathways, vasculature and neural regulation, proliferation, ECM, and immune signaling. Even more, due to the reduced, but diverse, number of markers, there is no function with a clear predominance, which reinforces the versatility of this population.

#### Proliferation and cell migration

One of the gene families relevant to E1 population is the insulin growth factor (IGF) signaling pathway, with the expression of *IGFBP2* and *IGF1*, which are associated with cell proliferation. *IGFBP2* may act by binding IGF1 or IGF2 (Shin et al., 2017). By binding to IGFs, it modulates their availability in the ECM. After proteolytic cleavage, IGFs may bind their receptors and their functions are related to angiogenesis and cell proliferation in general (Boughanem et al., 2021). This is related to its implication in energy metabolism; since its expression is increased after fasting, insulin suppresses it, and it is a target of PPARA (Shin et al., 2017). Similarly, IGF1 binds integrins αvβ3 and α6β4 to form a ternary complex with IGFR1 (Fujita et al., 2012; Saegusa et al., 2009) or with insulin receptor. In both cases, it strongly activates the Akt pathway, which stimulates growth and proliferation and inhibits apoptosis (Peruzzi et al., 1999; Juin et al., 1999). Interestingly, keratinocytes express IGF1R, while fibroblasts express IGF1; thus IGF1 in keratinocytes may work by paracrine modulation (Rudman et al., 1997).

Another proliferative marker is *SPON2*. It activates STAT3 signaling in cancer stem cells to reinforce their stemness state (Badarinath et al., 2022), and also upregulated NOTCH signaling in gastric cancer (Kang et al., 2020). This activation may be targeted to other cells, like macrophages, where SPON2 activation leads to its chemotaxis (Li et al., 2020).

Lastly, the E1 population expresses two markers related to migration: *PTN* and *GMFG*. *PTN* binds to several receptors, including PTPRZ1, ALK, SDC3, ITGAvB3, and NRP1 to induce cell migration (Papadimitriou et al., 2022). GMFG is a protein commonly found in T and B cells, neurons and cancer cells; and is located in the filopodia, pseudopodia, or other prolongations used in the movement of the cell (Lippert and Wilkins, 2012; Liu et al., 2022; Deretic et al., 2021).

#### HF

E1 population shows the expression of 4 markers linked to HF structure and homeostasis. Two of these markers, *IGF1* and *IGFBP2*, have already been mentioned before and are detected in DP and DS respectively (Panchaprateep and Asawanonda, 2014; Hagner et al., 2020). IGF1 is presumed to be linked to HF renovation. In fact, lower levels of IGF1 and IGFBP2 are observed in androgenic alopecia patients (Panchaprateep and Asawanonda, 2014). Similarly, *PGF* –placental growth factor–is also involved in maintaining HF growth (Hu et al., 2022; Yoon et al., 2014). Lastly, *LFNG*, a fucosyltransferase present in the Golgi apparatus that adds an N-acetylglucosamine to the fucose of the NOTCH1 receptor, reducing its activity (del Barco Barrantes et al., 1999; Shimizu et al., 2001). NOTCH signaling is necessary for HF cells to maintain cell differentiation (Nowell and Radtke, 2013). Therefore, NOTCH inhibition may be related to HF cycle, in line with some of these other markers.

#### Immune

Several previously described markers are also expressed by E1 fibroblasts, namely *MGP* and *ITM2A* from B4, and *ACKR3* from A1/A4. All these markers show a direct suppressive role in the immune system or in the clearance of immune products from the ECM. This population expresses two additional markers, *CMKLR1* and *TNFRSF21*. *CMKLR1* is the receptor for chemerin/RARRES2, predominantly expressed by B2 and B4 populations, and shown to have pro- and antiinflammatory functions; and TNFRSF21 is a proapoptotic TNF receptor (Dostert et al., 2019) that may decrease T cell and B cell proliferation (Liu et al., 2001; Schmidt et al., 2002), although it may also be necessary for their migration to the central nervous system (Schmidt et al., 2002). Therefore E1 population may show an immunomodulatory function, as transcriptomically similar populations like B4.

#### ECM

E1 population expresses 4 ECM-related markers: the *MMP6* metalloproteinase, which can degrade collagen III, fibronectin and laminin (Cabral-Pacheco et al., 2020), and *TIMP3* to counteract MMPs functions. Additionally, it expresses two collagens, *COL15A1* and *COL26A1*: *COL15A1* has already been mentioned as a non-fibrillar collagen that is located at the BM of microvessels (Manon-Jensen et al., 2019), nerves, adipocytes (Saarela et al., 1998), and HFs (Hagg et al., 1997); and *COL26A1*, specific of E1 population, is not sufficiently studied to have a clearly assigned function.

#### Neurovascular

E1 population, similar to D1 and D2, expresses genes related to neural and vascular development. Regarding the neural-related markers *SCN7A* and *SLC22A3*, already commented on D-axis populations, are also expressed by the E1 population. This population specifically expresses *NTRK3*, a member of the neurotrophin receptor family, involved in different proliferation and differentiation pathways (Ruiz-Cordero and Ng, 2020). In adults, they are usually expressed in the peripheral and central nervous system and are thought to maintain the regular neuron balance (Hechtman, 2022). In skin, it is expressed in the placode morphogenesis, specifically in Merkel cell precursors (Jenkins et al., 2019).

Regarding vascular markers, most of them are specific to E1 population, and act via VEGF signaling. For instance, *PTN* acts as a competitor of VEGFA to bind VEGFR2, showing pro-angiogenic potential (Koutsioumpa et al., 2015); *PGF* binds VEGFR1 and may act synergistically with VEGF (Odorisio et al., 2006; Chang and Johnson, 2012). Similarly, *PDGFD* is shown to increase the expression of VEGFA, as well as FGF1, INHBA and IL11 (Kim et al., 2015), thus it has a marked pleiotropic potential. Lastly, one of the many functions of IGFBP2 is also related to proliferation and angiogenesis (Boughanem et al., 2021).

#### Wnt and TGF-*β* signaling

It comes as no surprise that, as with other functions shared with D1, D2, B4 and other populations, E1 expresses markers related with Wnt and TGF-β signaling.

Regarding TGF-β signaling, there is a mixture of activatory–*SLC22A3*, *EGR2*–and inhibitory/modulatory–*LTBP2*, *IGFBP2*–signaling. Moreover, both activatory and inhibitory cases are, in some cases circumstantial. For instance, *EGR2* is observed to be induced by TGF-β in SSc fibroblasts (Fang et al., 2011), and *SLC22A3* could be indirectly related to *TGFB1* increased expression (Vollmar et al., 2019). Thus we cannot ensure that activatory signaling is clear in this population. Regarding *IGFBP2*, it was observed to decrease αSMA expression via TGFB1 inhibition (Park et al., 2015), but due to its pleiotropism, this function may not be active in dermal fibroblasts.

## Supplementary Figures

**Figure S1.**
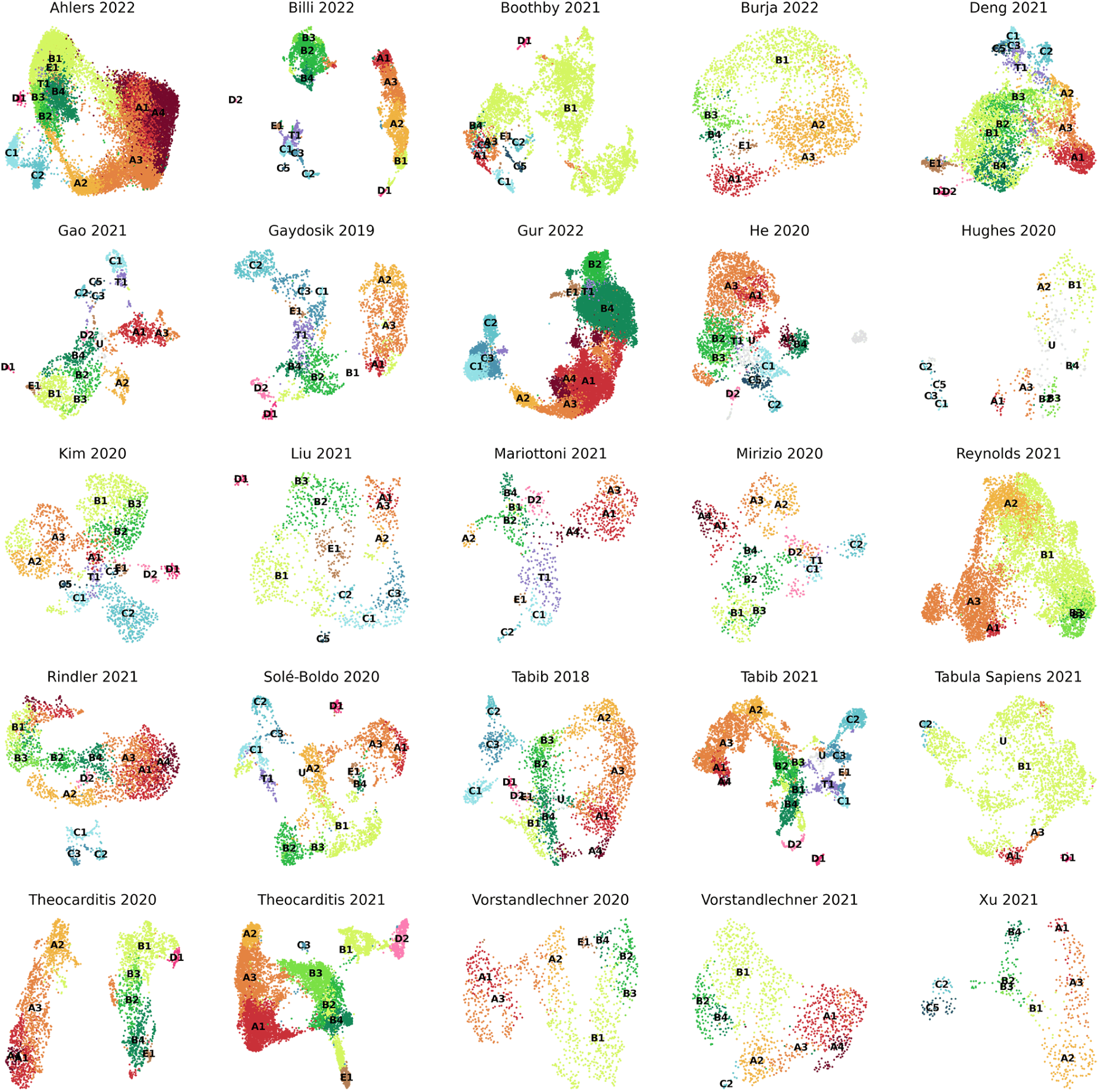
UMAP plots of all human fibroblast datasets from secondary analysis. Colours of populations are shared across datasets.

**Figure S2.**
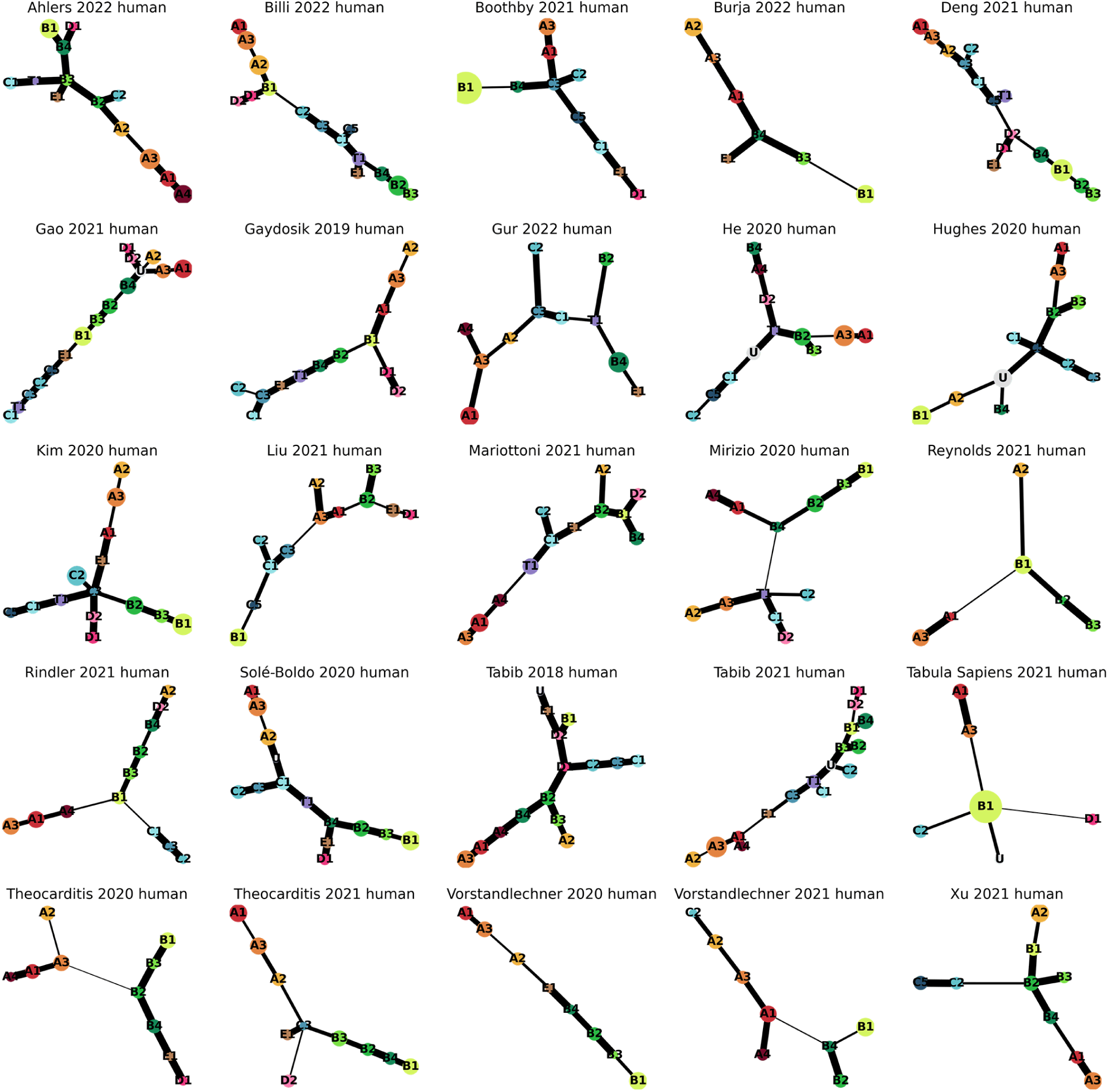
PAGA tree graph of human dermal populations. Colours of populations are shared across datasets. Thicker lines indicate greater connectivity between nodes.

**Figure S3.**
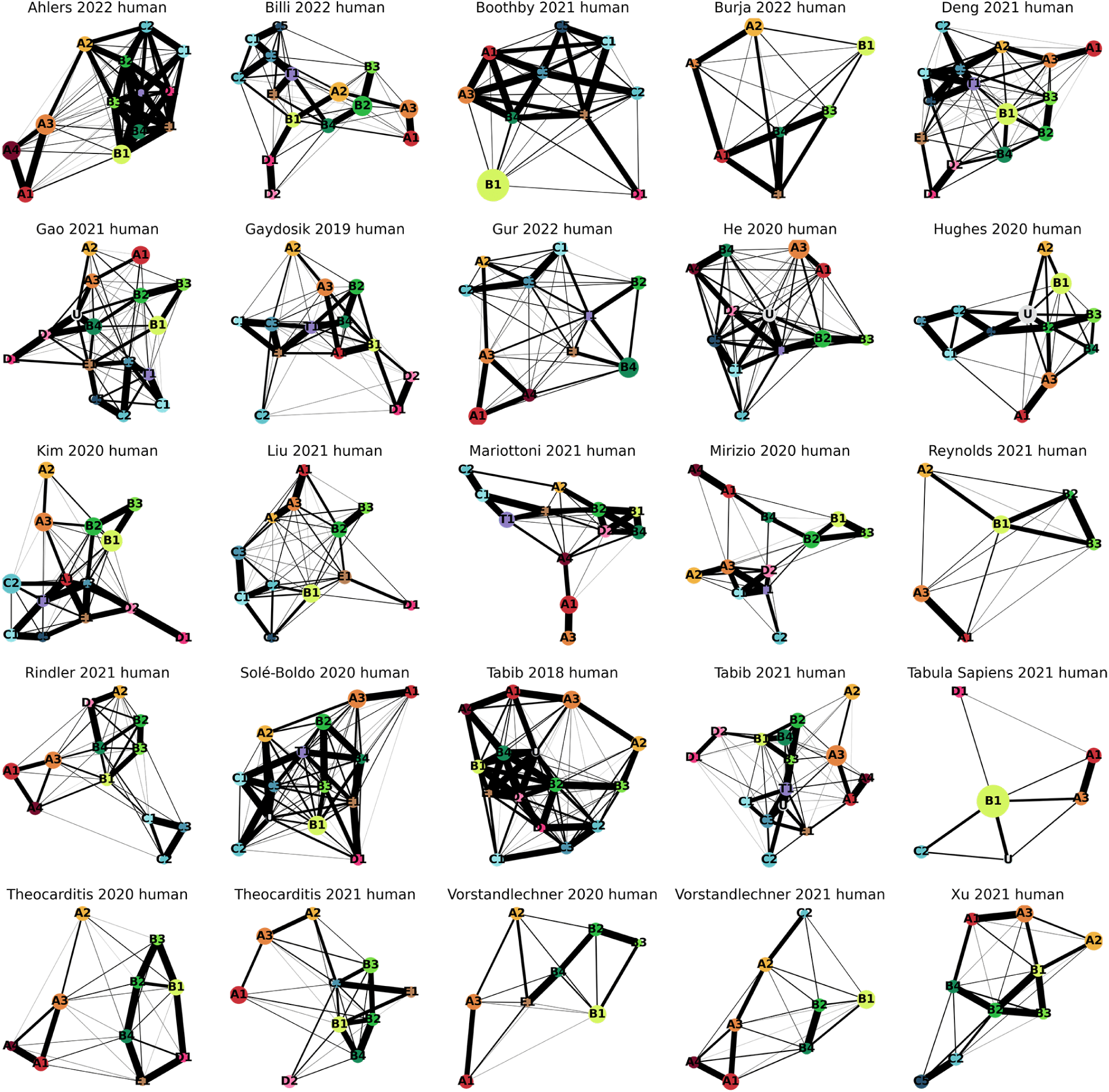
Full PAGA graph of human dermal populations. Colours of populations are shared across datasets. Thicker lines indicate a greater connectivity between nodes.

**Figure S4.**
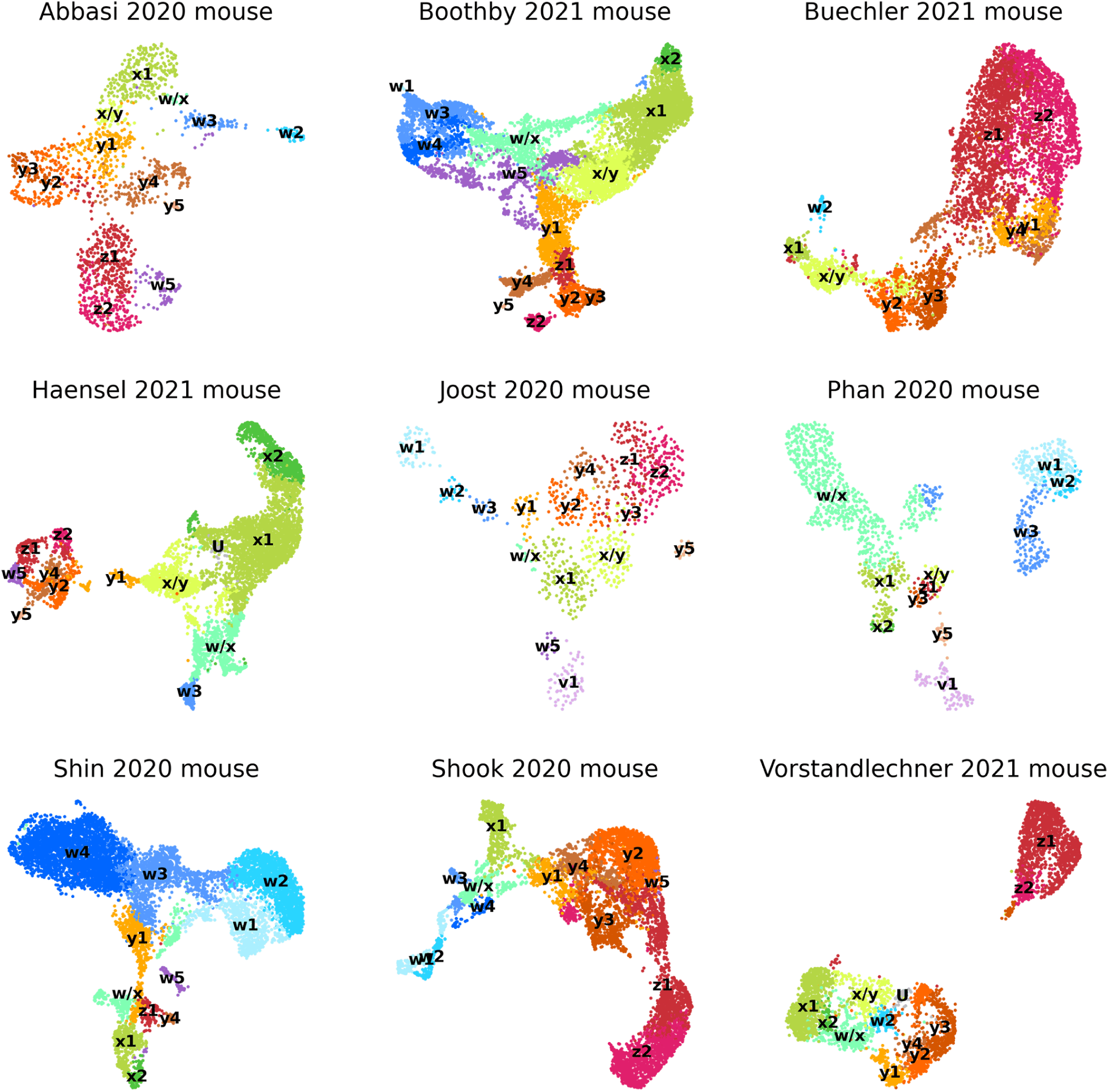
UMAP plots of all mouse fibroblast datasets from the secondary analysis. Colours of populations are shared across datasets.

**Figure S5.**
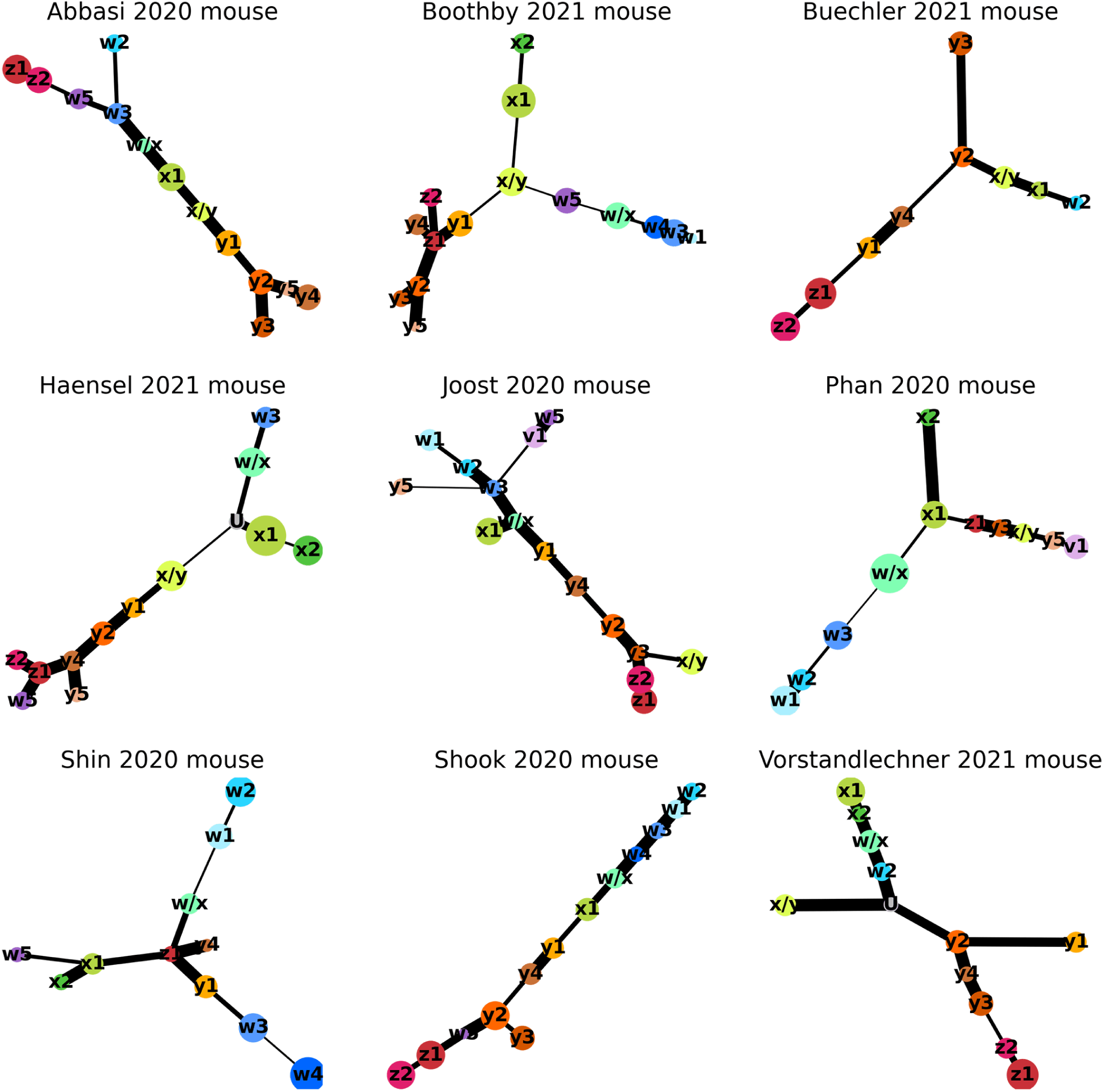
PAGA tree graph of mouse dermal populations. Colours of populations are shared across datasets. Thicker lines indicate greater connectivity between nodes.

**Figure S6.**
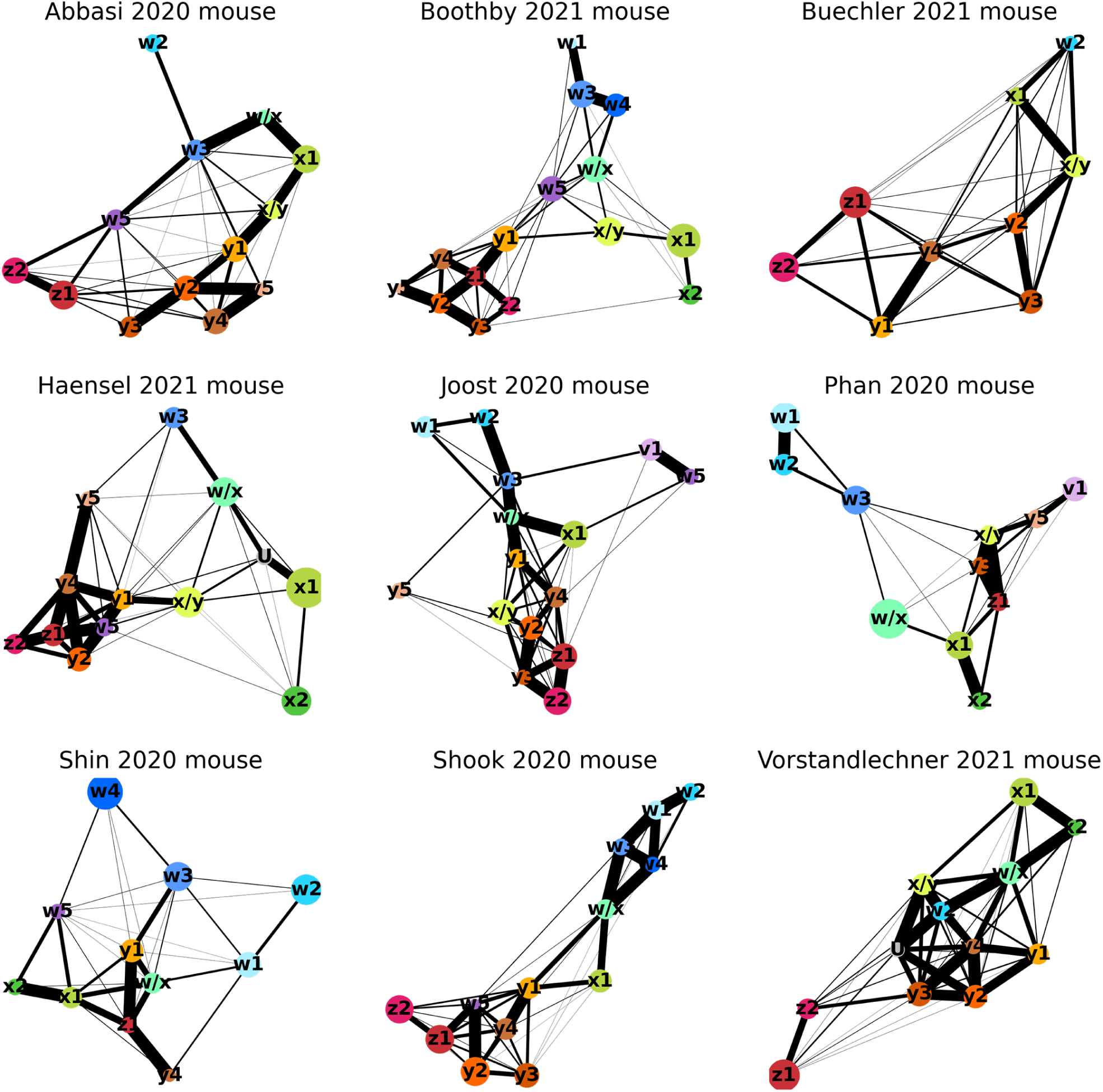
Full PAGA graph of mouse dermal populations. Colours of populations are shared across datasets. Thicker lines indicate a greater connectivity between nodes.

**Figure S7.**
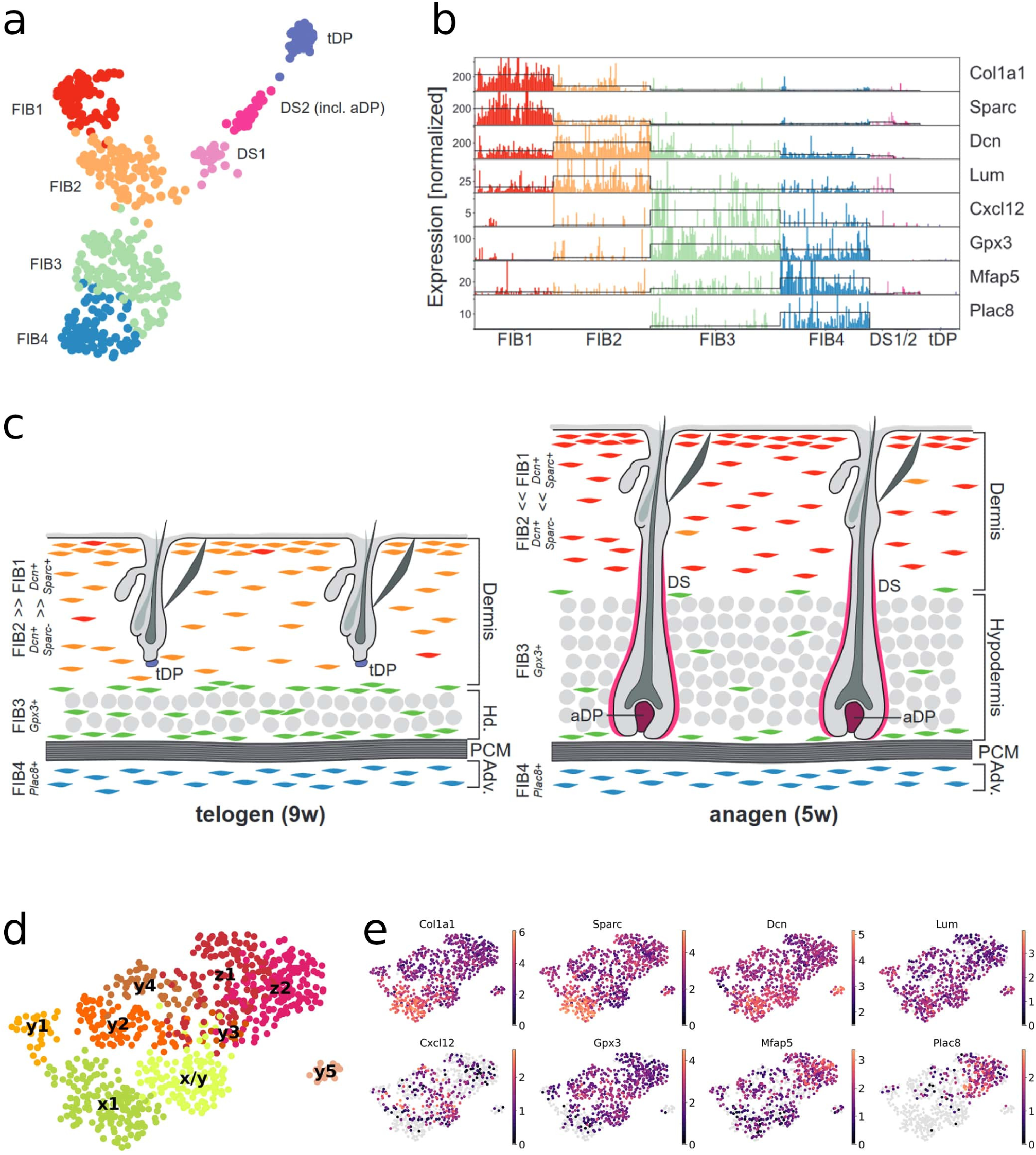
Mapping of Joost et al. (2020) populations. (a) UMAP plot of mouse dermal fibroblasts from Joost et al. (2020), adapted from Figure 6A of the publication. (b) Tracksplot of markers genes of populations FIB1 to FIB4, adapted from Figure 6B of the publication. (c) Scheme of the location of FIB1 to FIB4 populations in telogen and anagen phases, adapted from Figure 6I of the publication. (d) UMAP of dermal fibroblasts from Joost et al. (2020) after being reanalysed. Only cells from axes x, y and z are shown. (e) UMAP plots showing the expression levels of markers from subfigure (b). The naming of the nomenclature was inspired by the findings of (Joost et al., 2020). In their work, the authors analyze the composition of skin cells of hair follicles in anagen and telogen phases, including dermal fibroblasts. Regarding the fibroblast populations, they separate them into 7 populations: FIB1 to FIB4, DS1, DS2 and tDP. FIB populations are dermal fibroblasts, DS1 and DS2 are dermal sheath cells –in anagen and telogen phases–, and tDP are cells of the DP in telogen phase. Looking specifically at the location of the FIB populations using immunofluorescence, they observe FIB1 and FIB2 to be located within the dermis, where FIB1 has more presence during anagen and FIB2 during telogen; FIB3 is located within the hypodermis, and FIB4 in the adventitial layer beneath the *panniculus carnosus*–Figures 6F to 6H in the publication (Joost et al., 2020)–.

**Figure S8.**
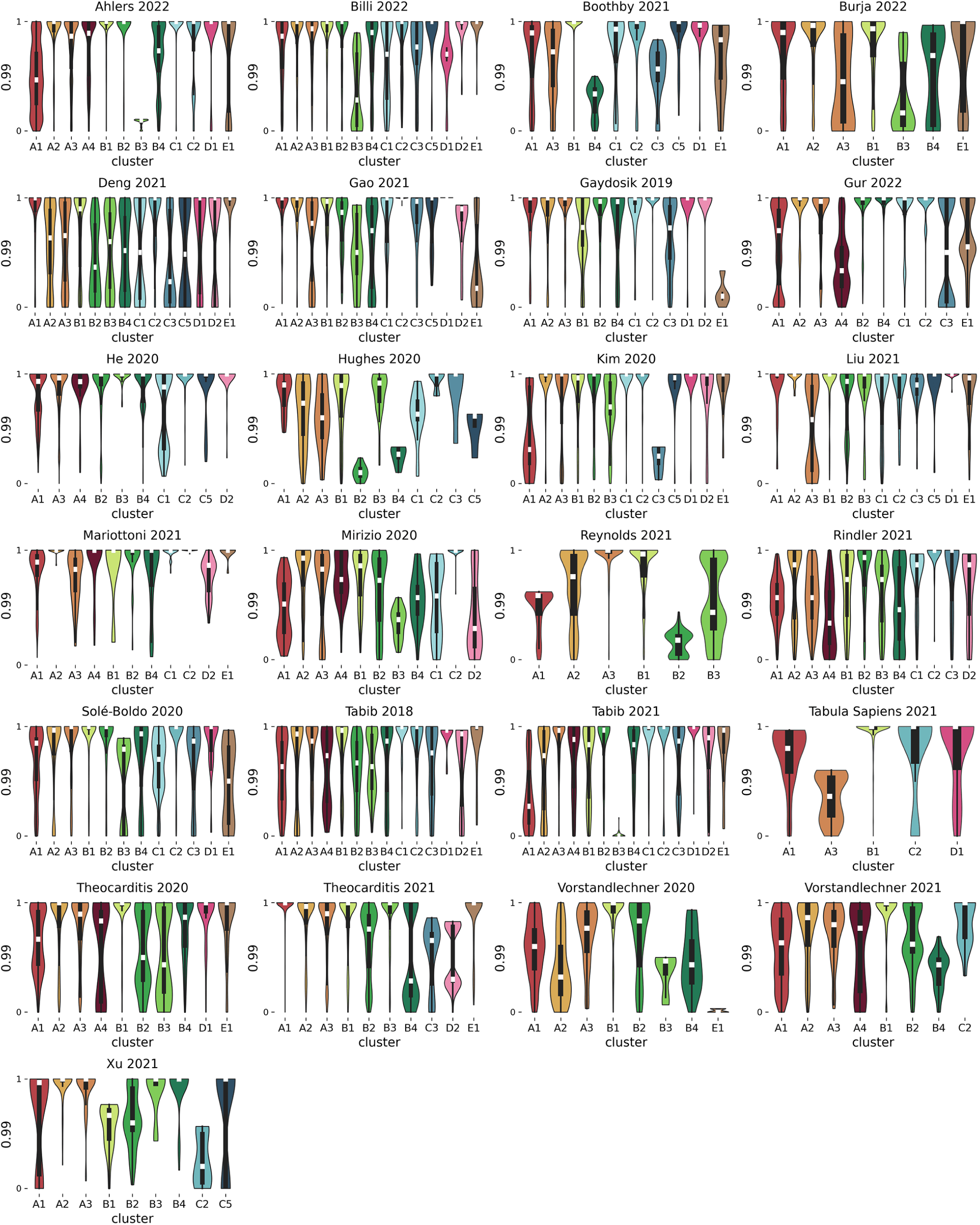
Violinplot of robustness score in human datasets. Each violin represents the distribution of robustness scores per dataset and population, across the representative cells. Boxes inside the violins represent the quartiles of the distribution, center bar represents the median, and whiskers extend to Q1 - 1.5 * IQR and Q3 + 1.5 * IQR.

**Figure S9.**
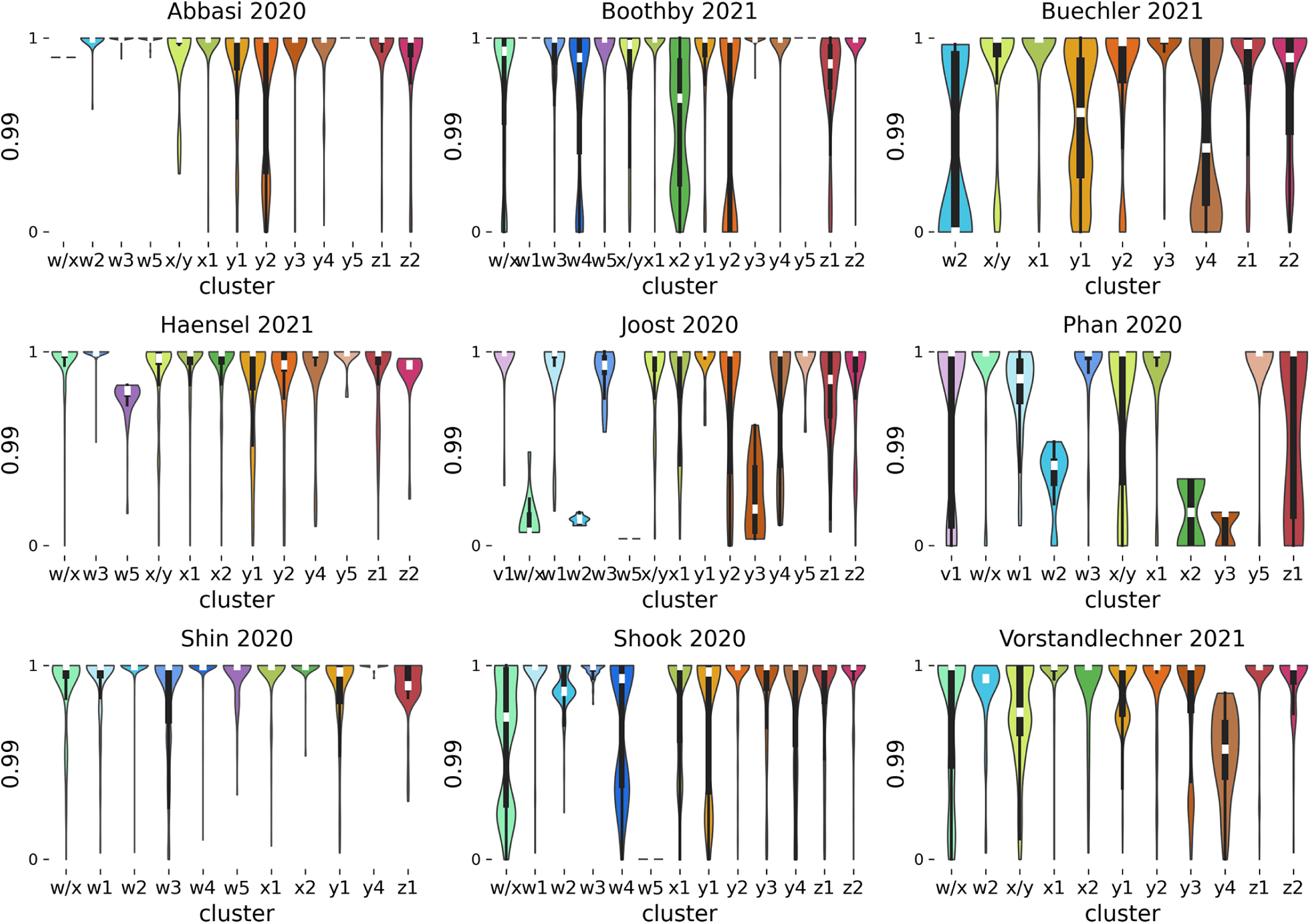
Violinplot of robustness score in mouse datasets. Each violin represents the distribution of robustness scores per dataset and population, across the representative cells. Boxes inside the violins represent the quartiles of the distribution, center bar represents the median, and whiskers extend to Q1 - 1.5 * IQR and Q3 + 1.5 * IQR.

**Figure S10.**
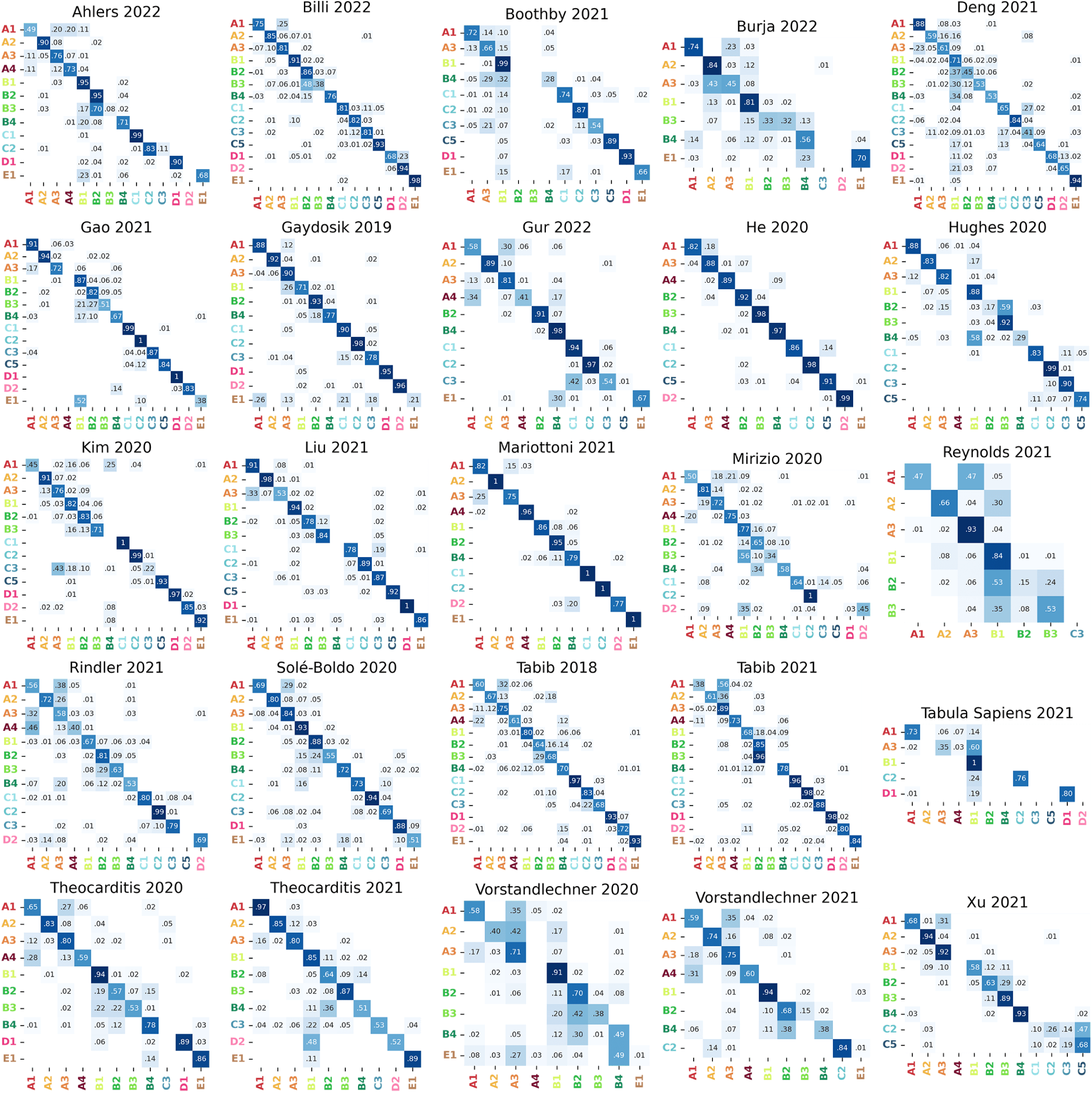
Adjacency matrix of robust cluster assignation in human datasets. Each row-column combination is the proportion of times the row cluster is assigned to the column cluster, considering all cells within the row clusters, and all the iterations.

**Figure S11.**
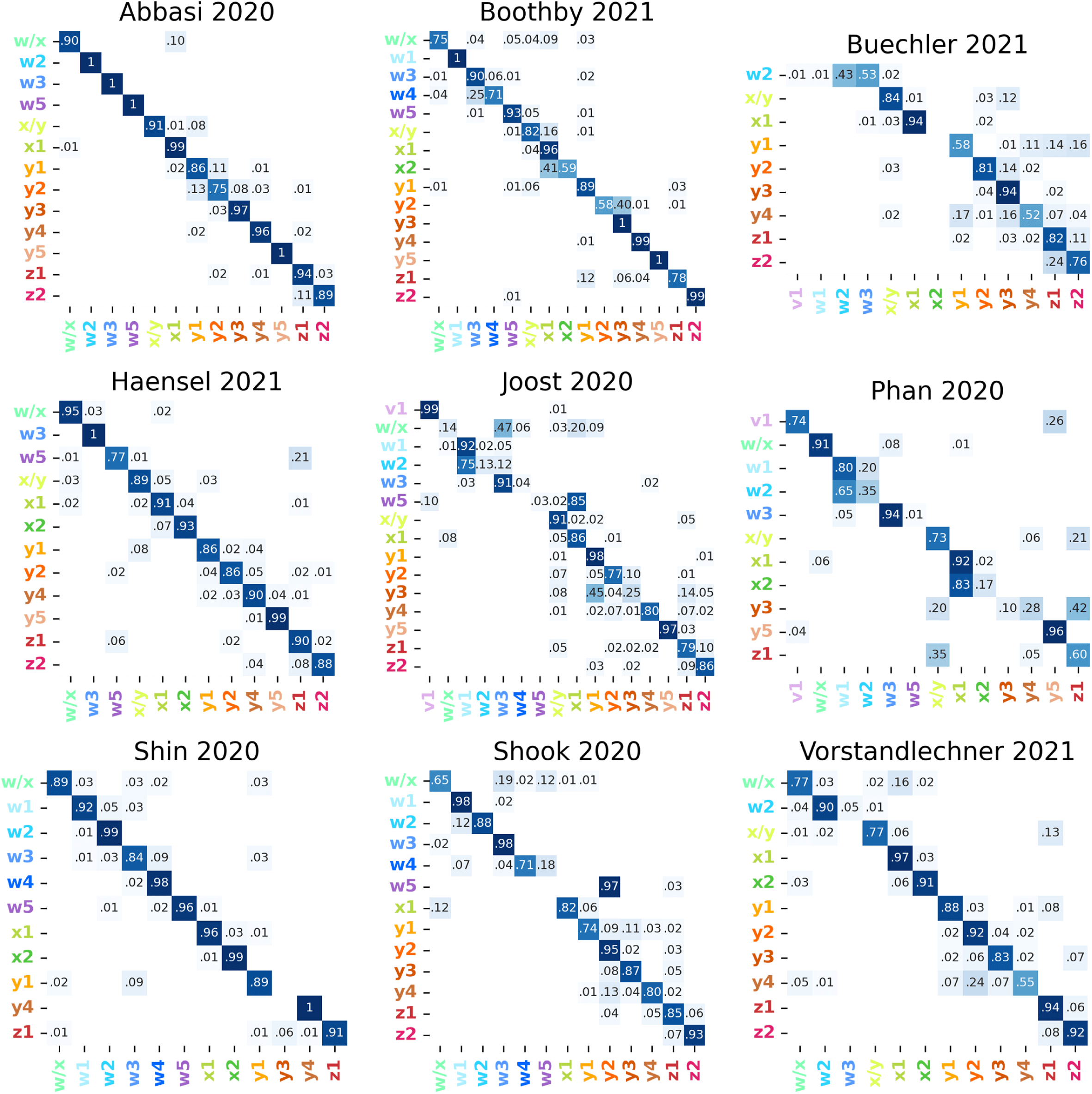
Adjacency matrix of robust cluster assignation in mouse datasets. Each row-column combination is the proportion of times the row cluster is assigned to the column cluster, considering all cells within the row clusters, and all the iterations.

**Figure S12.**
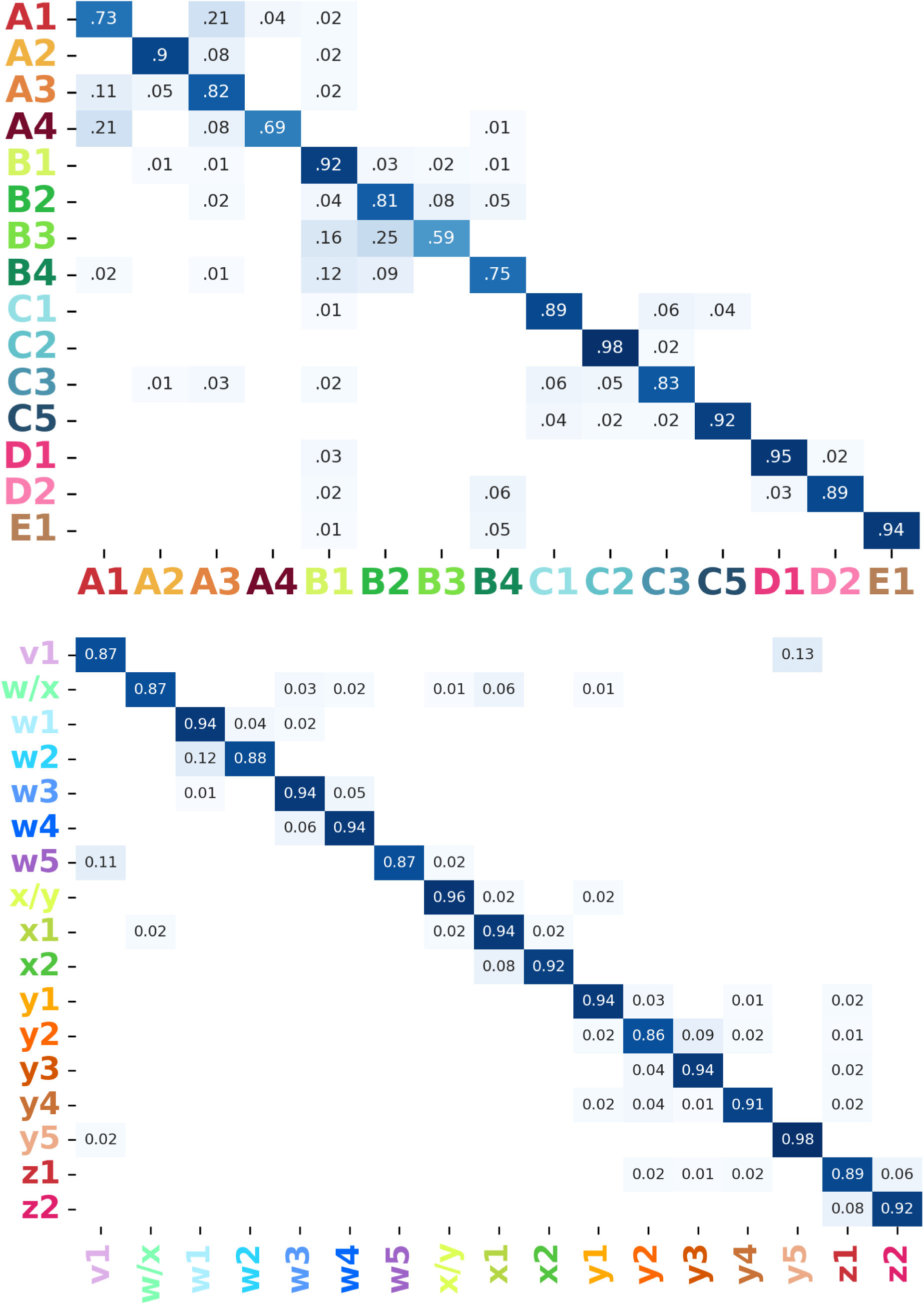
Adjacency matrix of robust cluster assignation. Each row-column combination is the proportion of times the row cluster is assigned to the column cluster. In this heatmap, numbers represent the median of the proportions across datasets.

**Figure S13.**
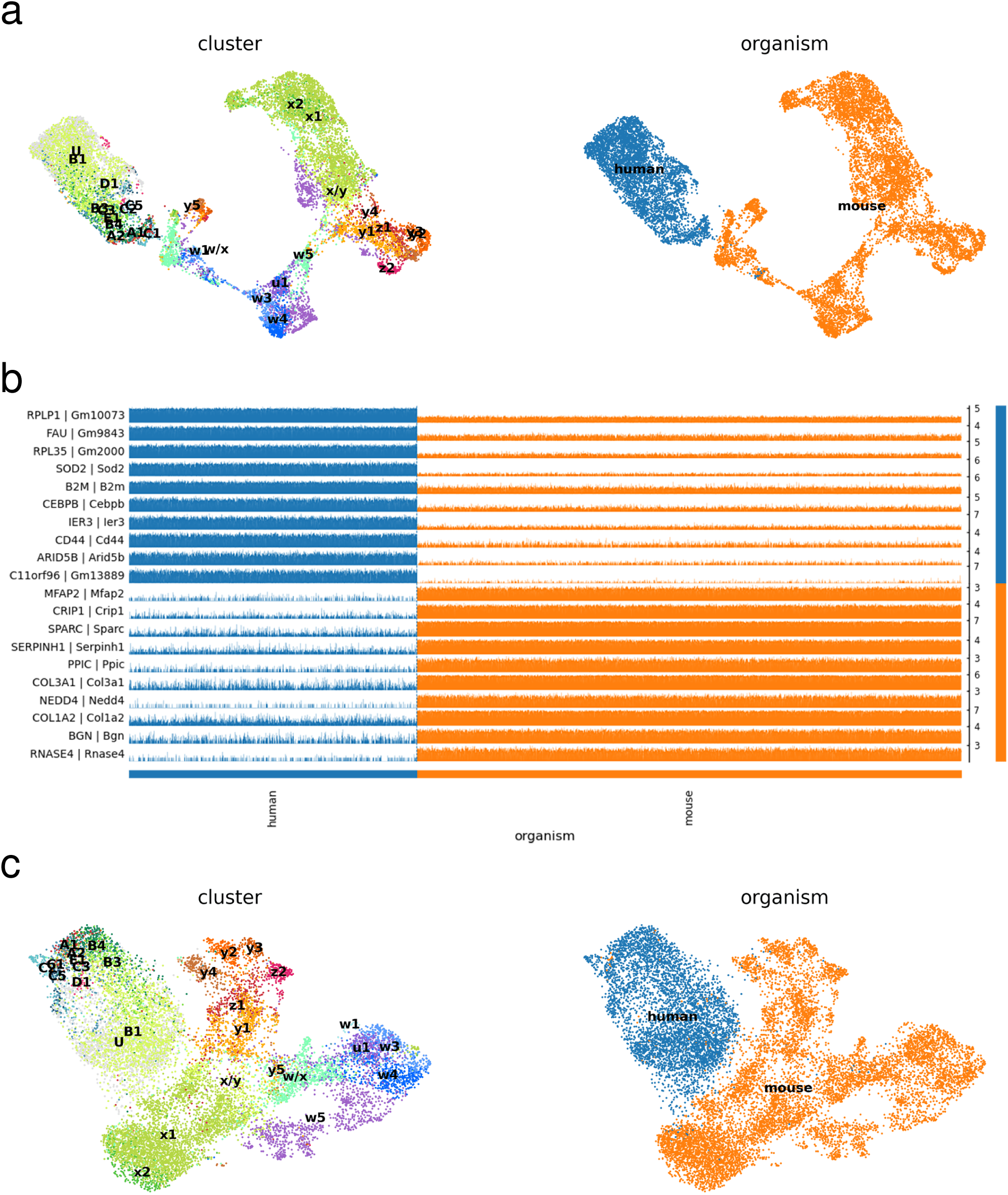
Analysis of mouse/human integration on Boothby et al.. (a) UMAP plots of clusters and organism of Boothby et al. dataset, using only harmony_integrate function. (b) First 10 DEGs between human and mouse populations. (c) UMAP plots of clusters and organisms of (Boothby et al., 2021) dataset, using regress_out and harmony_integrate functions.

**Figure S14.**
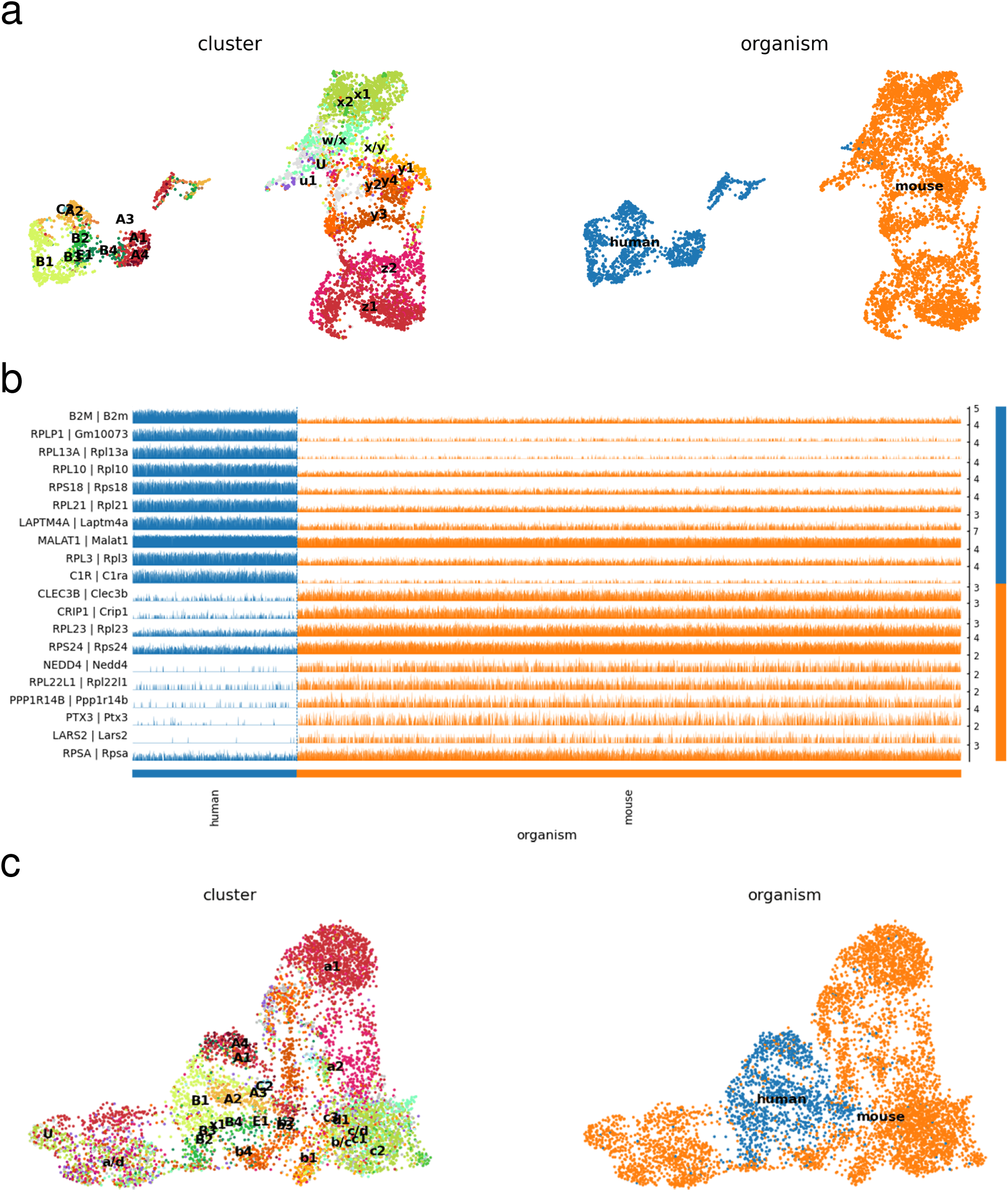
Analysis of mouse/human integration on (Vorstandlechner et al., 2021). (a) UMAP plots of clusters and organism of (Vorstandlechner et al., 2021) dataset, using only harmony_integrate function. (b) First 10 DEGs between human and mouse populations. (c) UMAP plots of clusters and organism of (Vorstandlechner et al., 2021) dataset, using regress_out and harmony_integrate functions.

**Figure S15.**
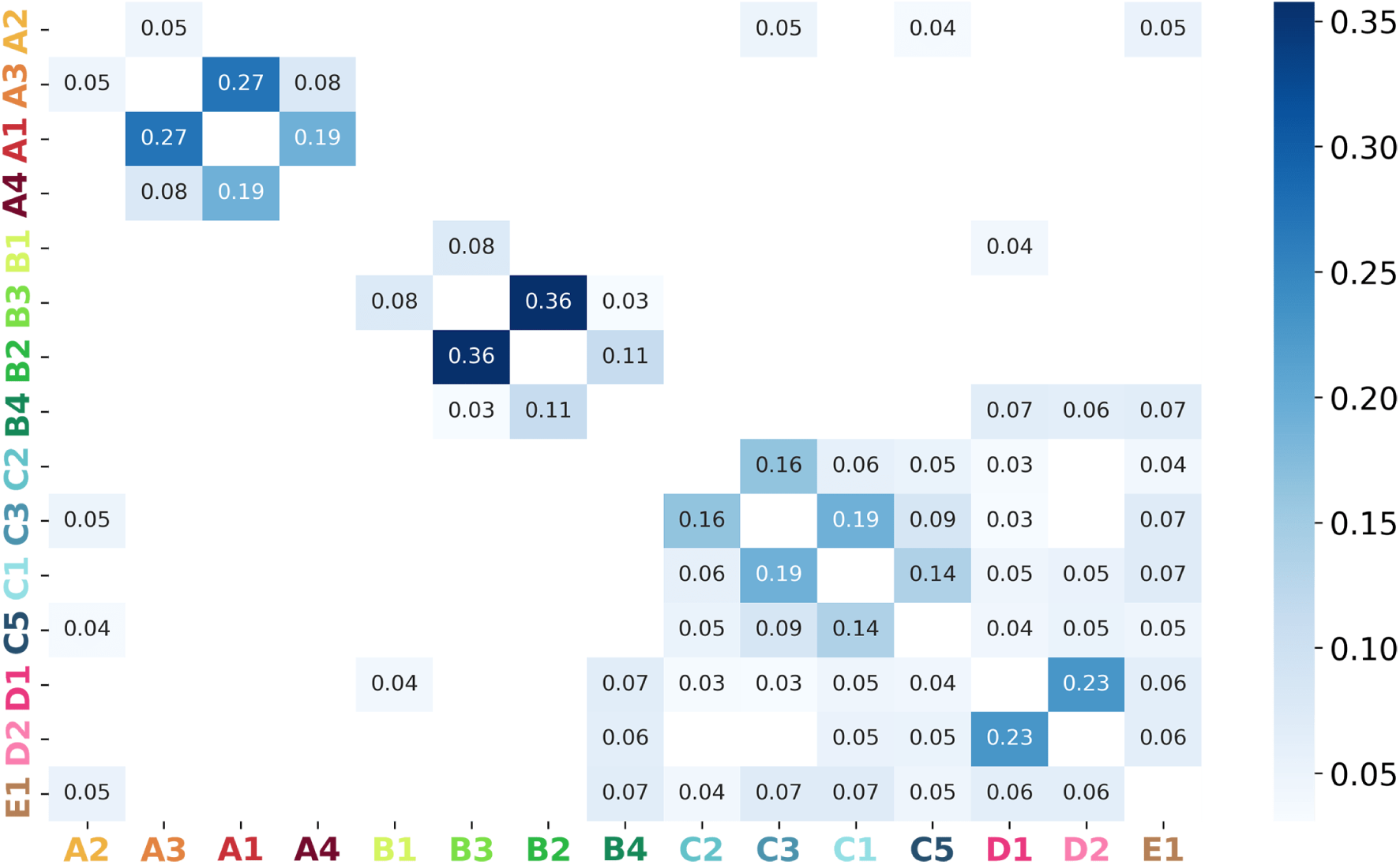
Heatmap of human marker overlap across populations. For each pair of populations, the number in the cell represents the proportion of overlapping markers.

**Figure S16.**
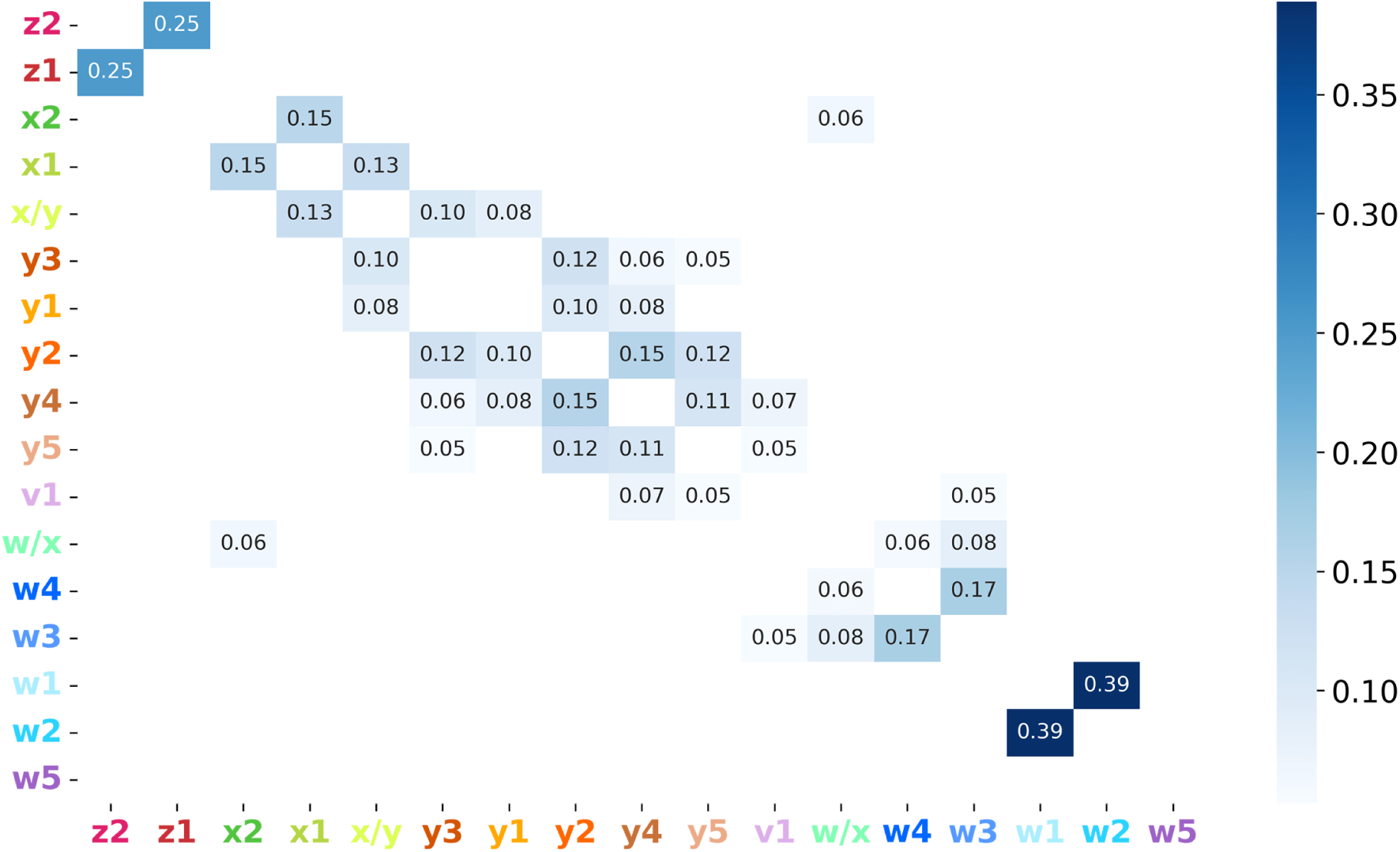
Heatmap of mouse marker overlap across populations. For each pair of populations, the number in the cell represents the proportion of overlapping markers.

**Figure S17.**
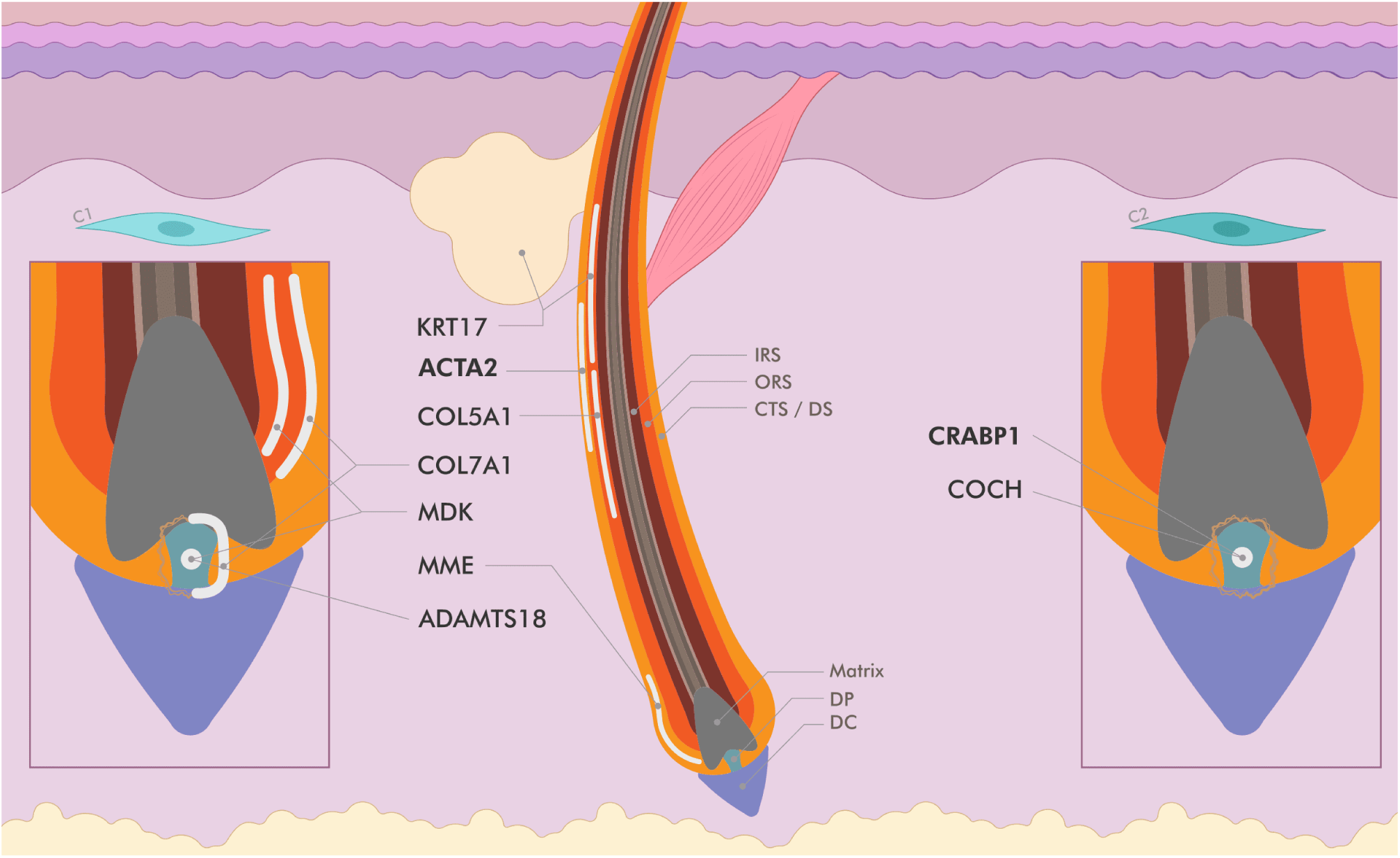
Location of proteins expressed by C1 and C2 fibroblasts. Schematic drawing of the HF showing an approximate expression pattern of reported proteins from the literature.

**Figure S18.**
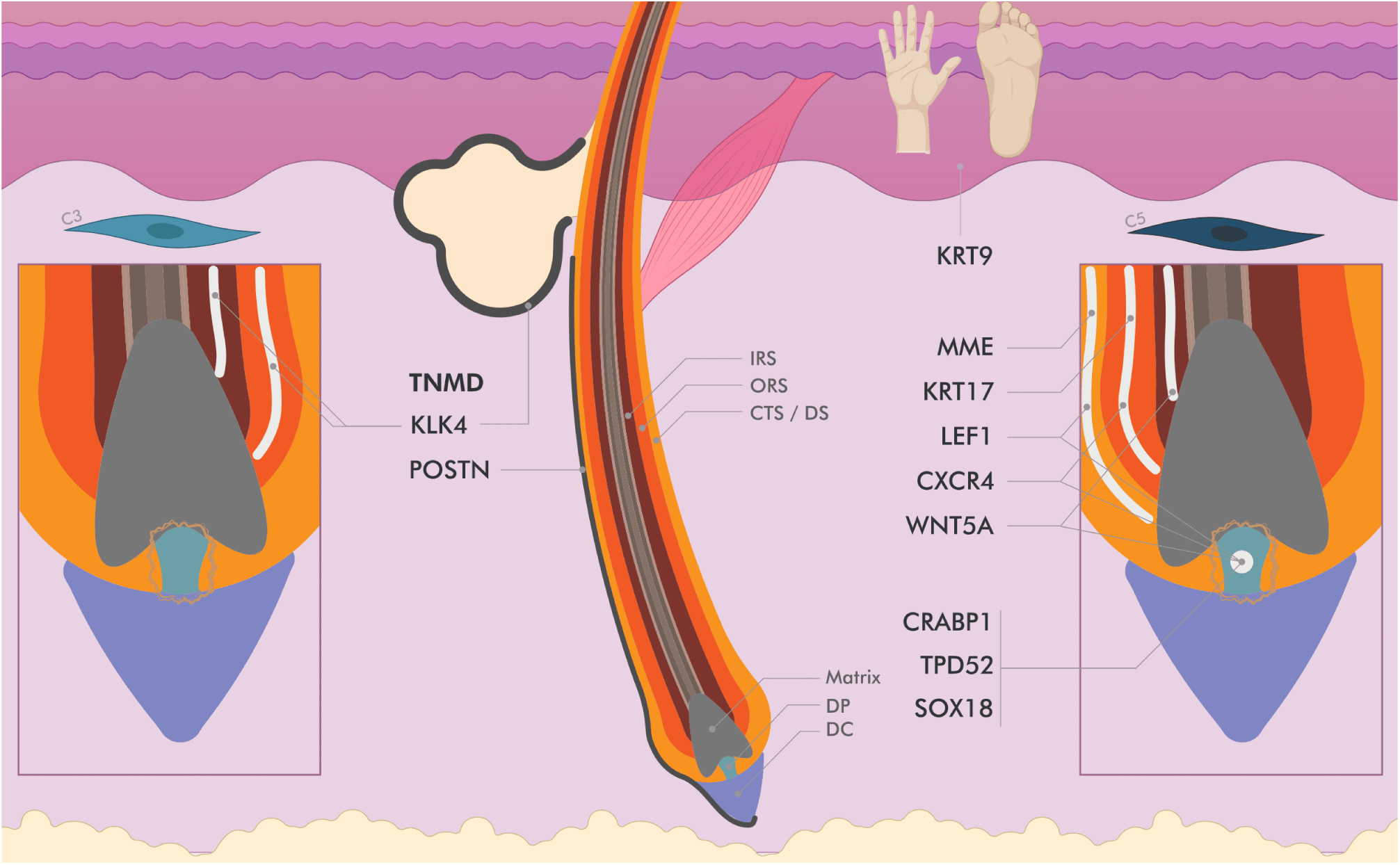
Location of proteins expressed by C3 and C5 fibroblasts. Schematic drawing of the HF showing an approximate expression pattern of reported proteins from the literature.

**Figure S19.**
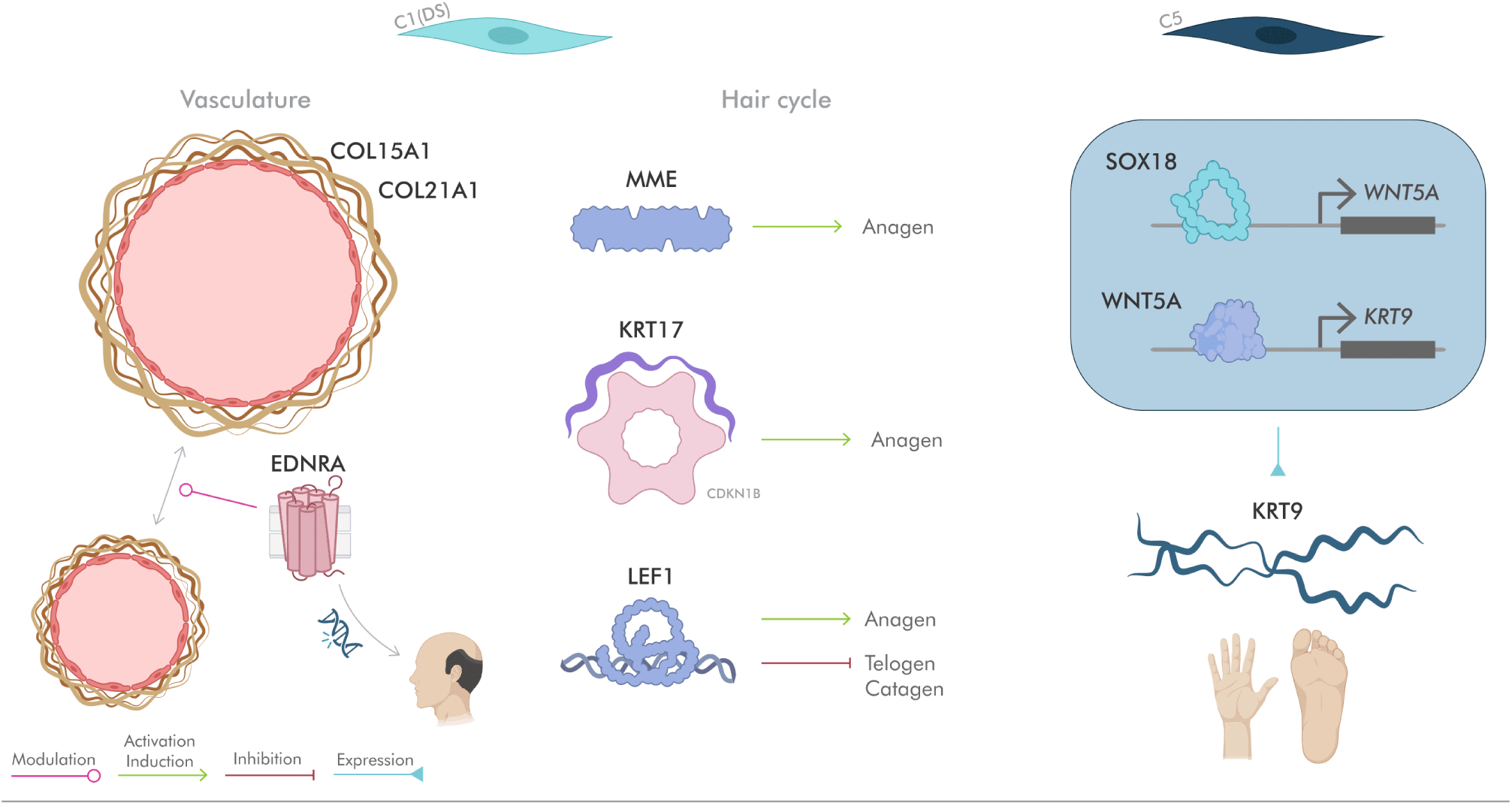
Specific functions of populations C1 and C5. Simplified scheme of specific functions of C1 and C5 populations, namely vasculature and HF cycle for C1 and *KRT9* expression for C5.

**Figure S20.**
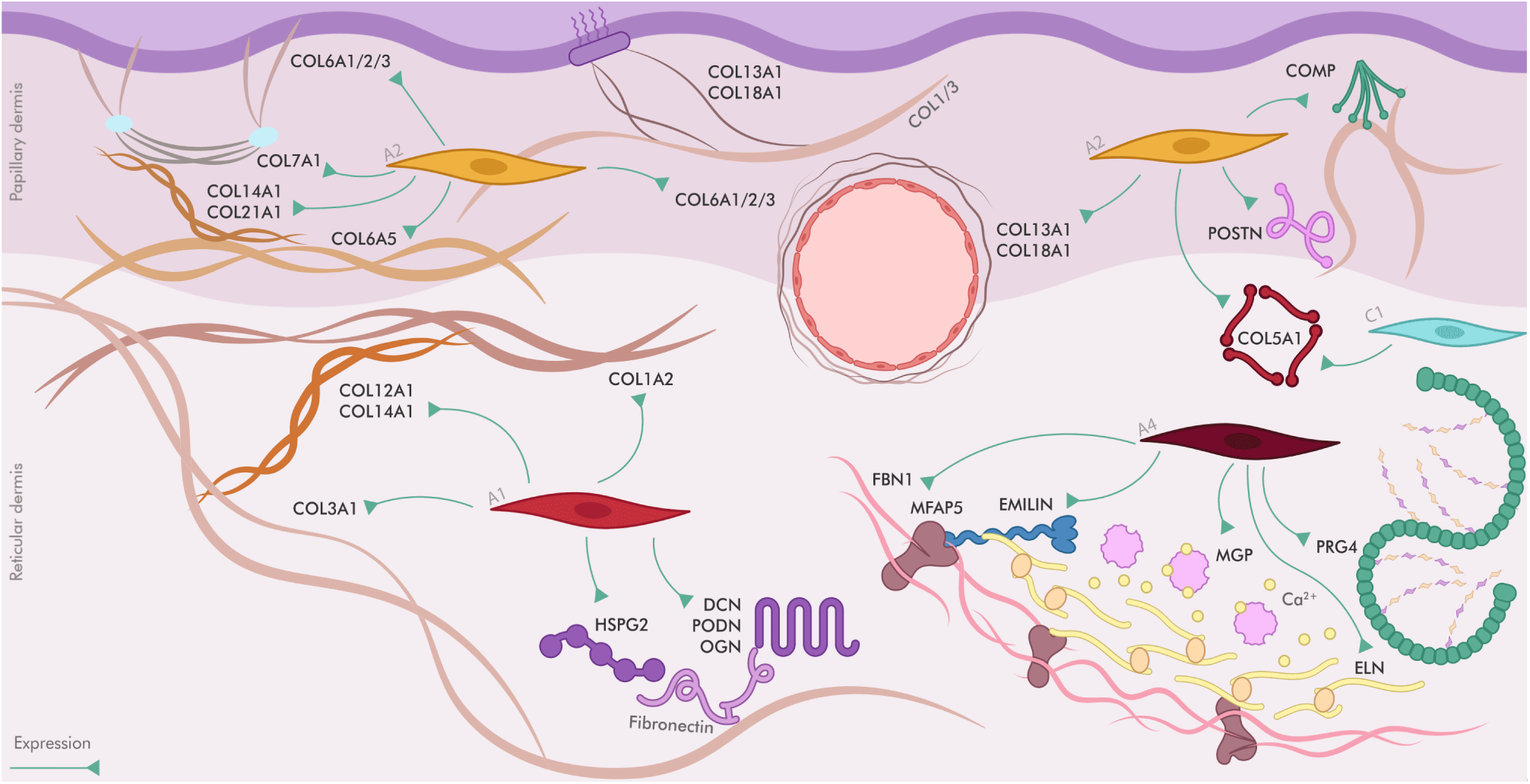
Expression of ECM components by type A fibroblasts. Scheme of location and expression of ECM components by type A fibroblasts. Components written in gray are not putatively secreted by these fibroblasts, but do interact with the former components.

**Figure S21.**
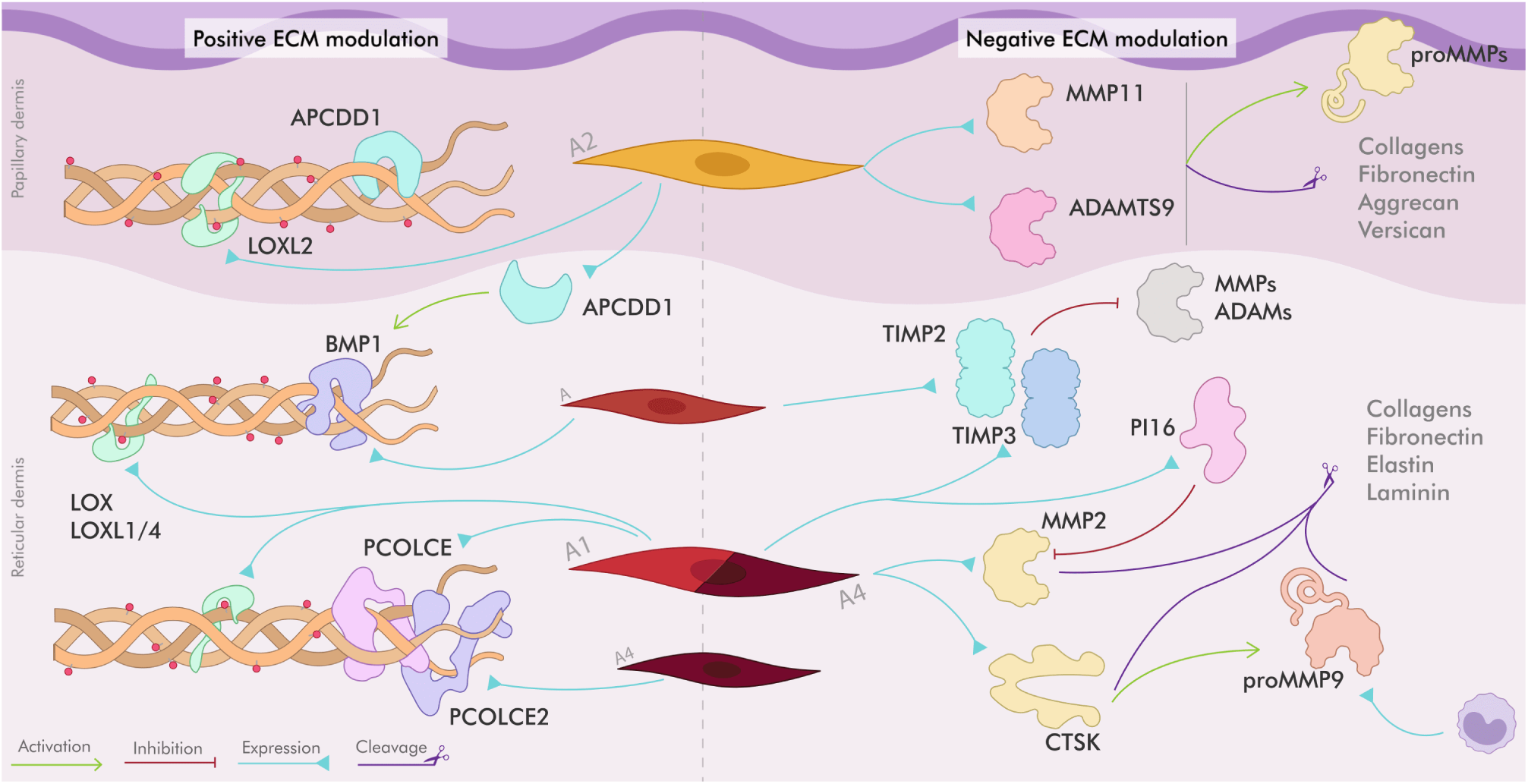
ECM modulation by type A fibroblasts. Scheme of functions of secreted components of type A fibroblast on the modulation of ECM.

**Figure S22.**
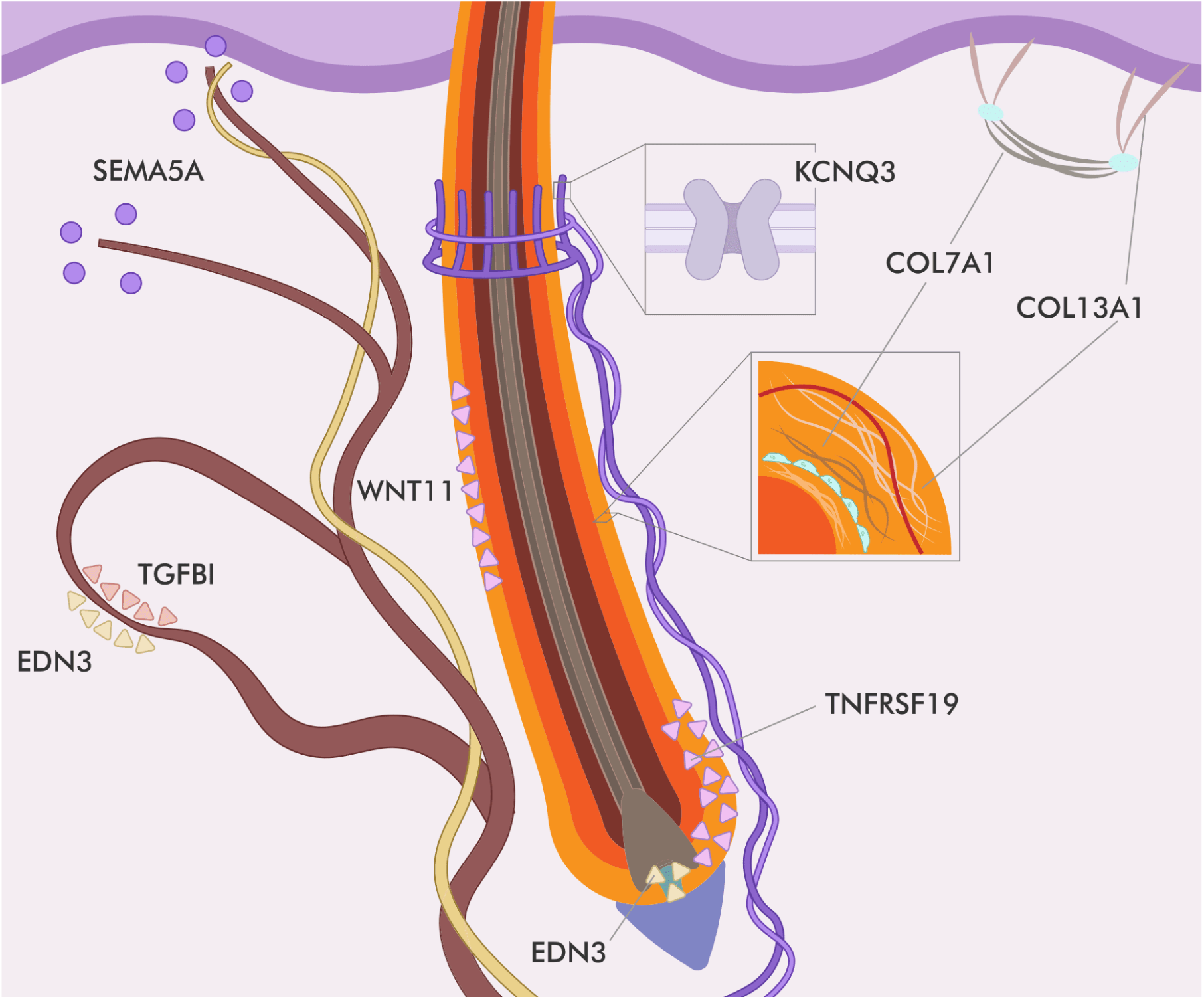
A2 fibroblast function in HF and other structures. Scheme of location and interaction of components secreted by A2 type fibroblasts in HF (KNCQ3, COL7A1, COL13A1, TNFRSF19, WNT11 and EDN3), nerve (SEMA5A) and blood vessel (TGFBI and EDN3).

**Figure S23.**
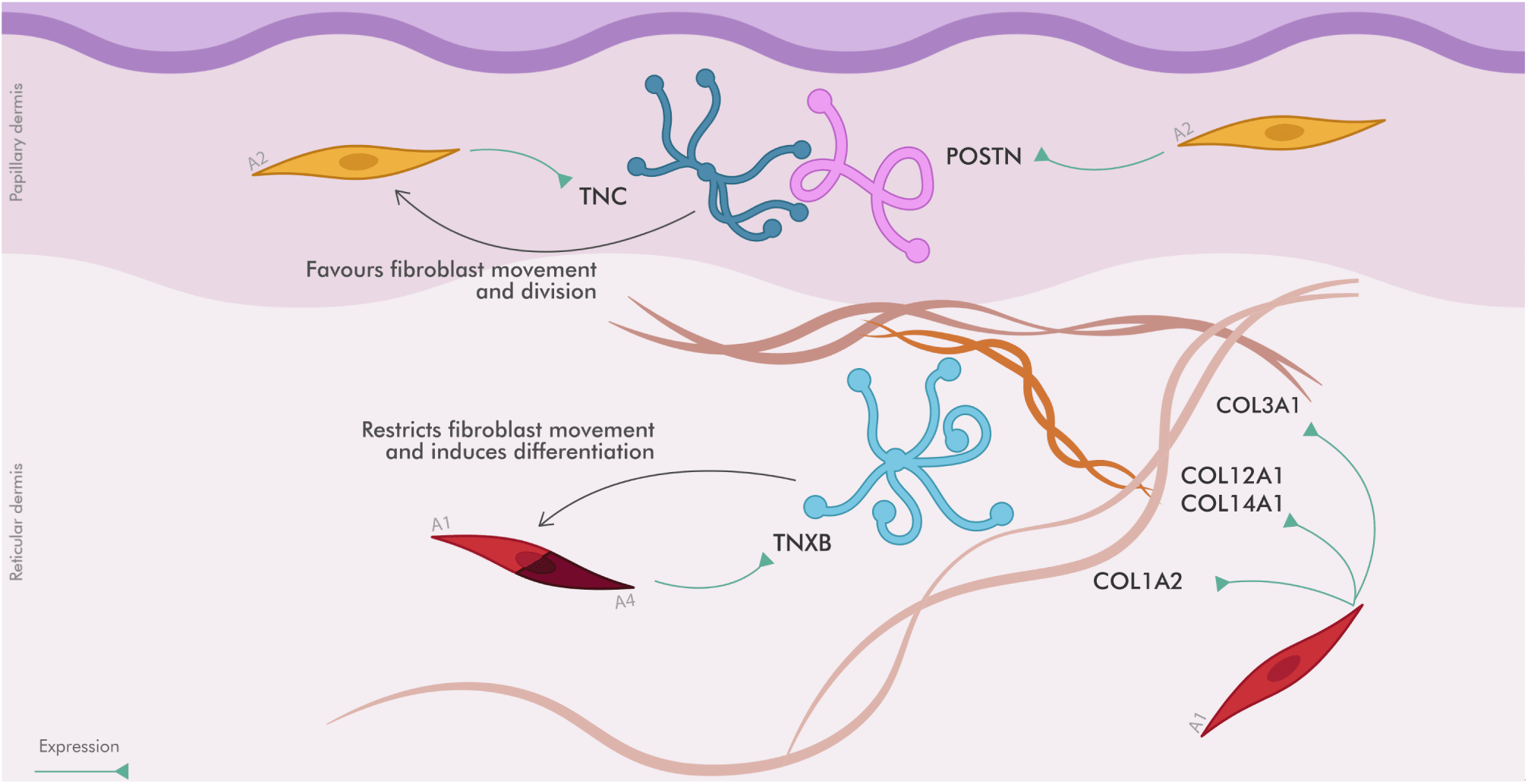
TNC and TNXB interaction scheme. Scheme of interaction of TNC and TNXB with A2- and A1-secreted components respectively.

**Figure S24.**
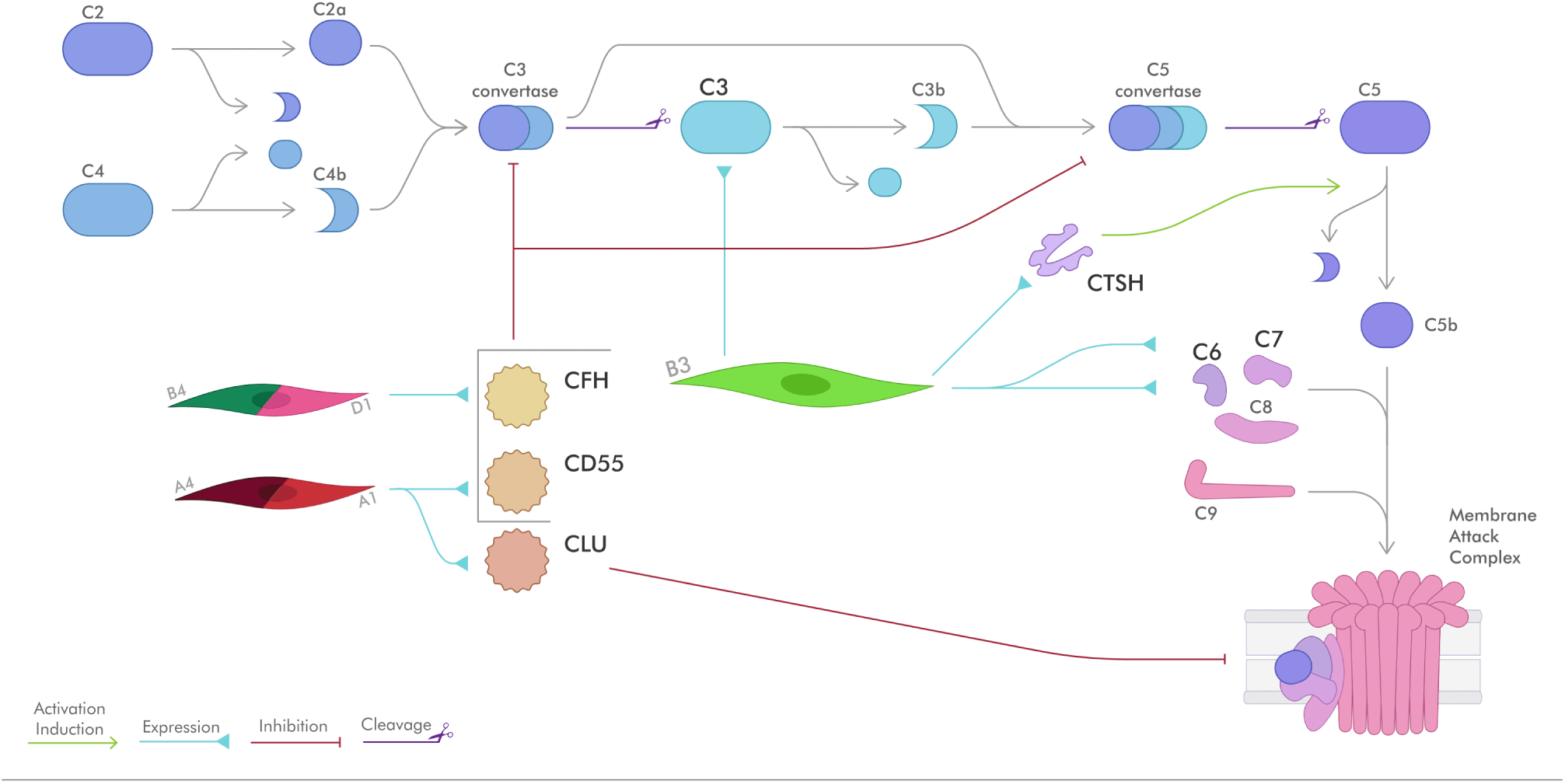
Interaction of fibroblast-expressed proteins with the complement system. Scheme of the classical complement pathway and its interaction with fibroblast-secreted proteins. These proteins are marked in a bolder typeface (CFH, CD55, CLU, C3, CTSH, C6, and C7).

**Figure S25.**
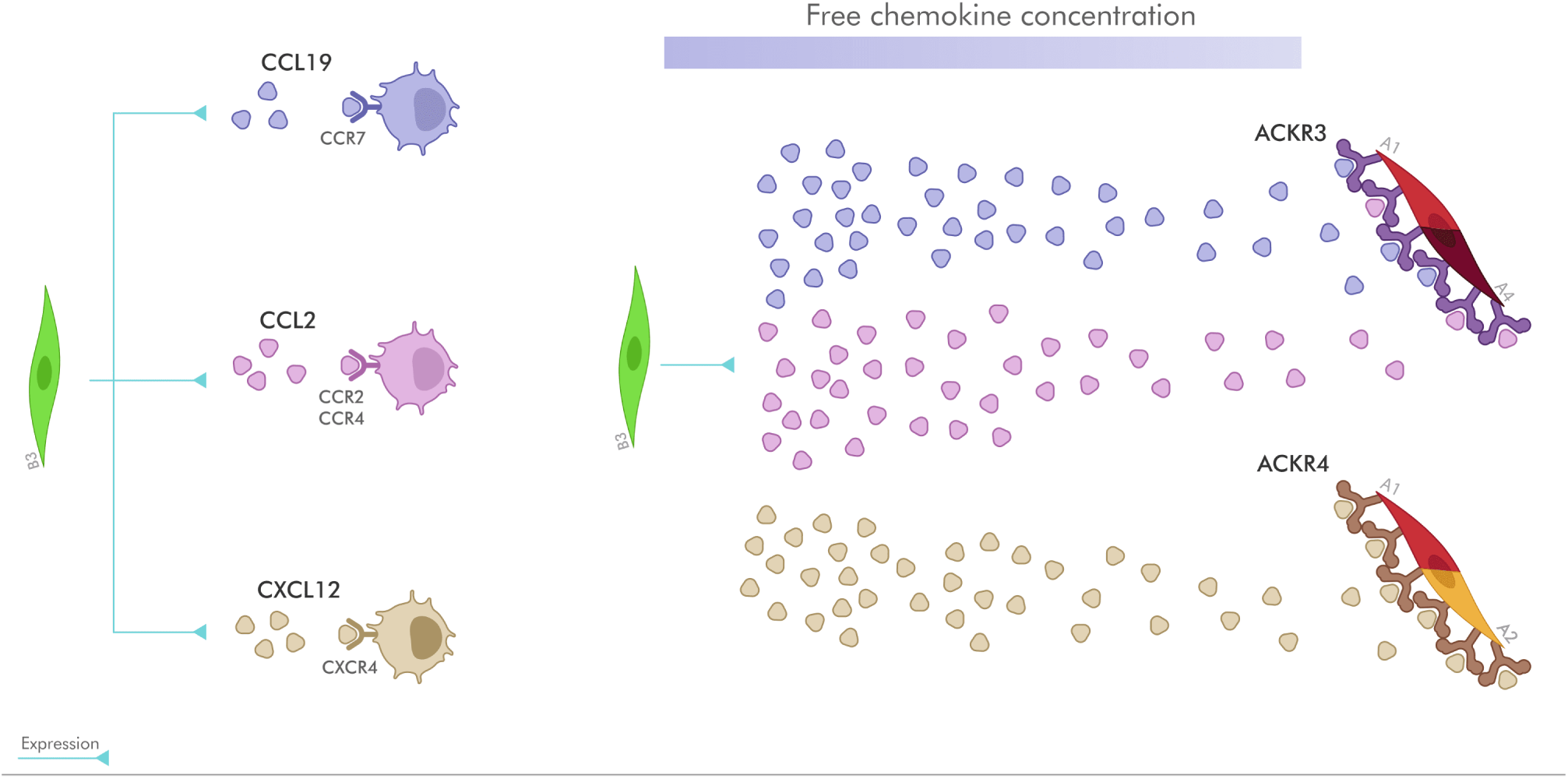
Chemokine modulation by ACKRs. Scheme of modulatory effect of ACKR3 and ACKR4 on B3-expressed CCL2, CCL19 and CXCL12 chemokines. ACKRs act as scavengers and can (1) create a chemokine gradient and (2) clear up chemokines from the environment.

**Figure S26.**
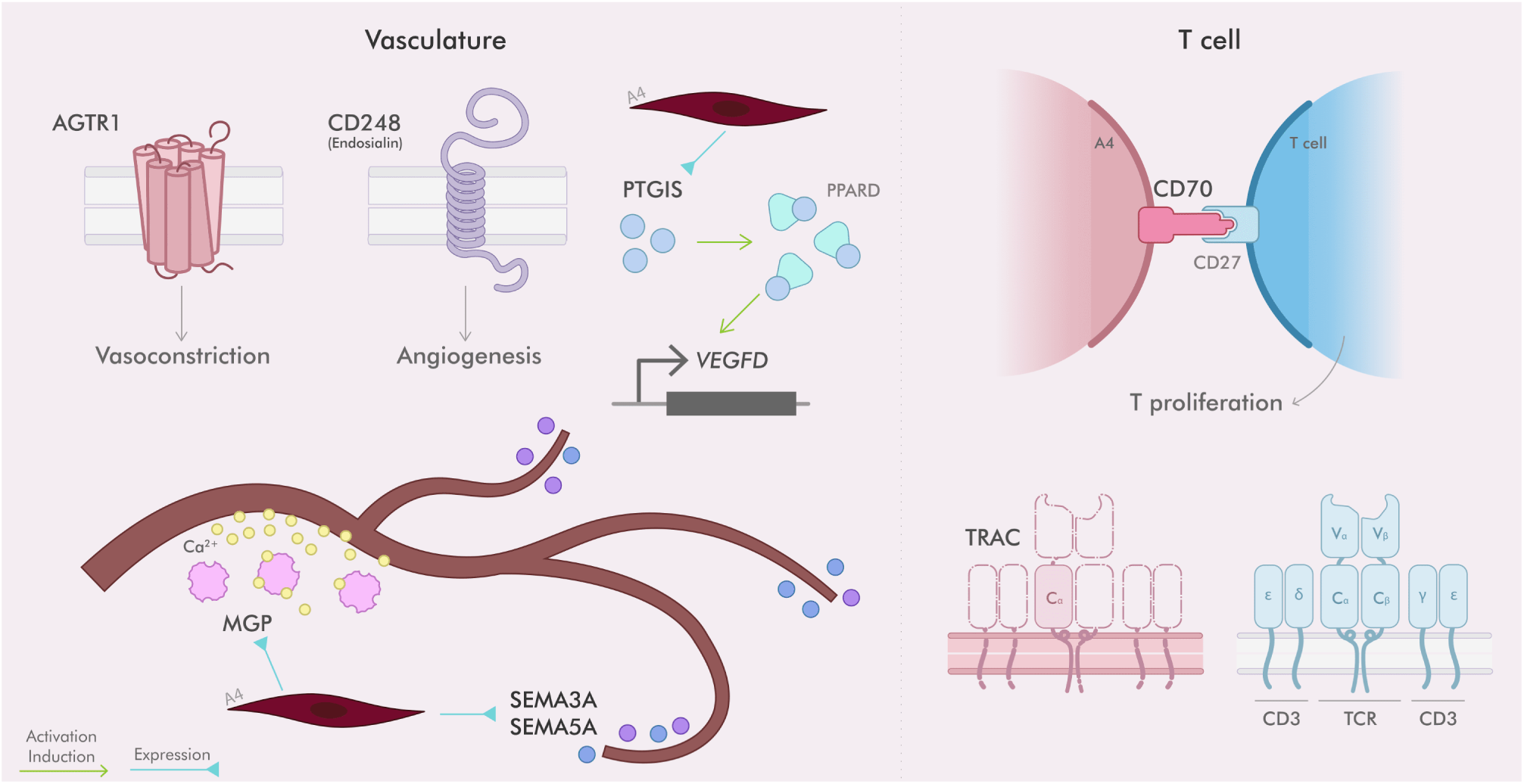
Specific functions of A4 type fibroblasts. Scheme of diverse functions from expressed markers, including possible interactions with the vasculature, and an unexplored possible interaction with T cells.

**Figure S27.**
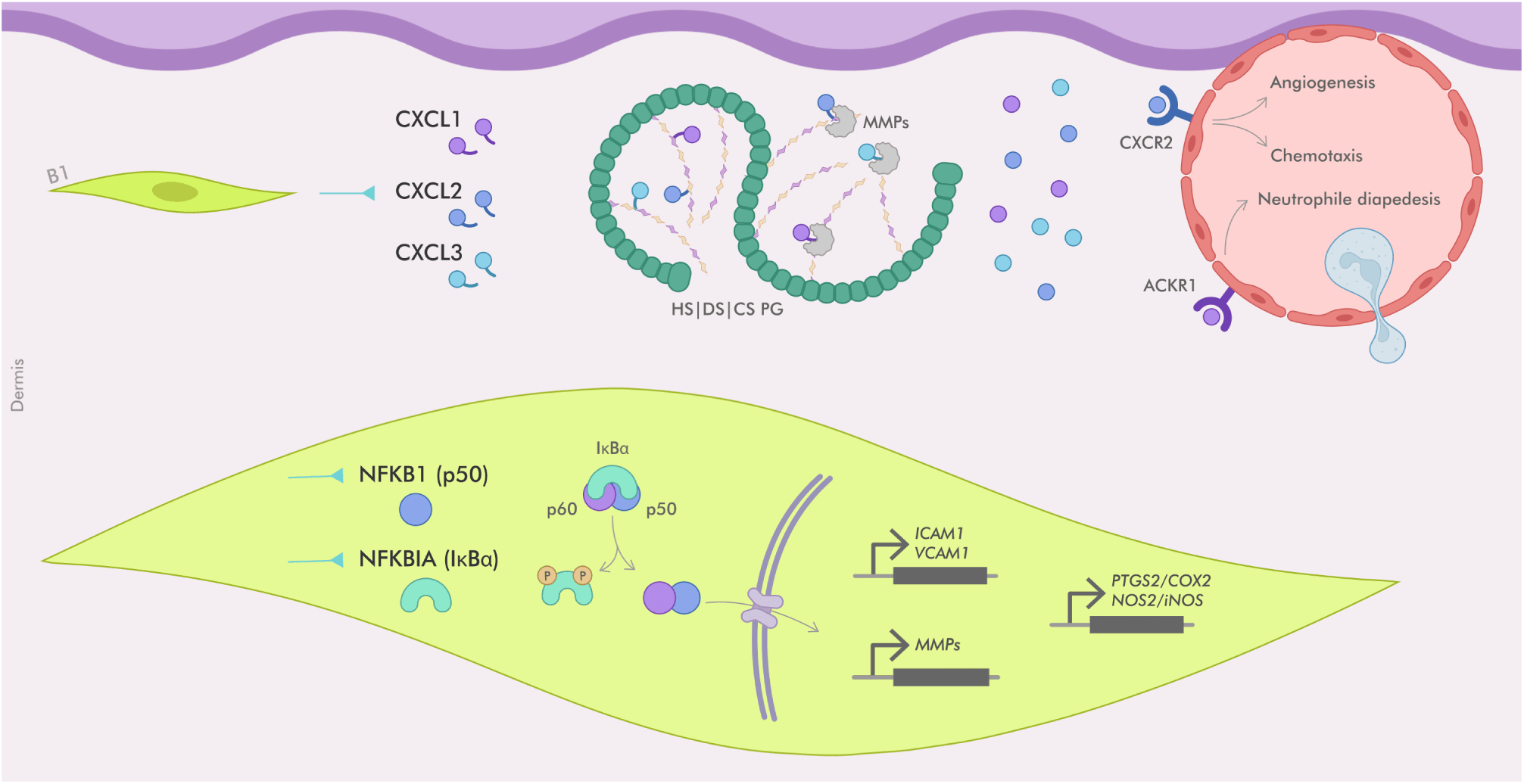
Mechanisms of acute phase chemoattraction of B1 fibroblasts. Mechanisms mediated by CXCL1/2/3 (up) and by NFKB1 and NFKBIA (down). Components with smaller font type and in gray may not be expressed by B1 fibroblasts, but interact with the other components.

**Figure S28.**
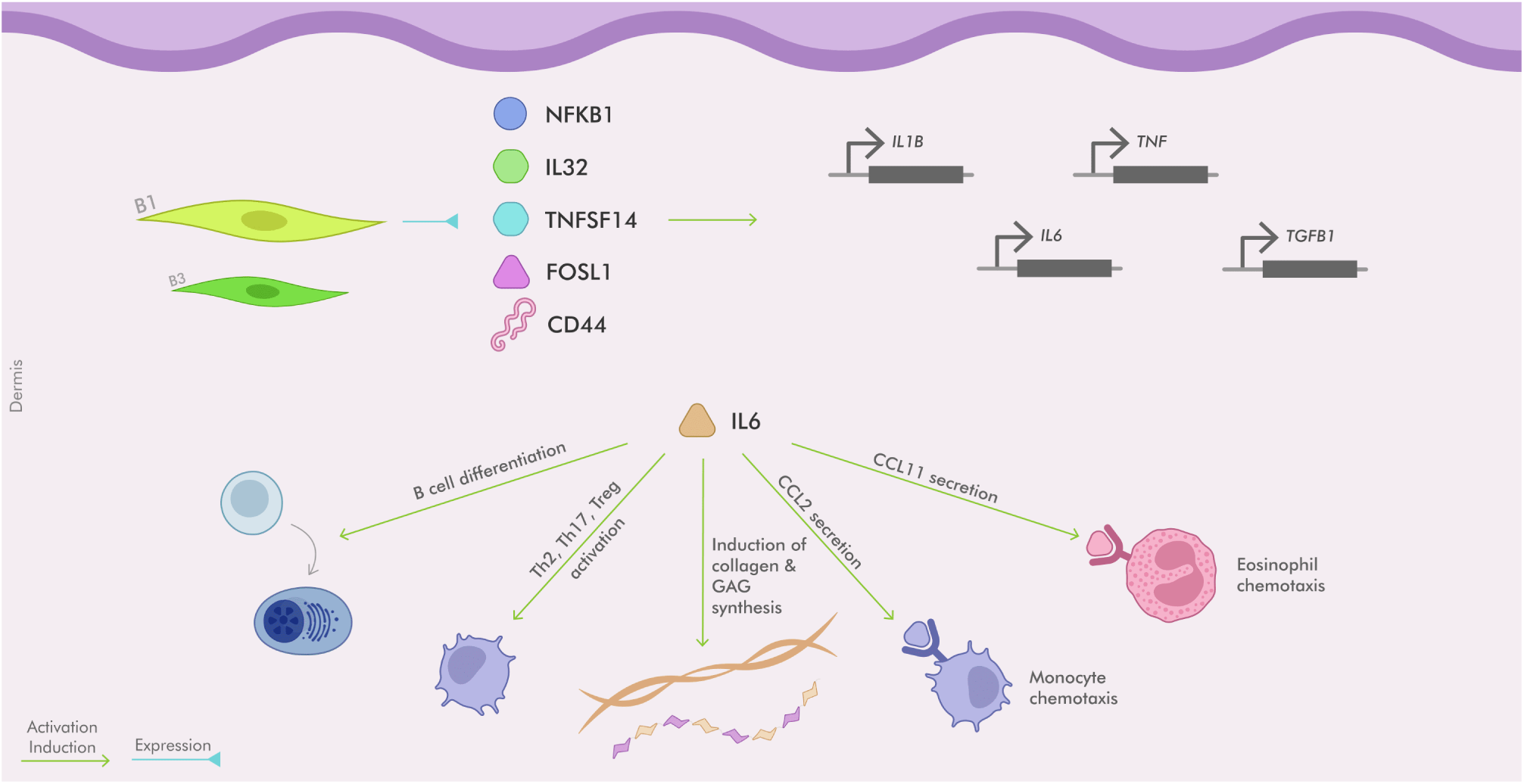
Acute phase chemokines secreted by B1 fibroblasts. Chemokines secreted by B1 fibroblasts and their relationshi with activation of downstream genes (up) and the functions related to IL6 (down).

**Figure S29.**
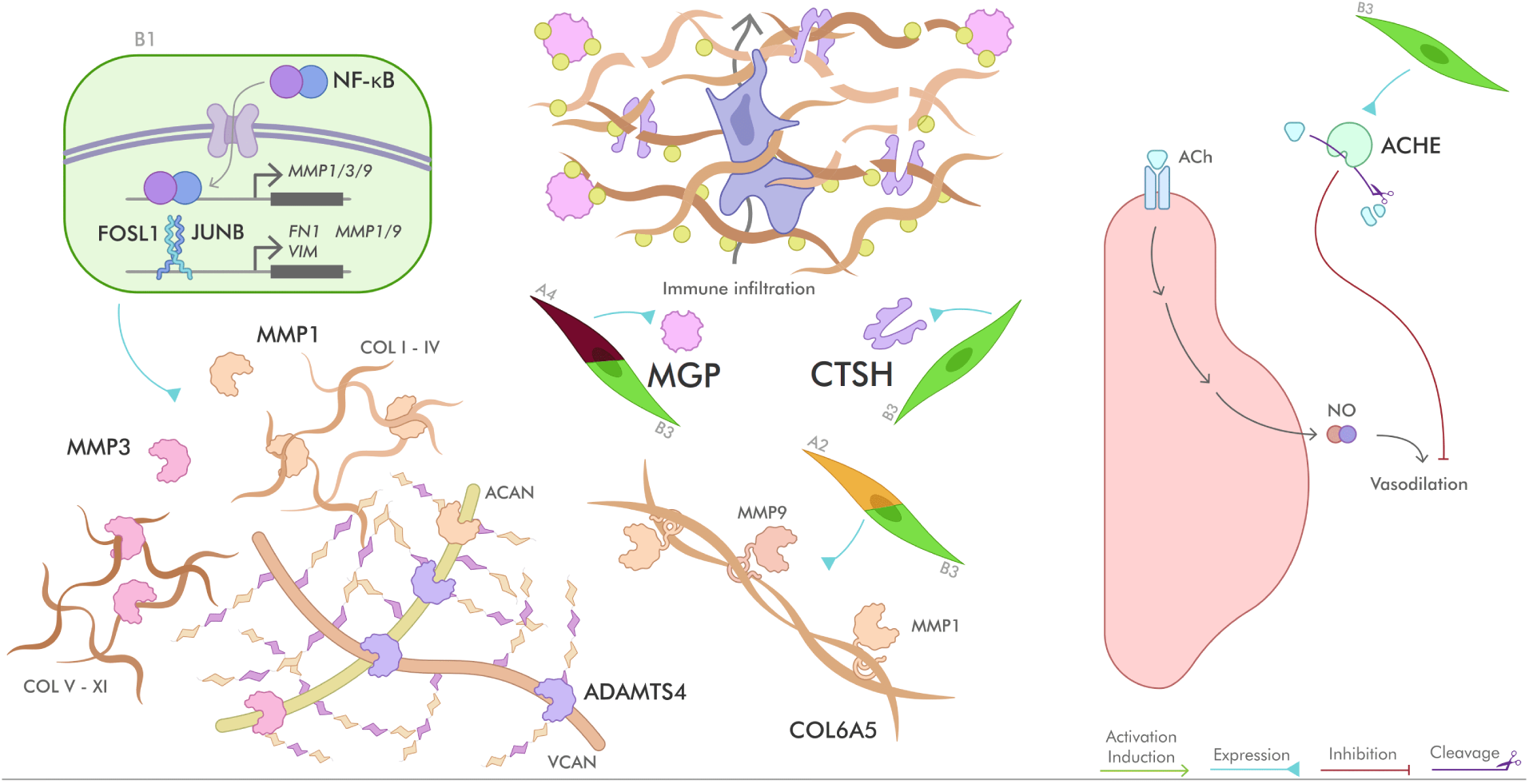
ECM/vasculature regulation mediated by B type fibroblasts. (Left) NF-*κ*B and FOSL1/JUNB mediate the expression of MMPs; (middle) MGP and CTSH, as well as MMP1/9 mediate ECM degradation for immune infiltration; (right) ACHE modulates vasodilation.

**Figure S30.**
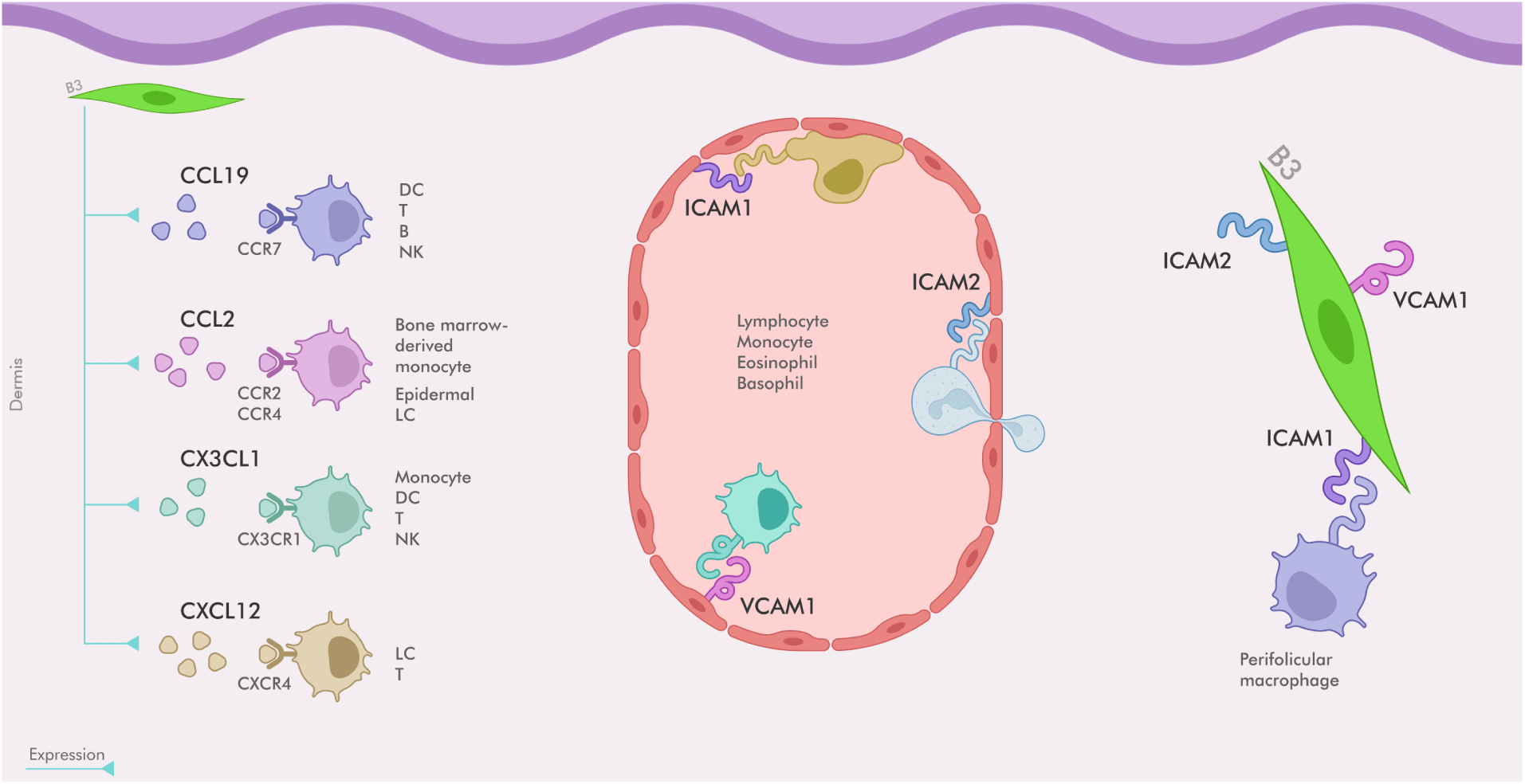
Chemokines and adhesion molecules secreted by B2/B3 fibroblasts. (Left) chemokines secreted and their main cell targets; (middle, right) expressed adhesion molecules (ICAM1/2 and VCAM) and their role in promotion of immune cell adhesion and diapedesis, as well as attraction of macrophages in the dermis.

**Figure S31.**
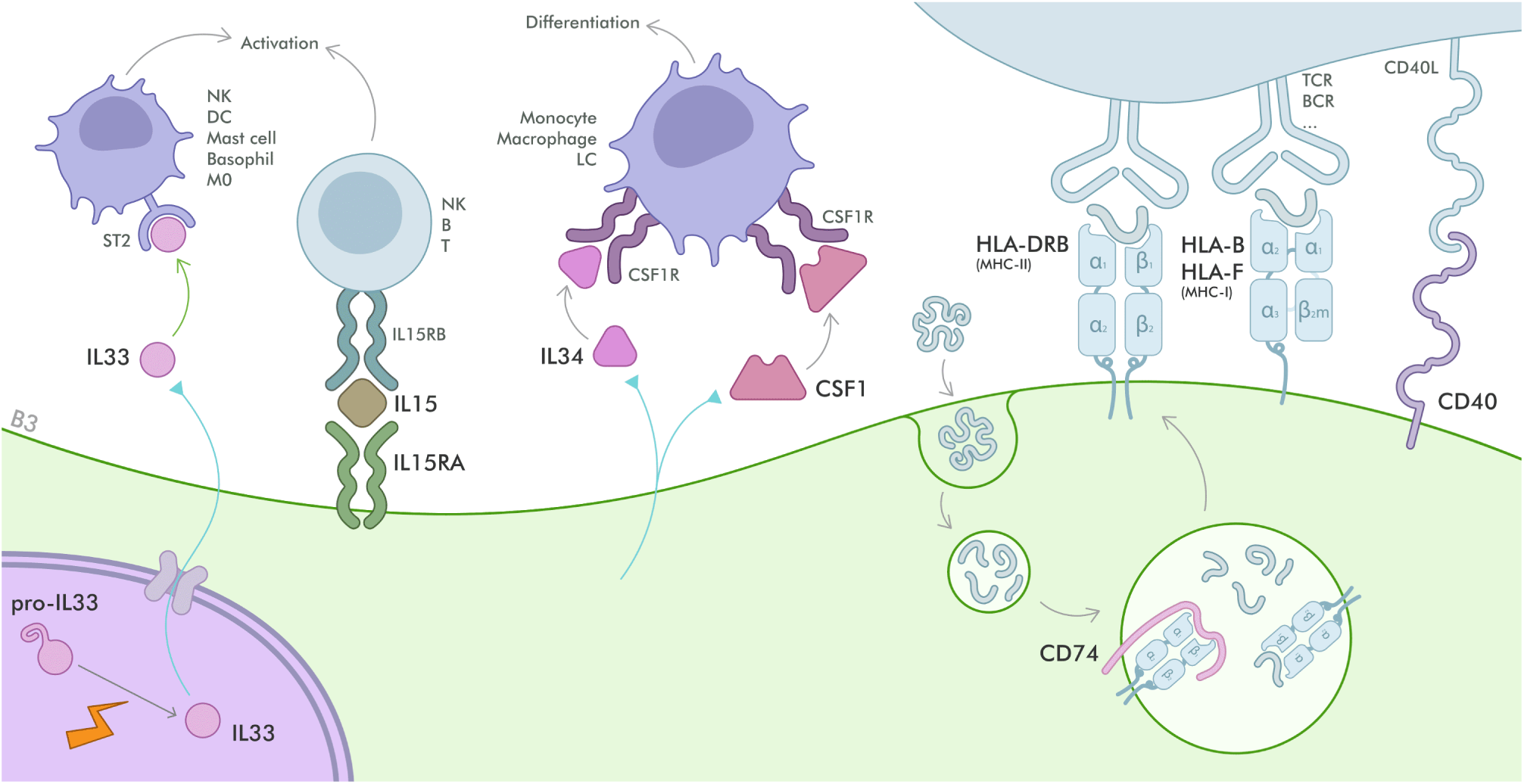
Positive immune modulation by B2/B3 fibroblasts. Mechanisms of modulation including activation of immune cells by IL33, IL15/IL15RA, IL34 and CSF1, as well as antigen processing and presentation by CD74, CD40 and HLA molecules.

**Figure S32.**
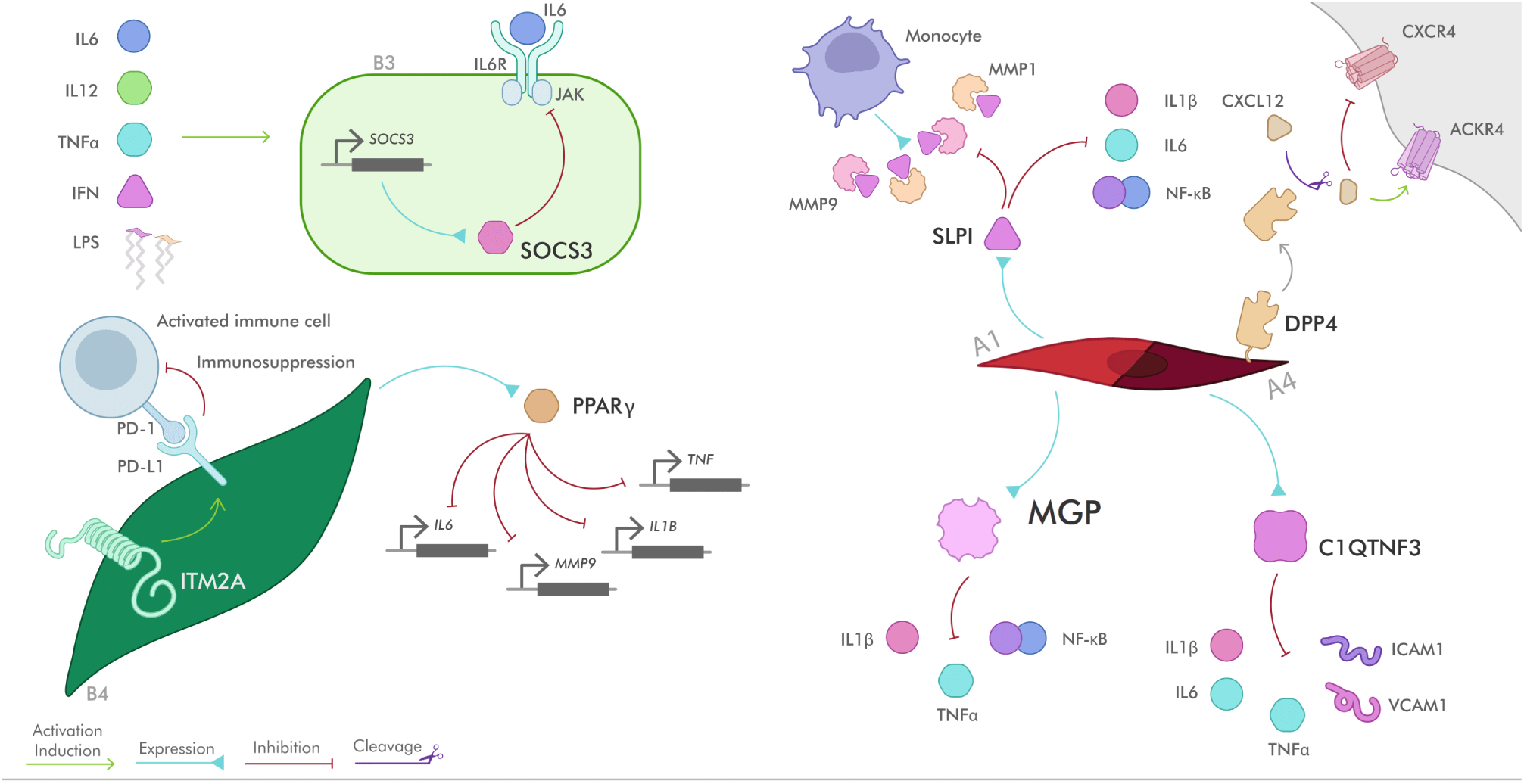
Negative immune modulation by B4 and A fibroblasts. Mechanisms of modulation mediated by SOCS3, ITM2A, PPAR*γ* by B4 fibroblasts, and by SLPI, DPP4, MGP and C1QTNF3.

## Supplementary Tables

**Table S1.** Human dermal fibroblast dataset information.

| Reference | Donors (n) | Age (y) | Sex | Ethnicity | # of fbs. |
| --- | --- | --- | --- | --- | --- |
| (Ahlers et al., 2022) | 3 | 22, 25, 29 | F | Caucasian | 20743 |
| (Billi et al., 2022) | 14 | - | - | - | 5142 |
| (Boothby et al., 2021) | 3 | 64, 66, 67 | M | - | 4915 |
| (Burja et al., 2022) | 2 | 46, 79 | F/M | Caucasian | 2587 |
| (Deng et al., 2021) | 3 | 26-39 | 1F/2M | Han chinese | 5983 |
| (Gao et al., 2021) | 3 | 23, 32, 47 | 1F/2M | - | 2099 |
| (Gaydosik et al., 2019) | 1 | 64 | M | - | 2640 |
| (Gur et al., 2022) | 60 | 28-75 | 49F/11M | - | 15393 |
| (He et al., 2020) | 7 | 38-82 | 4F/3M | - | 3443 |
| (Hughes et al., 2020) | 2 | 58,68 | 1F/1M | - | 450 |
| (Kim et al., 2020a) | 4 | - | 2F/2M | - | 2266 |
| (Liu et al., 2021) | 4 | 26-32 | F | Han chinese | 1270 |
| (Mariottoni et al., 2021) | 1 | 48 | M | Af. American | 894 |
| (Mirizio et al., 2020) | 6 | - | - | - | 1074 |
| (Reynolds et al., 2021) | 5 | 25-60 | F | - | 7327 |
| (Rindler et al., 2021) | 4 | 44-57 | 3F/1M | Caucasian | 2885 |
| (Solé-Boldo et al., 2020) | 2 | 25, 27 | M | - | 2739 |
| (Tabib et al., 2018) | 6 | 23-66 | 3F/3M | White | 2742 |
| (Tabib et al., 2021) | 10 | - | - | - | 4814 |
| (T. Sapiens Cons. and Quake, 2022) | 2 | 33, 59 | M | Hispanic | 2315 |
| (Theocharidis et al., 2020) | 7 | 49-75 | F | - | 3285 |
| (Theocharidis et al., 2022) | 10 | 34-75 | 6F/4M | White | 10441 |
| (Vorstandlechner et al., 2020) | 3 | 30, 36, 43 | F | - | 1161 |
| (Vorstandlechner et al., 2021) | 2 | 36, 43 | F | Caucasian | 1338 |
| (Xu et al., 2021) | 4 | 18-41 | 3F/1M | Han chinese | 494 |

**Table S2.** Mouse dermal fibroblast dataset information.

| Ref | Donors (n) | Age (y) | Sex | Ethnicity | Number of fbs. |
| --- | --- | --- | --- | --- | --- |
| (Abbasi et al., 2020) | 1 | - | - | C57BL/6J | 1718 |
| (Boothby et al., 2021) | 1 | 22 (d) | M | C57BL/6J | 9280 |
| (Buechler et al., 2021) | 1 | 9 (w) | F | C57BL/6J | 6621 |
| (Haensel et al., 2020) | 5 | 7 (w) | F | C57BL/6J | 6327 |
| (Joost et al., 2020) | 10 | 5(6), 9(4) (w) | F | C57BL/6J | 949 |
| (Phan et al., 2020) | 2 | 21 (d) | - | C57BL/6J | 1493 |
| (Shin et al., 2020) | 3 | 2, 12, 18 (m) | F/M | C57BL/6J | 8090 |
| (Shook et al., 2020) | 1 | 7 (w) | M | C57BL/6J | 7728 |
| (Vorstandlechner et al., 2021) | 1 | 7 (w) | F | BALB/c | 4343 |

**Table S3.**
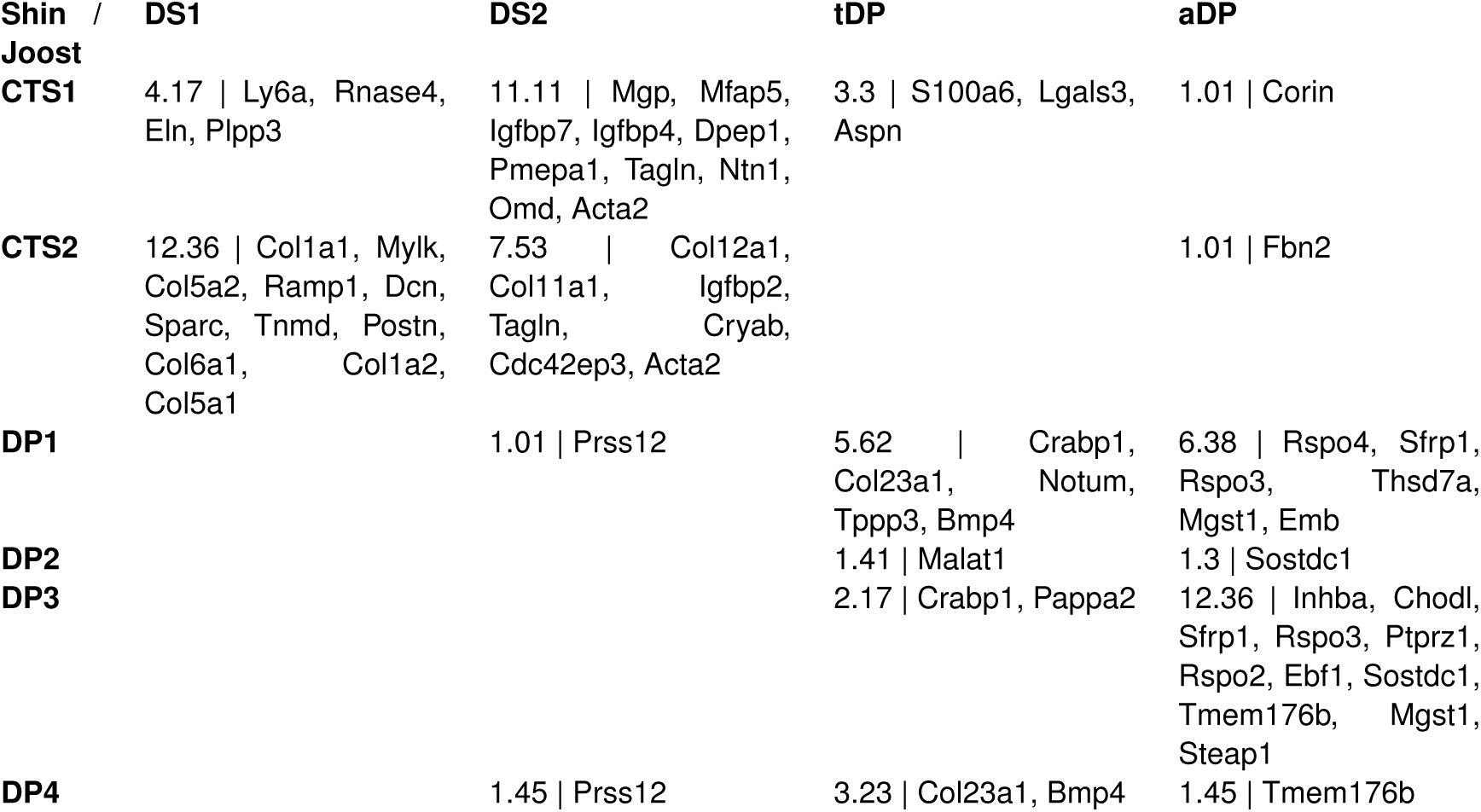
Comparison of Shin 2020 and Joost 2020 DP and DS populations. For each pair of clusters between (Joost et al., 2020) and (Shin et al., 2020), the top 50 genes were selected (based on Supplementary Material from each paper) and the Jaccard index was calculated. The overlapping markers for each compared pair are also shown.

